# Conformation control of the histidine kinase BceS of *Bacillus subtilis* by its cognate ABC-transporter facilitates need-based activation of antibiotic resistance

**DOI:** 10.1101/2020.02.28.969956

**Authors:** Alan Koh, Marjorie J. Gibbon, Marc W. Van der Kamp, Christopher R. Pudney, Susanne Gebhard

## Abstract

Bacteria closely control gene expression to ensure optimal physiological responses to their environment. Such careful gene expression can minimize the fitness cost associated with antibiotic resistance. We previously described a novel regulatory logic in *Bacillus subtilis* enabling the cell to directly monitor its need for detoxification. This cost-effective strategy is achieved via a two-component regulatory system (BceRS) working in a sensory complex with an ABC-transporter (BceAB), together acting as a flux-sensor where signaling is proportional to transport activity. How this is realized at the molecular level has remained unknown. Using experimentation and computation we here show that the histidine kinase is activated by piston-like displacements in the membrane, which are converted to helical rotations in the catalytic core via an intervening HAMP-like domain. Intriguingly, the transporter was not only required for kinase activation, but also to actively maintain the kinase in its inactive state in the absence of antibiotics. Such coupling of kinase activity to that of the transporter ensures the complete control required for transport flux-dependent signaling. Moreover, we show that the transporter likely conserves energy by signaling with sub-maximal sensitivity. These results provide the first mechanistic insights into transport flux-dependent signaling, a unique strategy for energy-efficient decision making.

## INTRODUCTION

Bacteria have exquisite abilities to respond to environmental conditions carefully balancing the benefits of a response against the associated metabolic cost. The importance of this is particularly evident in the context of antibiotic resistance, where one of the main counteracting forces working against the development and spread of resistance is the associated cost in fitness (Andersson and Hughes, 2010; Waglechner and Wright, 2017; Durão *et al.*, 2018). To minimize this cost, many resistance determinants are tightly regulated, which requires that the bacterium is able to accurately determine the severity of an antibiotic attack and adjust gene expression accordingly. We previously described a new sensory strategy in *Bacillus subtilis* that produces an exquisitely fine-tuned response, with three orders of magnitude of output modulation over an input dynamic range of three orders of magnitude (Fritz *et al.*, 2015). This regulatory logic is achieved by a two-component system (BceRS) whose histidine kinase, BceS, works in a sensory complex with an ABC-transporter, BceAB (Dintner *et al.*, 2014; Fritz *et al.*, 2015) (Fig. 1). BceRS controls expression of the *bceAB* operon, and the resulting increased production of the transporter provides resistance against the peptide antibiotic bacitracin (Ohki *et al.*, 2003). A defining characteristic of this system is that the histidine kinase lacks any apparent sensory domains and instead requires the transporter for activation in response to bacitracin (Rietkötter *et al.*, 2008; Hiron *et al.*, 2011).

**Figure 1.**
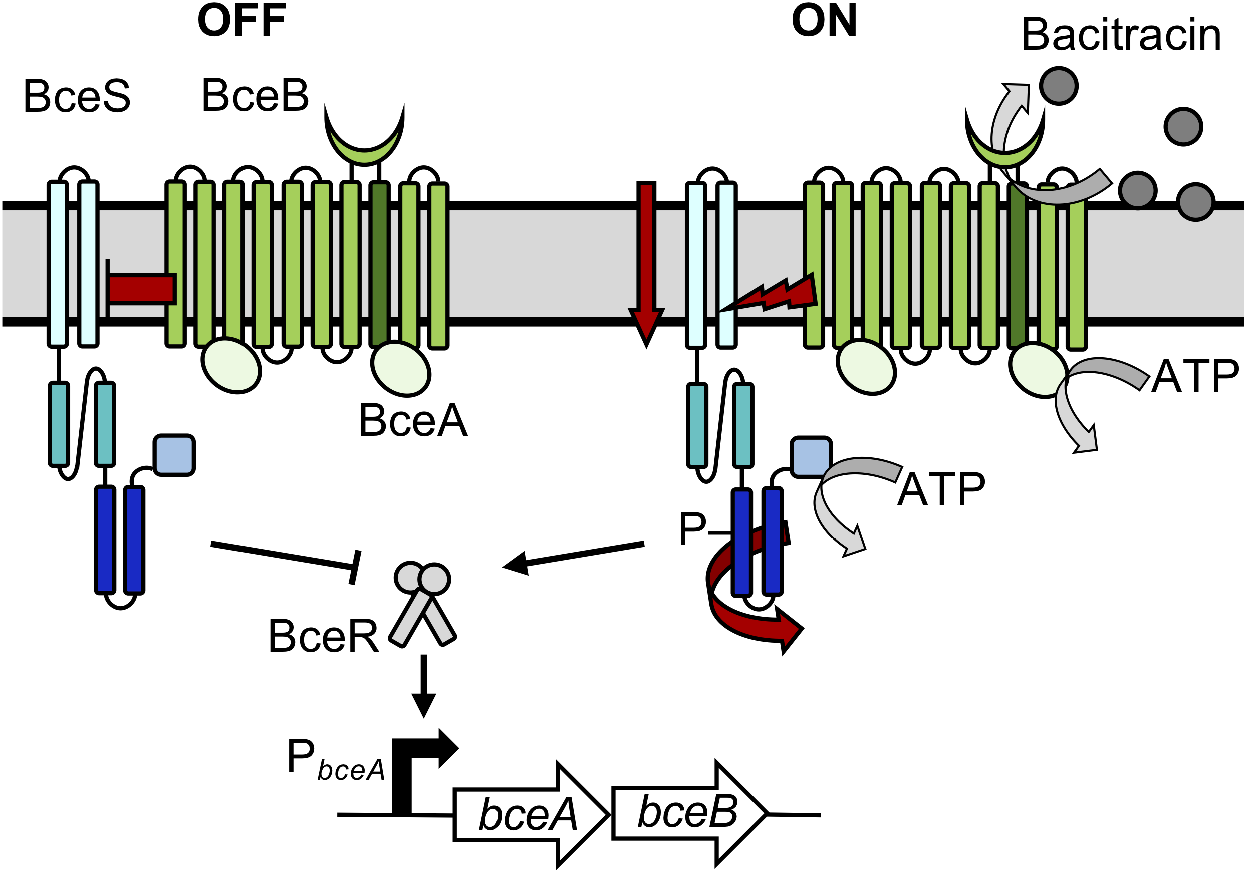
Function of the BceRS-BceAB resistance system of *B. subtilis.* BceS and its domains are shown in shades of blue, with the HAMP-like domain in dark turquoise, and DHp domain in dark blue. Only a single monomer of BceS is shown for clarity. The BceB permease is in bright green with the 8th transmembrane helix marked in dark green and the extracellular domain indicated by a green crescent; two molecules of the ATPase BceA are shown in pale green associated with the BceB monomer, according to the A2B stoichiometry (Dintner *et al.*, 2014). The left section of the schematic shows the system in its inactive (OFF) state, and the repressing function of the transporter indicated by a red blunt arrow. The right shows the active (ON) state in the presence of antibiotics, exemplified by bacitracin, with the activating function of the transporter shown as a red lightning bolt. The motions of kinase domains upon activation are shown as red block arrows and BceS autophosphorylation by a ‘P’. The positioning of the symbols within the sensory complex for repression/activation is arbitrary. ATP hydrolysis reactions and antibiotic removal are indicated by grey arrows; for simplicity ATP hydrolysis is only shown for one ATPase.

Such systems are wide-spread among low G+C content Gram-positive bacteria, including bacilli, lactobacilli, staphylococci, streptococci, enterococci and clostridia (Ouyang *et al.*, 2010; Staroń *et al.*, 2011; Hiron *et al.*, 2011; Dintner *et al.*, 2011; Revilla-Guarinos *et al.*, 2013; Gebhard *et al.*, 2014). They are collectively referred to as Bce-like systems and impart resistance against a broad range of cell wall-active peptide antibiotics (Gebhard and Mascher, 2011; Revilla-Guarinos *et al.*, 2014). Based on computational modelling of the response dynamics in the *B. subtilis* Bce system, we have established that signaling is proportional to transport activity (Fritz *et al.*, 2015). Because the transporter mediates the actual resistance, presumably by removal of the antibiotic from its target in the cell envelope (Gebhard, 2012; Fritz *et al.*, 2015; Greene *et al.*, 2018; Kobras *et al.*, 2020), increased transporter production acts a negative feedback on signaling, which creates the characteristic gradual, fine-tuned response (Fritz *et al.*, 2015). This flux-sensing mechanism effectively allows the bacterium to adjust its response according to its need for additional detoxification capacity, implementing a cost-effective ‘produce-to-demand’ strategy. However, the molecular mechanism of this new signaling strategy, i.e. how the activity of the histidine kinase is controlled within the sensory complex and how this creates the observed analog response behavior, remains unknown.

Histidine kinases typically relay extracellular information to the cytoplasm via autophosphorylation at a conserved histidine residue, which then leads to phosphorylation of a cytoplasmic response regulator that elicits changes in gene expression (Gao and Stock, 2009). Despite their importance in bacterial signaling, we still have limited understanding of the molecular mechanisms by which histidine kinases transmit information through the cytoplasmic membrane. This is in part due to the modular architecture of the kinases, which allows a plethora of different input domains to be coupled to a conserved catalytic core. The core consists of a DHp (also known as HisKA) domain that facilitates dimerization and histidine phosphotransfer, and a C-terminal ATPase domain (Gao and Stock, 2009). The extracellular input and cytoplasmic core domains are separated by transmembrane segments and usually at least one cytoplasmic signal transfer domain (e.g. HAMP or PAS domains) (Gao and Stock, 2009; Gushchin *et al.*, 2017). Recent studies of histidine kinase domains in isolation or together with an adjacent domain suggest that transmembrane signaling can occur via rotating and scissoring motions and piston-like helical displacements of transmembrane helices (Bhate, Molnar, Goulian, and DeGrado, 2015a; Monzel and Unden, 2015; Gushchin *et al.*, 2017; Bhate *et al.*, 2018). The nature of transmembrane motions largely appears to depend on their context with sensory and signal transfer domains, although the individual motions are not mutually exclusive (Gushchin and Gordeliy, 2017; Jacob-Dubuisson *et al.*, 2018).

Additionally, several kinases are controlled by accessory proteins. In many cases, these accessory proteins are transporters (Tetsch and Jung, 2009; Piepenbreier *et al.*, 2017), and most act as repressors of kinase activity, with deletion of the transporter causing constitutive activation of signaling (Hsieh and Wanner, 2010; Witan *et al.*, 2012; Unden *et al.*, 2016). The flux-sensing Bce-like systems are unique in that the histidine kinases fully rely on their cognate transporters for activation, and remain inactive when the transporter is deleted (Rietkötter *et al.*, 2008; Gebhard and Mascher, 2011). Moreover, implementing the reported flux-sensing strategy requires kinase activity to be directly proportional to transport activity (Fritz *et al.*, 2015). To our knowledge, no information is available on how accessory proteins may achieve such positive and precise control over a histidine kinase.

In the present study, we sought to understand the mechanistic basis of histidine kinase activation in the Bce system of *Bacillus subtilis* and how this links to the system’s analog signaling behavior. Through computational predictions, detailed mutagenesis and cysteine cross-linking and accessibility analyses, we provide evidence that the kinase BceS is activated via helical rotations in the DHp domain, accompanied by a piston movement in the second transmembrane domain. Physical motions are most likely relayed via a poorly conserved intervening HAMP-like domain. We also show that the transporter not only acts as the activator of the kinase, but also controls its inactive state. We propose it is this complete control over kinase conformation that couples its activity to that of the transporter. Surprisingly, mutagenesis of the putative signaling domain of the transporter, BceAB, highlighted that the system has not evolved for maximal signaling sensitivity, which likely represents an additional layer of fine-tuning and energy conservation.

## RESULTS

### Generation and validation of a BceS homology model

Analysis of the domain architecture of BceS using SMART (Letunic *et al.*, 2012) showed two N-terminal transmembrane helices, predicted to end at residue F55, and a prototypical cytoplasmic kinase core with a DHp domain from residues E115 to I172 and ATPase domain from S216 to N326. The 60 amino acid linker between the transmembrane and DHp domains contained no predicted domains. To investigate the cytoplasmic composition of BceS in more detail, we used I-TASSER (Zhang, 2008) to predict the tertiary structure of the linker region and DHp domain (residues R56 to N174; Fig. 2a). The closest structural analog was a hybrid protein consisting of the HAMP domain of Af1503^A291V^ from *Archaeoglobus fulgidus* and the DHp domain of *E. coli* EnvZ (PDB 3ZRW; see methods for quality scores), which shares 20% sequence identity and 40% similarity with the BceS fragment used for structure prediction. The homology model of BceS showed the typical DHp domain fold of two antiparallel helices (DHp1 and DHp2), joined by a short loop. In the functional kinase dimer, this will form the four-helix bundle characteristic for most histidine kinases (Gao and Stock, 2009) (Fig.2c). Interestingly, the linker between the membrane interface and DHp domain adopted a HAMP-like conformation with two short parallel helices (HAMP-like 1 and HAMP-like 2), where HAMP-like 2 continues into DHp1 as a single continuous helix (Fig. 2a). The relevance of this domain, which had not been predicted in previous analyses (Mascher, 2006; Dintner *et al.*, 2011), is discussed in more detail below. It should be noted that confidence in the exact conformation of the loops linking the two helices in the HAMP-like domain, and the short loop linking the two DHp domain helices is low in the homology model due to the presence of Ramachandran outliers (see Methods). However, the overall model still provided a plausible illustration of the BceS structure and was used to inform further experimental design.

**Figure 2:**
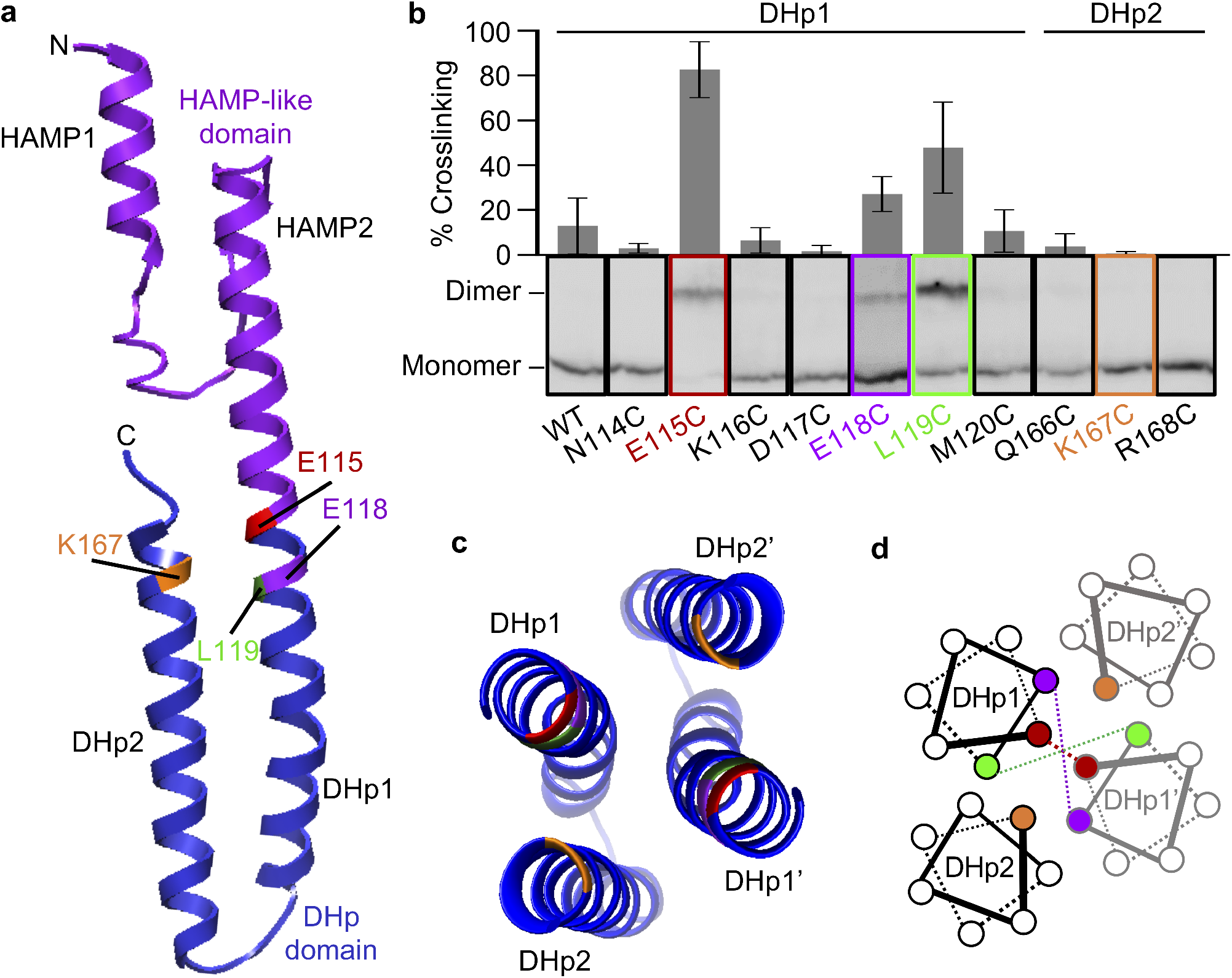
BceS DHp domain forms a four-helix bundle. a, Homology model of the BceS HAMP-like and DHp domains generated by I-TASSER (C-score = −0.76). The HAMP-like domain is shown in purple and the DHp domain in blue. The residues that form Cys-crosslinks in panel b are highlighted, E115 (red), E118 (purple) and L119 (green). Residue K167, which sits opposite E115, is shown in orange. b, Percentage of crosslinking for each BceS Cys variant was calculated from relative band intensities between monomer and dimer within each lane using Fiji (Schindelin et al., 2012). Amino acid substitutions were introduced in parent construct bceSWT-His_8_ (SGB369) and BceS production was controlled by addition of 0.02 % (w/v) xylose. Data are shown as mean ± SD from 2-5 independent repeats. c, d, Schematic diagram of the BceS DHp dimer configuration. Helices labelled DHp1’ and DHp2’ show the second protomer. Left, top down view of the helices in the structural model; right, helical wheel diagram of the top two helical turns in the DHp domain; residues that form crosslinks are highlighted as before.

To test the accuracy of the homology model, we probed the relative orientation of helices within the DHp domain four-helix bundle, based on proximity of individual residues, using oxidative cysteine crosslinking. These experiments were done *in vivo* using cells deleted for the native copy of *bceS* and carrying a single-copy ectopic construct for expression of *bceS* with a C-terminal His_8_-tag, allowing slight over-production of the kinase to facilitate detection by Western blot analysis. BceS contains three native Cys residues, C45 in the second transmembrane helix and C198 and C259 in the ATPase domain. Based on their physical distance from each other and the DHp domain, we did not expect these residues to interfere with our analysis, and indeed no crosslinking product was observed upon oxidation with iodine in wild-type BceS (Fig. 2b). Next, single cysteine substitutions were introduced in the first two helical turns of the helix DHp1 (N114 to M120), and in equivalent positions near the end of the helix DHp2 (Q166 to R168). Disulfide bond formation in these variants should only occur if the substituted residues come into close proximity in the kinase dimer (Pakula and Simon, 1992). The only residues that consistently formed detectable crosslinks were at positions 115, 118 and 119, with the highest degree of crosslinking at position 115 (Fig. 2b). This is in good agreement with their positioning at the core of the four-helix bundle and shows that the structural model closely reflects the conformation of BceS in the cell (Fig. 2c).

To ensure that the introduction of cysteines had not affected the integrity of the protein, we tested the activity of each BceS variant *in vivo.* This was based on their ability to induce expression from the system’s target promoter fused to a reporter gene (P*_bceA_-luxABCDE* or P*_bceA_-lacZ*), following exposure to the antibiotic bacitracin (Fritz *et al.*, 2015). Only the K116C and K167C substitutions led to a complete loss of activity (Fig. S1a). It is therefore possible that the lack of crosslinking observed for the latter, which also faces the core of the DHp domain (Fig. 2c), could be explained by defects in the protein rather than the distance between DHp2 helices in the kinase dimer.

### Charge substitutions in the DHp core suggest a rotation mechanism of kinase activation

Inspection of the DHp domain revealed a striking concentration of charged residues at the membrane-proximal end of the four-helix bundle (Fig. 3). In the wild-type kinase, E115 and K167 create a net neutral charge in the bundle core. As described above and previously (Fritz *et al.*, 2015), cells harbouring wild-type BceS strongly activate expression of the P*_bceA_-lux* reporter in the presence of bacitracin, while in the absence of the antibiotic the activity of the reporter is below the detection limit (Fig. 3a, WT). Interestingly, creation of an ‘all negative’ (K167D substitution) or ‘all positive’ (E115K substitution) DHp core resulted in a strong increase in the basal level of the kinase, while the activity in the presence of the inducer remained unaffected (Fig. 3a). To test whether this effect was indeed due to the changes in core charge and not to substitutions at this position *per se*, we introduced a series of other amino acids into position 115. Exchanging E115 for a different positively charged amino acid, Arg, again resulted in an increase in basal activity, although less pronounced (Fig. S1c). Substitution for Ala or Cys had no detectable effect on activity, showing that the presence of charged amino acids was not in itself important for kinase function. This is consistent with the high density of charged residues in this part of BceS not being conserved among its homologues ((Dintner *et al.*, 2011) and data not shown). Moreover, the constitutive kinase activity could be returned to near wild-type levels by restoring the net neutral charge through combining the E115K and K167D substitutions (Fig. 3a). We therefore concluded that the high constitutive activity in the absence of the inducer was indeed due to the presence of four residues with the same charge in the DHp domain core.

**Figure 3:**
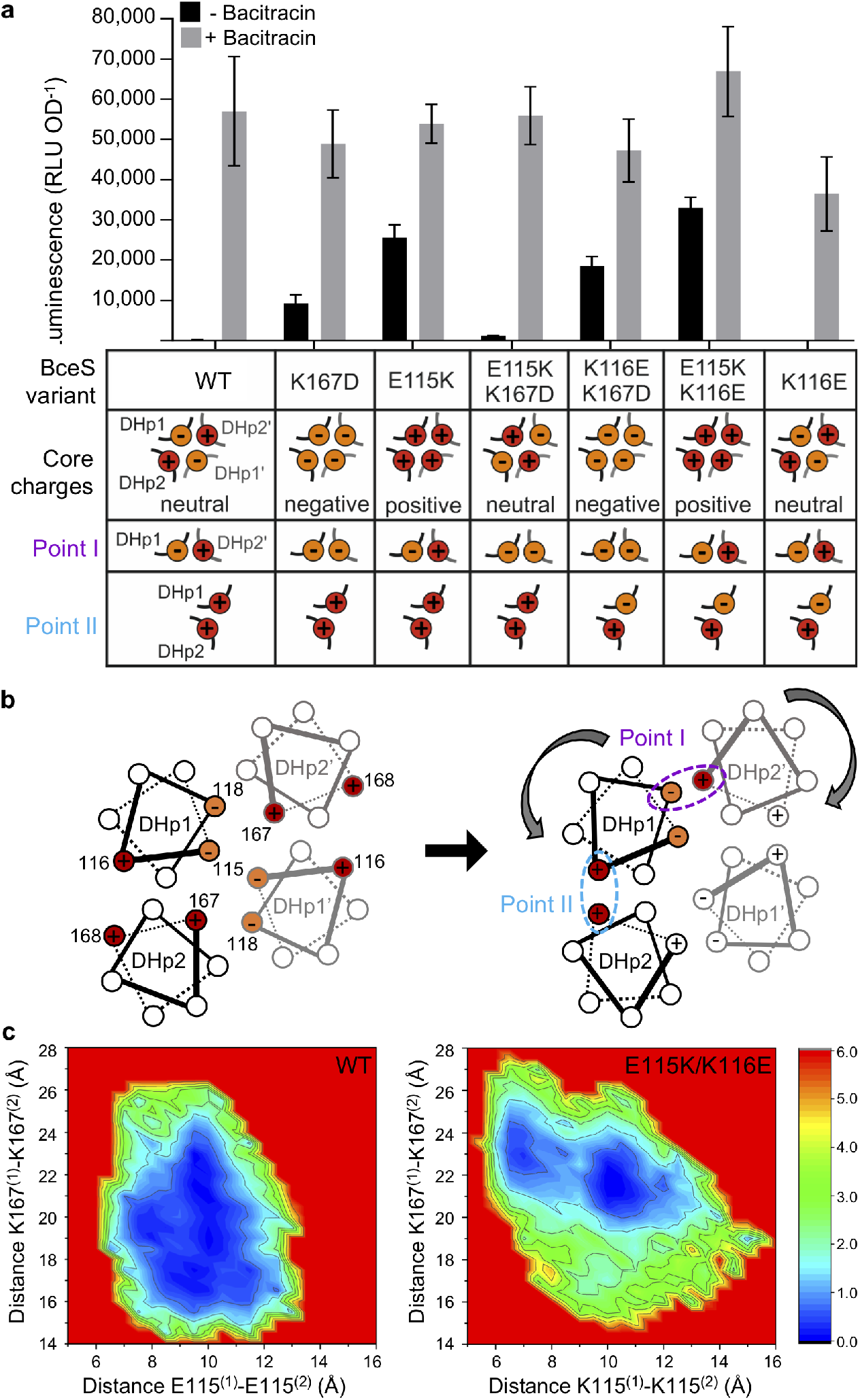
Charges within the BceS DHp domain core affect kinase activity. a, BceS signaling activity in strains harbouring PbceA-luxABCDE and Pxyl-bceS. Amino acid substitutions were introduced in parent construct bceSWT-His8 (SGB369). Signaling was induced with 1μg ml-1 of bacitracin. Data are shown as mean ± SD from 4-22 biological repeats. The schematic below shows the location of individual and net charges in the core and stabilisation points within the four-helix bundle, as illustrated in panel b. Red, positive charge; orange, negative charge. Unpaired t-test comparison to WT showed significant differences (p<0.0001) in the absence of bacitracin for all variants except K116E (p=0.8), and no significant differences (p>0.1) in the presence of bacitracin for all variants, except K116E (p=0.04). Differences with vs without bacitracin were significant (p<0.005) in all strains. b, Helical wheel diagram of the top two helical turns in the DHp dimer configuration showing location of charges and stabilisation points (pink and cyan). Helices labelled DHp1’ and DHp2’ show the second protomer. Grey curved arrows denote the direction of helical rotation. For orientation, the distance between residues at position 115 is likely to be in the range of 4-12 Å based on our cysteine cross-linking results. c, 2D plots of potential of mean force (PMF) obtained in GaMD simulations of the wild type (WT, left) and E115K/K116E variant (right) protein. Intermonomer distances shown are between the Cα carbon atoms of residues at positions 115 (x-axes) and 167 (y-axes). The PMFs are projected onto the surface in kcal mol-1 indicated by the heat scale.

In the current model for histidine kinase activation, the DHp domain undergoes a helical rotation (Gushchin *et al.*, 2017). The strong activating effect we observed upon introduction of four residues of the same charge at the core of the domain made it tempting to speculate that charge repulsion may force BceS to undergo such a rotating motion, placing it in its activated state. According to our homology model, rotation of the helix DHp1 in a counter-clockwise direction and DHp2 in clockwise should create two new points of contact between charged residues: point I between E118 and K167, and point II between K116 and R168 (Fig. 3b). The opposing charges in point I should have a stabilizing effect on the rotated state, and removing this in the K167D variant may explain its lower basal activity compared to E115K (Fig. 3a). In contrast, the proximity of two positive charges in point II should have a destabilizing effect. Replacing one of these with a negative charge (K116E) by itself had no effect on kinase activity. However, combining K116E with the activating effect of the E115K substitution led to a further increase of basal kinase activity compared to E115K alone (Fig. 3a). This effect is entirely consistent with kinase activation through helical rotations in the DHp domain, where we could achieve the highest artificial activation of the protein through creating charge repulsion in the kinase core, while stabilizing the rotated state through opposing charges in the two new points of contact.

Attempts were made to detect this DHp domain rotation *in vivo* using cysteine crosslinking, either following induction with bacitracin or using the constitutively active E115K/K116E variant. No quantitative conclusions could be drawn, however, due to the very large experimental errors (Fig. S2). To provide further evidence of conformational change in the DHp domain, we turned to molecular dynamics (MD) simulations. As discussed above, the homology model (Fig. 2) shows excellent agreement with experimental validation, indicating that it is a reasonable starting point for MD simulations. Given the relatively large putative rotation, we applied Gaussian accelerated MD (GaMD) to more completely sample conformational space (Miao *et al.*, 2015).

To assess conformational differences in the DHp domain between the wild-type kinase and the constitutively active E115K/K116E variant, GaMD simulations were performed on both for a total of 2 μs each (four replicas of 500 ns; Figure S3a). From our simulations, we calculated the potential of mean force (PMF) for a series of internuclear distances for both wild-type and variant residues, to serve as a proxy for the energetic barrier between conformational sub-states. To best illustrate the relative orientation of helices in the DHp domain bundle, we focused on positions 115 and 167, facing the bundle core (Fig. 3). The PMF plots for the internuclear pairs of these residues between monomeric units of the DHp domain are shown in Figure 3c. The lower the PMF value, the more likely the helices will occupy the conformational state defined by the internuclear distances shown. In the wild-type DHp domain, the PMF plot for distances between monomers at amino acids 115 and 167 shows a broad minimum across most of conformational space (Fig. 3c, left). This suggest that there is a relatively high propensity for the domain to explore a diversity of conformational space. Figure 3c (right) shows the same data for the variant DHp, which, instead of the broad minimum, now shows two distinct, more restricted minima. More specifically, the tendency towards a longer K167 distance in the variant would seem consistent with our model for helix rotation (Figure 3b). To more clearly explore the differences between conformational space accessed, we performed principal components analysis (PCA) of the data from Figure 3c. This indicates that there is a significant difference in the principal component space sampled (Fig. S3b), meaning the variant DHp occupies a conformational space that is not accessible to the wild-type domain. Based on the high constitutive activity of the variant, it is plausible to assume that this new conformational space represents the active state of the kinase. Importantly, our simulations show that the wild-type kinase is unable to adopt this conformation on the simulated time scales. This is consistent with our *in vivo* data that shows no detectable kinase activity without activation through the transporter.

### BceS contains a HAMP-like signal transfer domain

Contrary to previous predictions (Mascher, 2006; Dintner *et al.*, 2011), the BceS homology model showed a domain with structural similarity to a HAMP domain between the second transmembrane helix and the DHp domain. HAMP domains adopt a parallel four-helix bundle arrangement with two alternative states of helix packing that transmit a conformational change from an upstream input to a downstream output domain by turning various input movements consistently into helical rotation (Hulko *et al.*, 2006; Lai and Parkinson, 2014; Gushchin *et al.*, 2017). The four-helix bundle is stabilized through multiple packing layers, predominantly mediated by hydrophobic residues (Hulko *et al.*, 2006; Zhou *et al.*, 2011). A sequence alignment of known HAMP domains and the HAMP-like domain of BceS is shown in Fig. 4a, with the positions of the critical packing points highlighted. The hydrophobicity at these positions is largely conserved in BceS, with the exception of two positions occupied by negatively charged Glu instead. Additionally, the conserved Pro residue in helix 1, and Glu in helix 2, typical for HAMP domains, are not present in BceS (Fig. 4a).

**Figure 4:**
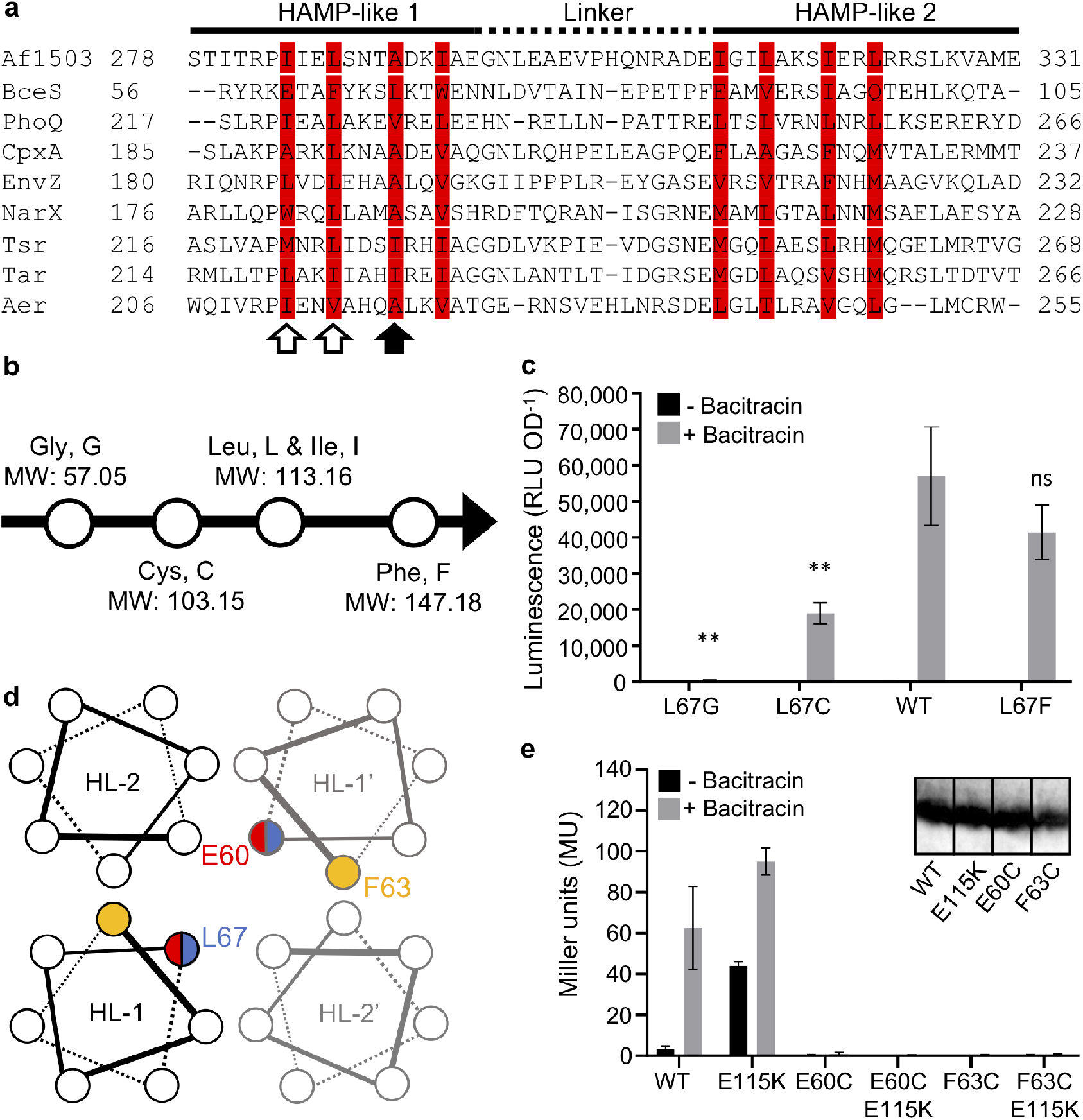
BceS contains a HAMP-like signal transfer domain. a, Sequence alignment of known HAMP domains and the BceS HAMP-like domain generated with ClustalO. Residues in red form the packing layers in the Af1503 HAMP domain. The black solid arrow indicates the packing residue that influences signaling state in some HAMP domains. White open arrows indicate E60 and F63 critical for BceS function. Proteins included and their GenBank identifiers are Af1503 (O28769), BceS (BSU30390), PhoQ (P23837), CpxA (P0AE82), EnvZ (P0AEJ4), NarX (P0AFA2), Tsr (P02942), Tar (P07017), Aer (P50466). b, Schematic showing amino acids and their sizes in MW used in size substitutions. c, BceS signaling activity following L67 amino acid size substitution harbouring PbceA-luxABCDE and Pxyl-bceS. Amino acid substitutions were introduced in the parent construct bceSWT-His_8_ (SGB369). Signaling was induced with 1μg ml-1 of bacitracin. Data are shown as mean ± SD from 3-18 biological repeats. **, significant difference (p<0.01) in activity compared to WT construct, determined by un-paired t-test; ns, not significant; in the absence of bacitracin, no significant differences were detected. d, Helical wheel diagram of the HAMP-like domain. Individual helices are abbreviated HL-1 and HL-2; the labels HL-1’ and HL-2’ show the second protomer. Residues essential for BceS activity are E60 (red) and F63 (yellow); the key packing residue L67 is shown in blue; red/blue semicircles are used to indicate positioning of residues in subsequent helical layers. e, BceS signaling activity of the constitutive ON variant (BceSE115K) combined with loss-of-function substitutions at positions 60 and 63. Amino acid substitutions were introduced in parent construct bceSWT-His_8_ (SGB401). Signaling in cells harbouring PbceA-lacZ and Pxyl-bceS was induced with 1μg ml-1 of bacitracin, BceS production was controlled by addition of 0.02 % (w/v) xylose. Signalling is reported as -galactosidase activity in Miller units, and data are shown as mean ± SD from 3-7 biological repeats. The inset shows an anti-His Western blot analysis to confirm membrane localisation and protein levels of wild-type (WT) BceS and its variants.

Previous investigations of HAMP domains have highlighted one particular residue with key influence on the signaling state, most commonly occupied by small hydrophobic residues that favour bundle packing (Hulko *et al.*, 2006). This residue corresponds to L67 in BceS (Fig. 4a, black arrow). Based on structural work on the HAMP domain of the *A. fulgidus* protein Af1503, this residue (A291 in Af1503) is crucial in controlling the domain’s conformational state. Specifically, it is thought that increasing residue size at this position decreases the threshold for conversion between packing states and can cause increased signal responsiveness or elevated kinase activity, while smaller residues decrease activity (Hulko *et al.*, 2006; Ferris *et al.*, 2011; Ferris *et al.*, 2012) (Matamouros *et al.*, 2015).

To test if the HAMP-like domain in BceS fulfils an equivalent function to *bona fide* HAMP domains, we therefore investigated the role of amino acid size at this position by changing L67 to the smaller Gly and Cys, as well as the larger Phe (Fig. 4b). Indeed, smaller size substitutions reduced BceS activity, up to a near-complete loss of induction in the Gly variant (Fig. 4c). Increasing the size to Phe, however, did not lead to an increase of signaling nor basal activity of BceS. This is consistent with small residues potentially increasing the threshold for conversion between domain conformations (Hulko *et al.*, 2006; Ferris *et al.*, 2011), but we could not find a stimulating effect through a larger residue. We nevertheless conclude that the HAMP-like domain in BceS is likely a functional equivalent to conventional HAMP domains in other proteins contributing to signal transfer within the histidine kinase.

For further functional investigation, we carried out Cys scanning mutagenesis of the HAMP-like domain. The majority of variants maintained at least partial signaling activity (Fig. S4), and in contrast to other studies (Appleman and Stewart, 2003; Matamouros *et al.*, 2015), we did not find any substitutions that led to constitutive activation. The only two substitutions that caused a complete loss of function were E60C and F63C (Fig. 4 and S4). Interestingly, these fall on the first two positions of the packing motif (Fig. 4a, white arrows and 4d), with E60 located one packing layer above the critical L67 discussed earlier, and are therefore likely of structural importance. Both variant proteins are present in the cytoplasmic membrane at the same levels as the wild type protein according to Western blot analysis (Fig. 4e inset), showing that the lack of signaling is not caused by general protein misfolding. To test if the defect of the two variants was due to a loss of signal transfer from either the membrane region of the protein or the transporter, we combined each of the LOF mutations with the E115K substitution that constitutively activates BceS. All resulting variants remained inactive (Fig. 4e), suggesting that the substitutions in the HAMP-like domain likely caused structural disruptions, rather than specifically affecting signal transfer.

### Activation of BceS involves a piston-like displacement in transmembrane helix 2

We next investigated the conformation of the second transmembrane helix in BceS. In many histidine kinases, conformational changes in an extracellular sensory domain are transmitted to the cytoplasmic region through physical movement of the intervening transmembrane helix (Gao and Stock, 2009). In the case of BceS, however, signaling is entirely reliant on the BceAB transport system and therefore does not involve canonical transmembrane information transfer. To investigate if BceS activation nevertheless involved transmembrane helix movements, we carried out a cysteine accessibility scan (Scan-SCAM, (Monzel and Unden, 2015)) of its second transmembrane helix. This analysis requires a cysteine-free background, and we therefore replaced the three native Cys residues mentioned earlier by serine and confirmed that he C45S/C198S/C259S BceS variant was still fully functional (Fig. S5). Individual cysteine replacements were done from positions 51 to 65, surrounding the predicted membrane interface at W54 or F55. The essential residues E60 and F63 were not replaced. All variants were signaling competent (Fig. S5).

The position relative to the membrane was then tested for each Cys residue based on its solvent accessibility, using a modified version of the established *E. coli* protocol (Monzel and Unden, 2015). In brief, treatment of whole cells with N-ethyl-maleimide (NEM) should block any solvent-exposed Cys residue, whereas in untreated cells the sulfhydryl groups should remain available for later reactions. Following isolation of membranes, any unblocked Cys residues are then reacted with methoxypolyethylene glycol maleimide (PEG-mal), resulting in a 10 kDa increase in size, detectable by Western Blotting. For cytoplasmic Cys residues in BceS, we therefore expected to see a shift in size only in samples that had not been treated with NEM (Fig. 5, blue labels). In contrast to the established *E. coli* methodology (Monzel and Unden, 2015), we could not observe PEGylation of membrane-embedded Cys residues in *B. subtilis,* regardless of the presence of SDS, possibly due to differences in membrane composition. We therefore interpreted those residues as being membrane embedded that did not show a size shift in either presence or absence of NEM treatment (Fig. 5, red labels).

**Figure 5:**
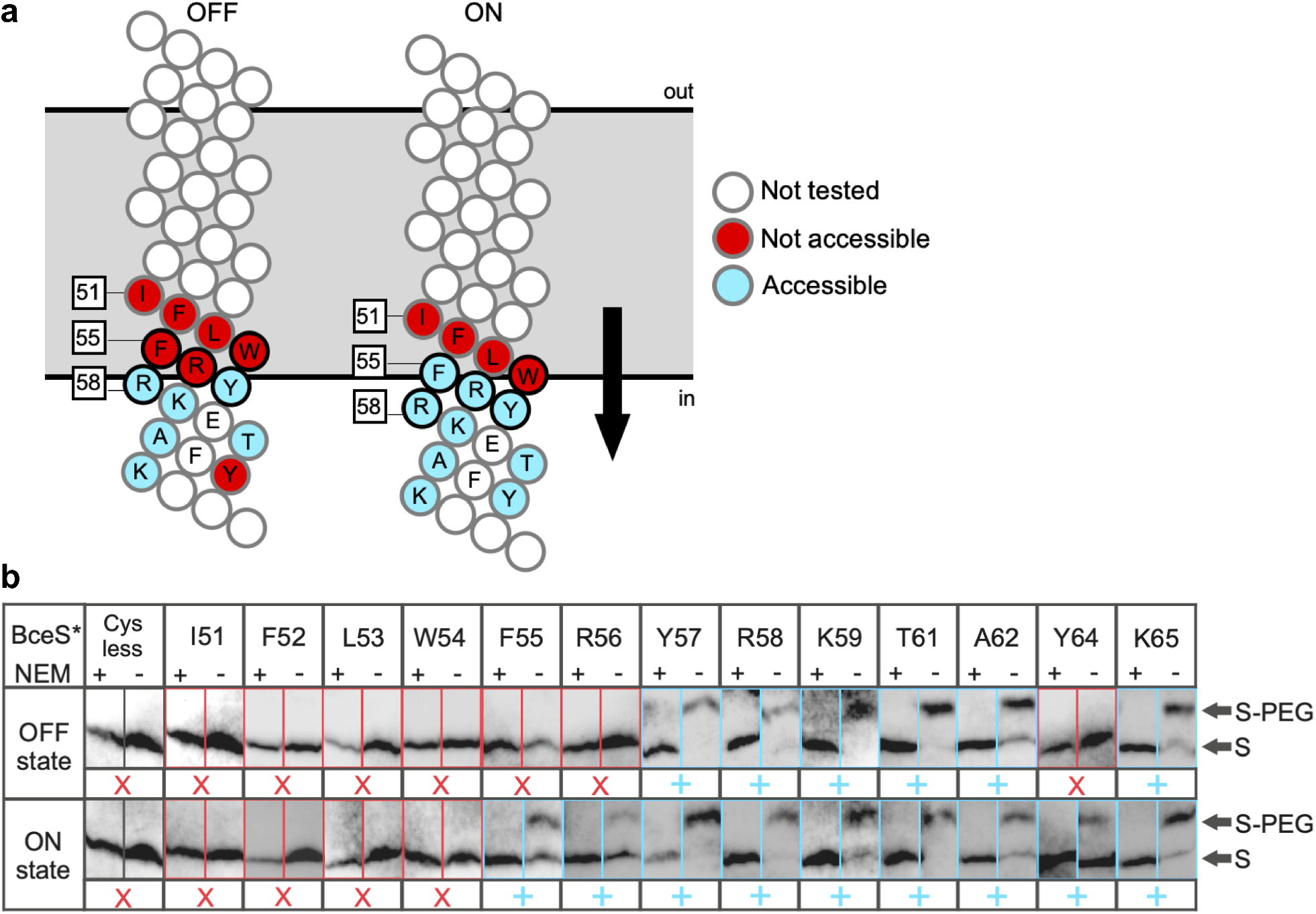
Accessibility of Cys residues at the membrane interface shows a piston-like displacement upon BceS activation. a, Schematic representation of transmembrane helix 2 in the OFF and ON states, respectively. Relevant native amino acids are indicated in one-letter code. Solvent-accessible (PEGylated) Cys replacements are shown in blue, inaccessible (unPEGylated) Cys replacements in red and residues not tested in white. Residues circled in black were tested in triplicate, those circled in grey in duplicate. b, Anti-His Western blot analysis of solvent-accessibility of Cys replacements (Scan-SCAM). Bands corresponding to PEGylated (S-PEG) and unmodified (S) BceS are indicated; individual lanes are outlined in the same colour code as used in panel a. ‘Cys-less’ indicates the parent variant of BceS lacking its native Cys residues. OFF state: amino acids substituted in parent construct bceSC45S/C198S/C259S-His_8_ (SGB936). ON state: amino acids substituted in parent construct bceSE115K/K116E/C45S/C198S/C259S-His_8_ (SGB951).

Our results showed that residues up to position 56 resided in the solvent-inaccessible membrane environment, while residues from Y57 onwards were accessible and therefore cytoplasmic (Fig. 5, ‘OFF state’). We then repeated the same analysis in the background of the E115K/K116E substitutions that place the kinase in a constitutively active conformation. Here, residues from F55 onwards became consistently solvent-accessible (Fig. 5, ‘ON state’), showing that BceS activation involved an inward piston-like displacement by two amino acids of the second transmembrane helix. An interesting observation was residue Y64, which labelled as inaccessible in the OFF state, despite being surrounded by solvent-exposed positions. It is likely that this residue is involved in protein-protein interactions, e.g. within BceS or potentially with BceB, that in the inactive state prevent its labelling through steric effects, as has been proposed before (Monzel and Unden, 2015).

We next sought to test whether BceS could be activated by artificially inducing a shift through introduction of positively charged arginine residues near the membrane interface. This approach has been shown to cause signal-independent activation of the *E. coli* histidine kinase DcuS (Monzel and Unden, 2015). We therefore substituted three of the residues near the cytoplasmic end of transmembrane helix 2 of BceS (W54, F55 and Y57) with Arg, individually and in pairs. (Fig. S6). Variants containing the W54R substitution had approximately 60% higher activities upon stimulation with bacitracin (Fig. S6), which may indicate a stabilisation of or lowered transition threshold to the activated state. However, none of the variants showed a detectable change of basal activity in the presence or absence of the input transporter BceAB, leading us to conclude that Arg substitution alone did not remove the strict reliance on BceAB for signaling.

### The transporter BceAB has complete control over kinase conformation

It was previously shown for several Bce-like sytems that signaling is completely reliant on the transporter (Rietkötter *et al.*, 2008; Ouyang *et al.*, 2010; Staroń *et al.*, 2011; Hiron *et al.*, 2011; Gebhard *et al.*, 2014). We further showed through computational modelling that the signaling output via BceS is directly proportional to the activity of BceAB and does not follow the switch-like on/off behaviour seen in most other systems using transporters as accessory sensing proteins (Fritz *et al.*, 2015; Piepenbreier *et al.*, 2017). This led us to hypothesise that BceAB completely controls the conformation of BceS within the sensory complex and thus couples kinase activity directly to its own transport activity. Our discovery of the DHp domain variants that cause artificial activation of BceS now for the first time enabled us to test this hypothesis experimentally by investigating the role of BceAB in controlling the OFF state of the kinase.

In cells harbouring the wild-type sequence of BceS, the OFF-state activity was below the detection limit of our assays, and deletion of the transporter therefore had no measurable effect (Fig. 6). However, in the E115K or the E115K/K116E variants, basal kinase activity was high, allowing us to test the effect of *bceAB* deletion in the absence of bacitracin.

**Figure 6:**
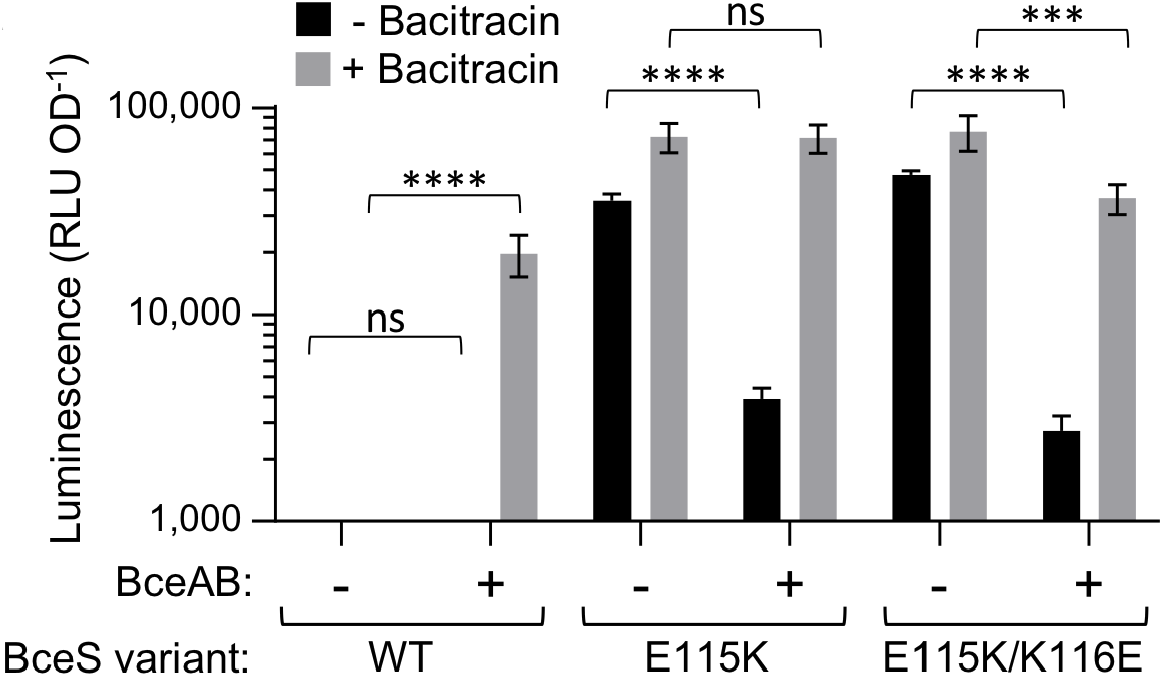
Control of BceS activity by the transporter BceAB. Signaling activity of BceS was measured in strains harbouring PbceA-luxABCDE and challenged with 10μg ml-1 of bacitracin (grey) or left untreated (black). Amino acid substitutions were introduced into parent construct Pxyl-bceS-His_8_, in a strain background carrying a ΔbceS deletion (BceAB +; SGB792), or a ΔbceSAB deletion (BceAB -; SGB818). Data are shown as mean ± SD from 5-12 biological repeats. Results from unpaired t-test are shown by brackets across the strains and conditions compared, analyzing basal and induced activities separately. ****, p<0.0001; ***, p <0.001; ns, p>0.05. Differences with versus without bacitracin were significant (p<0.005) in all strains.

Removing the transporter in this background caused a striking further increase in activity, rendering the kinase fully activated irrespective of the presence of the inducing antibiotic (Fig. 6). Therefore, the inactive transporter can inhibit the activity of the constitutively active DHp domain variants approximately ten-fold. Our data are consistent with a model where inactive BceAB exerts close control over BceS keeping the kinase in its inactive state (Fig. 1, left). This reveals a further layer of kinase control in addition to the transporter’s known activating effect, e.g. preventing spurious signaling in the absence of antibiotics.

### BceAB has not evolved for maximal signaling sensitivity

Recent structural work has provided new insights into the functional mechanism of the Type VII superfamily of ABC transporters, to which BceAB belongs. They act as mechanotransmitters, using cytoplasmic ATP hydrolysis to provide energy for periplasmic processes, such as removal of antibiotics (Crow *et al.*, 2017). In *E. coli* MacB, mechanotransmission is achieved by the transmembrane helix linking the cytoplasmic ATPase interaction site to the periplasmic domain (Crow *et al.*, 2017). In BceAB, the equivalent region is its eighth transmembrane helix (Fig. 1, dark green) (Greene *et al.*, 2018). Based on our earlier discovery that this region of BceB contained a particularly high density of mutations leading to a loss of signaling (Kallenberg *et al.*, 2013), we speculated that this region was crucial for the communication between transporter and kinase. We therefore carried out another Cys scan of residues 533 to 556 in BceB, encompassing approximately half of transmembrane helix 8 and the first alpha-helical segment of the adjacent cytoplasmic region. The native Cys at position 544 was replaced by Ser. Expression of *bceAB* was from a xylose-inducible promoter for consistent transporter production irrespective of signaling activity. The resulting variants, except G535C, displayed comparable bacitracin resistance to the isogenic progenitor strain (Table S2), confirming full functionality of the transporters.

Signaling assays showed that all variants except G543C reached a similar amplitude of reporter induction as strains containing the wild-type construct (Fig. S7). However, we observed some clear difference in the sensitivity of signaling, i.e. the minimal concentration of antibiotic required to elicit a response. In cells with the wild-type construct, the lowest bacitracin concentration to trigger signaling was 3 μg ml^−1^, while a number of variants had a higher threshold of 10 μg ml^−1^ (Fig. 7, green vs pink dots; Fig. S7). The majority of these fell into the C-terminal part of the scanned region, indicating potential defects in coupling to the ATPase. Surprisingly, we found several variants that showed a marked increase in signaling sensitivity and responded already at 0.3 μg ml^−1^ or even 0.03 μg ml^−1^ bacitracin (Fig. 7, dark blue dots; Fig. S7). We had previously seen that a reduction of BceAB transport activity can cause such a shift in signaling threshold, because the negative feedback created by removal of the antibiotic is missing (Fritz *et al.*, 2015). However, as none of these variants showed any change in resistance (Table S2), loss of antibiotic removal cannot explain the increased signaling sensitivity.

**Figure 7:**
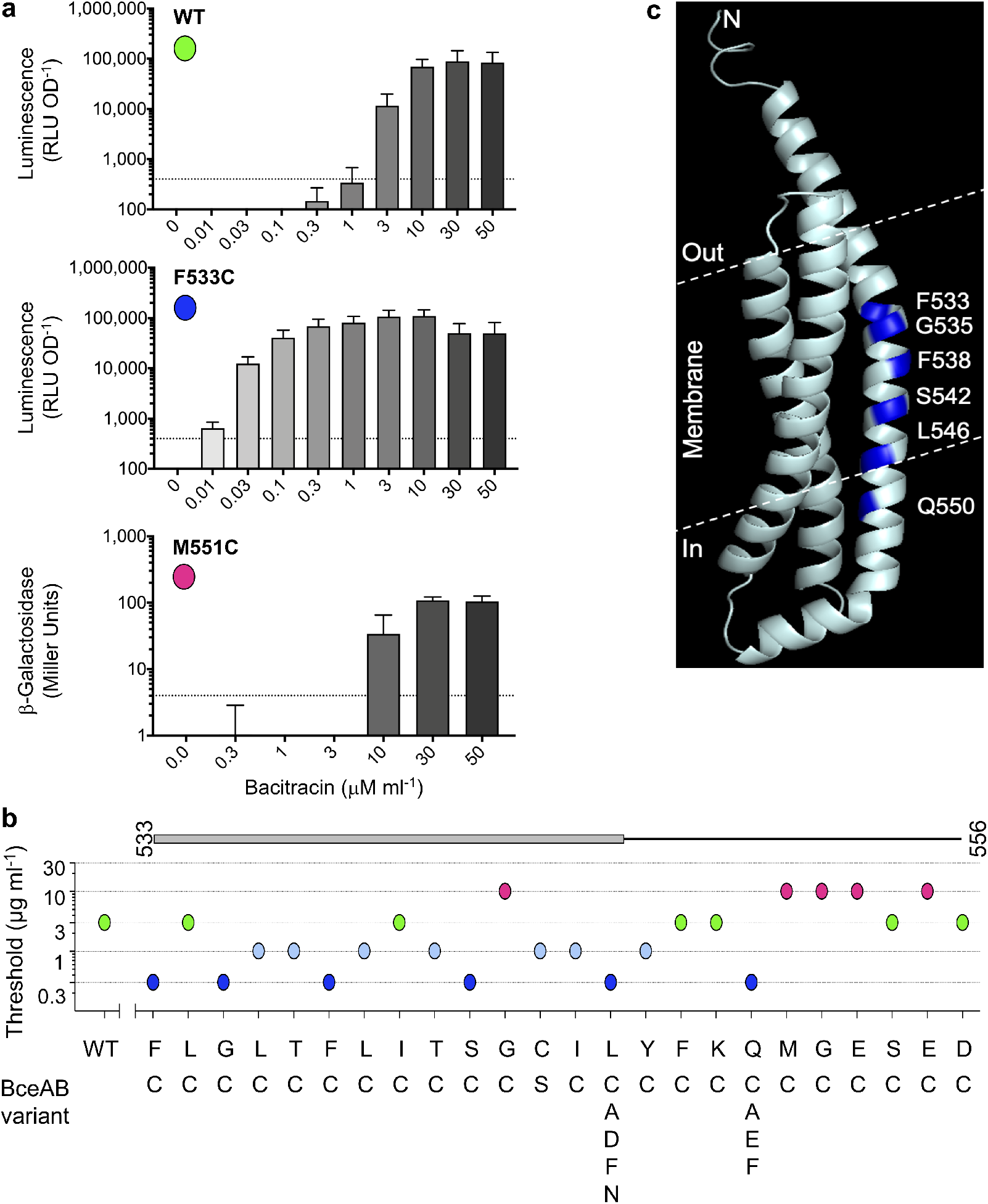
BceAB signals with sub-maximal sensitivity. Amino acid substitutions were introduced into the Pxyl-bceAB-FLAG3 parent construct and signaling assessed using either a PbceA-lacZ (SGB370 and derivatives) or PbceA-luxABCDE (SGB731 and derivatives) reporter. Cells were challenged with a series of bacitracin concentrations for 45 minutes, before signalling activities were determined. a, Signaling phenotype of wild-type (WT) BceAB and representative variants (see Fig. S7 for complete datasets). The colored dots indicate the type of signaling behavior as depicted in panel b (green, WT; blue, hypersensitive signaling; pink, defective signaling). Data are mean ± SD of triplicate experiments. b, Distribution of signaling phenotypes along the transmembrane helix 8 of BceB. The same color symbols are used as in panel a, and the y-axis indicates the threshold bacitracin concentration giving signaling above background. For clarity, threshold concentrations below 0.3 g ml-1 are not resolved. The first line of x-axis labels gives native amino acids, the lines below indicate substitutions tested. The bar above the graph shows positions in the BceB sequence and areas within the membrane (grey bar) or in the cytoplasm (black line). c, Homology model of the C-terminal three transmembrane helices of BceB (from amino acid 507, C-score −0.66) generated using I-TASSER; the top structural analog was MacB of Acinetobacter baumanii (PDB 5WS4). Positions where amino acid substitutions caused increased signaling sensitivity are highlighted in blue. The approximate predicted membrane boundaries are indicated by a white dashed line.

Residue mapping showed that they occurred with a regular periodicity of every four amino acids (Fig. 7b), suggesting positions along the same face of the transmembrane helix. Figure 7c shows a homology model from I-TASSER of the final three transmembrane helices of BceB for illustration. Substitutions with the relatively small amino acid Cys may have enabled tighter helix packing, potentially contributing to more effective signaling. To test this, we replaced two of the positions, L546 and Q550, with a series of other amino acids, including small Ala; charged Glu or Asp; and large Phe. Regardless of the substitution, deviation from the wild-type sequence at these positions consistently led to an increase in signaling sensitivity (Fig. 7b). It is difficult to envisage that all substitutions caused tighter protein interactions. An alternative explanation may be that they are involved in coupling extracellular ligand binding to ATP hydrolysis, and that weakened coupling through the substitutions allowed signaling to occur at lower concentrations. Despite numerous genetic and biochemical attempts, the precise residues with which BceS and BceB interact remain unknown, likely due to an extensive interface between the proteins involving multiple contacts.

While it was beyond the scope of this study to pursue a further functional characterization of the transporter, our findings clearly show that BceB contains a series of residues that prevent signaling from occurring at lower bacitracin concentrations. Given that any replacement we tested led to more sensitive signaling, we therefore concluded that evolution of the transporter has actively avoided such hyper-sensitive signaling, constituting a further layer in minimizing the fitness cost of resistance. Under laboratory conditions, we have not yet been able to detect a competitive disadvantage from harbouring hyper-sensitive variants of BceAB. However, fast growth in carefully controlled conditions is not representative of the complex microbial interactions in natural habitats, where energy-conserving regulatory strategies will likely contribute to the competitive success of a microorganism.

## DISCUSSION

In this study, we set out to investigate the molecular mechanisms of the flux-sensing strategy employed by Bce-like antibiotic resistance systems. Their hallmark feature is that histidine kinase activity is directly proportional to transport activity across a large input dynamic range of antibiotic concentrations (Fritz *et al.*, 2015). This careful adaptation of signaling ensures the resistance transporter is produced precisely to the cell’s demand for detoxification capacity, a very cost-efficient strategy to control gene expression. The results presented here now provide molecular level understanding of how such close control over kinase activity can be achieved.

Analyses of BceS conformation in the inactive and active state showed a piston-like inward displacement of the second transmembrane helix upon kinase activation, associated with helical rotation within the DHp domain as inferred from mutagenesis analysis and GaMD simulations. Tertiary structure predictions and experimental validation indicate that BceS possesses an intervening HAMP-like domain that likely fulfils a signal transfer function.

While bacterial transmembrane histidine kinases have been studied for decades and we have a very good understanding of the function of the individual domains (reviewed in Gao and Stock, 2009; Jacob-Dubuisson *et al.*, 2018)), a complete understanding of how extracellular events (e.g. ligand binding) are transmitted to the cytoplasmic catalytic domains to trigger autophosphorylation is only beginning to emerge (Gushchin and Gordeliy, 2017; Jacob-Dubuisson *et al.*, 2018). In a canonical transmembrane histidine kinase, extracellular ligand binding causes conformational changes in the sensory domain, which will directly influence the transmembrane segments. Due to the diversity of sensory domains, there is also a diversity in the conformational changes received by the transmembrane helices. In the nitrate sensor NarQ of *E. coli,* for example, nitrate binding by the sensory domain causes a complex combination of a helical break between sensory and transmembrane domains with scissoring and twisting motions at the periplasmic end of the membrane helices. The net result of this is a piston-shift displacement perpendicular to the membrane, which in turn elicits conformational changes in the adjacent cytoplasmic HAMP domain (Gushchin *et al.*, 2017). In the *E. coli* kinases DcuS and CitA, binding of fumarate or citrate, respectively, by an extracellular PAS domain causes compaction of the sensory domain, which displaces the adjacent transmembrane helix towards the periplasm, again resulting in a piston-shift (Monzel and Unden, 2015; Salvi *et al.*, 2017). A piston-shift in the opposite direction, i.e. towards the cytoplasm, is seen in activation of *E. coli* chemoreceptors like Tar (reviewed in Parkinson *et al.*, 2015).

Alternative motions in transmembrane signalling are diagonal scissoring displacements, for example in the antimicrobial peptide and cation sensor PhoQ (Molnar *et al.*, 2014), or helical rotations in kinases such as the quorum sensor AgrC of *Staphylococcus aureus* (Wang *et al.*, 2014). Due to the diversity of conformational changes, there is currently no consensus model for transmembrane signaling by histidine kinase, and it is likely that the different motions can work together within a single protein (Gushchin and Gordeliy, 2017; Jacob-Dubuisson *et al.*, 2018). Our data for BceS suggest that despite its unusual input mechanism via the BceAB transporter, transmembrane signaling is consistent with existing models of kinase activation and occurs via an inward piston displacement. It is of course possible that further structural changes of higher complexity occur in BceS, or indeed within BceS-BceB sensory complex. The unusual behavior of residue Y64 in the Scan-SCAM analyses may provide a first glimpse of such, but this is not currently resolved by our experimental data.

To complete signaling and kinase activation, the transmembrane motions will have to be translated into changes within the conserved kinase core to cause autophosphorylation, the common theme for which is helical rotation (Gushchin and Gordeliy, 2017; Jacob-Dubuisson *et al.*, 2018). In kinases where transmembrane signaling already occurs via rotation, the conformational change can likely be directly propagated to the core (Gushchin and Gordeliy, 2017). Kinases that signal via piston-displacements, however, require a means of converting this motion into a helical rotation. Recent crystallographic evidence has provided detailed molecular insights into how HAMP domains act as such converters of conformational change (Gushchin *et al.*, 2017). While BceS lacks a *bona fide* HAMP domain, our results are in good agreement with such a mode of signaling, where the piston-shift of transmembrane helix 2 is translated into helical rotations within the DHp domain via the intervening HAMP-like domain.

An earlier NMR study reported on the structure of the BceS-like kinase NsaS from *S. aureus,* which possesses a much shorter linker region between its transmembrane and DHp domains (Bhate, Molnar, Goulian, and DeGrado, 2015b). This linker region adopts a coiled-coil conformation instead of the HAMP-like fold of BceS. Inspection of kinase sequences in an existing collection of Bce-like resistance systems (Dintner *et al.*, 2011) showed that the vast majority of these proteins contained the larger version of the linker (approximately 60 amino acids), and only a small number possessed short (around 20 amino acid) linkers with a predicted coiled-coil motif. BceS therefore appears to be the more typical subtype of this group of kinases than NsaS. Interestingly however, akin to the alternative helix packing mechanism proposed for HAMP domains (Hulko *et al.*, 2006), activation of NsaS was also proposed to involve switching between helical packing states of the coiled-coil (Bhate, Molnar, Goulian, and DeGrado, 2015b). The molecular principles of activation may therefore be conserved between both subtypes of BceS-like kinases.

To further investigate the activation mechanism of BceS, we sought to force each of the conformational changes via amino acid substitutions. Charge repulsion at the core of the DHp domain to trigger helical rotation consistently led to kinase activation without a physiological stimulus (Fig. 3). However, we were unable to artificially trigger activation through substitutions that should have induced piston displacements in the membrane (Fig. S6) or induced the active conformation of the HAMP domain (Fig. 4 and S4). This is in contrast to studies of other kinases where such substitutions least partially raises the basal activity (Appleman and Stewart, 2003; Matamouros *et al.*, 2015; Monzel and Unden, 2015). In canonical histidine kinases, the input signal is detected directly, e.g. through ligand binding, and it might be speculated that this might be relatively easy to mimic through amino acid substitutions. In contrast, BceS-like proteins are subject to control by the transporter. It is conceivable that in such a kinase small changes in individual domains are insufficient for artificial activation, unless the DHp domain is directly manipulated. This may be supported by the MD simulations, which showed that the wild-type DHp domain does not by itself sample the conformational space observed in the active variant (Fig. S3), implying the transporter may be required to provide the necessary force for transition to the active state.

Transporters are commonly used for accessory signaling control, but BceAB is unusual in that it is an activator of signaling, while most others act as repressors (Piepenbreier *et al.*, 2017). We previously showed that transporter and kinase constitutively form a signaling complex in the membrane, irrespective of the inducer, bacitracin (Dintner *et al.*, 2014). To activate the kinase proportionally to transport activity, the transporter must therefore be able to directly influence kinase activity within this complex. When we first tested this *in vitro,* we noted that BceAB reduced kinase autophosphorylation in the OFF state, but could not explain the observation at the time (Dintner *et al.*, 2014). Here, we now showed that the transporter indeed not only controls the ON but also the OFF state of BceS (Fig. 6). We propose that through controlling both states of the kinase, the transporter is able to exert the close control needed to explain how signaling is maintained in proportion to transport activity over more than three orders of magnitude in antibiotic concentrations (Fritz *et al.*, 2015). A crucial feature of the flux-sensing mechanism is an immediate reduction of signaling if transport activity is low, either because the antibiotic threat is over or because of sufficient detoxification capacity (Fritz *et al.*, 2015). The active switching off of BceS by BceAB described here explains how this is realized on the molecular level. Complete control also allows the transporter to actively prevent spurious signaling by the two-component system. Finally, our mutagenesis analysis of BceAB revealed that evolution of the transporter itself appears to have actively avoided signaling at very low antibiotic concentrations, highlighting yet another level of cost reduction.

Taken together, we here present a molecular level model for activation of antibiotic resistance based directly on the cell’s need for protection, using a flux-sensing signaling strategy that employs multiple layers of active energy conservation to minimize fitness costs of resistance. The system is centered on two-component signaling, one of the most important regulatory features of bacteria. Our findings should therefore have wide relevance and serve as a model for understanding how fine-tuned gene expression in bacteria can achieve an optimal cost-benefit balance.

## EXPERIMENTAL PROCEDURES

### Bacterial strains and growth conditions

All strains used in this study are listed in table S1 and were routinely grown in Lysogeny Broth (LB; 10 g l^−1^ tryptone, 5 g l^−1^ yeast extract, 5 g l^−1^ NaCl) at 37 °C with aeration. Solid media contained 15 g l^−1^ agar. Selective media contained chloramphenicol (5 μg ml^−1^), macrolide-lincosamide-streptogramin B (erythromycin; 1 μg ml^−1^, lincomycin; 25 μg ml^−1^), spectinomycin (100 μg ml^−1^), kanamycin (10 μg ml^−1^), or ampicillin (100 μg ml^−1^).

### DNA manipulation and strain construction

All primers and plasmids used in this study are given in table S1. Site-directed mutagenesis was carried out according to the manufacturer’s instructions for the QuikChange II site-directed mutagenesis kit (Agilent Technologies), or by PCR-overlap extension (Ho *et al.*, 1989). *Bacillus subtilis* transformations were carried out via natural competence as previously described (Harwood and Cutting, 1990). DNA for transformation was either provided as linearised plasmid DNA, or as genomic DNA from a *B. subtilis* strain already carrying the desired genetic construct.

To allow manipulation of both *bceS* and *bceAB* sequences, an unmarked deletion strain lacking all three genes *(ΔbceSAB,* SGB377) was constructed as described previously (Arnaud *et al.*, 2004), employing plasmid pAK102. Strains lacking *bceS* only *(ΔbceS;* TMB1036) or *bceAB* only (*bceAB*::kan; TMB035 (Rietkötter *et al.*, 2008)) were available from previous work. Signaling activity was assessed in strains harbouring a chromosomally integrated transcriptional reporter construct of the target promoter P*_bceA_*, fused to either *luxABCDE* (pSDlux101 (Kallenberg *et al.*, 2013)) or *lacZ* (pER603 (Rietkötter *et al.*, 2008)). Details of the assays used are given below.

Genetic manipulation of *bceS* was performed using previously constructed plasmids as template. For Western blot detection, the construct pSD2E01 encoding BceS with a C-terminal octa-histidine tag (P*_xyl_-bceS*^WT^*-*His_8_ (Dintner *et al.*, 2014)) was employed. Un-tagged constructs were derived from pAS719 (P_xyl_*-bceS*^WT^). Genetic manipulation of *bceAB* was performed using the previously constructed plasmid pFK727, encoding BceAB with a C-terminal triple FLAG-tag on the permease (P*_xyl_*-*bceAB*-FLAG3 (Kallenberg *et al.*, 2013)). All constructs used in this study integrate stably and as a single copy into the *B. subtilis* W168 chromosome.

### Signaling activity assays

In strains carrying the P*_bceA_-lacZ* reporter, signaling was assessed following challenge of exponentially growing cells with Zn^2+^-bacitracin (bacitracin) and measured as β-galactosidase activity (expressed in Miller Units), as described previously (Rietkötter *et al.*, 2008). For strains carrying the P*_bceA_-luxABCDE* reporter, luciferase activities were assayed using a Tecan® Spark® microplate reader controlled by SparkControl™ V2.1 software. Starter cultures were grown at 37 °C in 2 ml of LB medium overnight, then diluted 1:1000 into fresh medium before adding 100 μl into 96-well plates (black, clear flat bottoms; Corning®). Samples were grown at 37 °C with orbital agitation at 180 rpm and allowed to achieve early exponential growth (OD_600_ 0.3 – 0.4), before adding Zn^2+^-bacitracin to a final concentration of 1 μg ml^−1^ unless otherwise indicated. OD600 and relative luminescence units (RLU; integration time 1 s) were measured every 5 min and were background-corrected using values from LB-only wells. RLU was normalised for cell density (RLU OD^−1^). The reported values are steady-state activities achieved 60-75 min after bacitracin addition.

Cells were grown without the addition of xylose, as the basal activity of the promoter P*_xylA_* (Radeck *et al.*, 2013), used to drive expression of the ectopic copies of *bceS* and/or *bceAB,* was sufficient to achieve full signaling activity. In strains containing the native copy of *bceAB (ΔbceS* background) signaling was induced with 10 μg ml^−1^ of Zn^2+^-bacitracin. In strains containing only the ectopic construct for BceAB production *(ΔbceSAB* background), signaling was induced with 1 μg ml^−1^ of Zn^2+^-bacitracin. These concentrations were established in our earlier work as eliciting maximum signaling activity in the respective genetic backgrounds (Fritz *et al.*, 2015).

### Bacitracin sensitivity assays

The sensitivity of *B. subtilis* strains to bacitracin was determined as described (Kallenberg *et al.*, 2013), with minor modifications. Serial two-fold dilutions of Zn^2+^-bacitracin from 32 μg ml^−1^ to 0.5 μg ml^−1^ were prepared in Mueller-Hinton medium containing 0.2 % (w/v) xylose to ensure maximal production of BceAB. For each concentration, 100 μl volumes in 96-well plates were inoculated 1:400 from overnight cultures grown in Mueller-Hinton medium with xylose and selective antibiotics. Each culture was scored for growth after 20 to 24 h of incubation at 37°C with aeration. The MIC was determined as the lowest bacitracin concentration where no visible growth was detected.

### Cysteine cross-linking *in vivo*

Cysteine cross-linking was assessed in strains carrying the *ΔbceSAB* deletion complemented with xylose-inducible ectopic copies of *bceAB*-FLAG_3_ and *bceS*-His_8_ with single-cysteine substitutions and the P*_bceA_-lux* reporter (SGB369 and derivatives). Starter cultures were grown in 2 ml of LB medium overnight at 37 °C with agitation, then diluted 1:500 into 55 ml of fresh medium supplemented with 0.02% (w/v) xylose. In mid-exponential growth (OD_600_ 0.5 – 0.7), oxidative cross-linking of cysteine residues was triggered by addition of 250 μM of iodine for 15 min under continued incubation. Where required, cells were induced with 60 μg ml^−1^ Zn^2+^-bacitracin for 45 min prior to cross-linking. Cross-linking reactions were stopped with 20 mM N-ethylmaleimide (NEM) for 15 min under continued incubation. Cells were harvested by centrifugation (3,000 × *g,* 20 min, 4 °C) and resuspended in 5 ml of ice cold lysis buffer (100 mM Tris-HCl [pH 8.0], 150 mM NaCl, 10% (w/v) glycerol, 20 mM NEM). Cell disruption was achieved by at least three cycles of 15s sonication on ice with 15s rest. Debris was removed by centrifugation (3,000 × *g*, 20 min, 4 °C), and membranes were collected from supernatants by ultracentrifugation (150,000 × *g*, 1 h, 4 °C). Membranes were resuspended in 50 μl lysis buffer, and approximately 150 mg protein per sample was mixed with non-reducing sample buffer (1× final concentration; 1.5 M Tris/HCl [pH 6.8], 30 % (w/v) glycerol, 350 mM SDS, 0.12 mg ml^−1^ bromophenol blue) for SDS-PAGE and Western blot analysis (see below). For quantification of band intensities Fiji (Schindelin *et al.*, 2012) was used to obtain a profile plot of each lane, and the relative intensity of each band (monomer and dimer BceS) within the lane was expressed as a percentage of the total band intensities.

### Solvent-accessibility of Cysteine residues *in vivo* (Scan-SCAM)

For solvent-accessibility tests in the second transmembrane helix of BceS, the *ΔbceSAB* deletion strain, complemented with xylose-inducible ectopic copies of *bceAB*-FLAG3 and *bceS*-His_8_ and the P*_bceA_-lux* reporter was employed. The *bceS* complementation construct was made cysteine-free via C45S/C198S/C259S substitutions, followed by introduction of single cysteines to be tested (strain SGB936 and derivatives). The same was done in the E115K/K116E variant to assess transmembrane topology in the active state (strain SGB951 and derivatives). Cultures were grown to mid-exponential phase as described above. Cells were harvested, washed twice with 50 mM HEPES buffer (pH 6.8) and resuspended in 5 ml of the same buffer. Water exposed (i.e. cytoplasmic) cysteine residues were masked with 40 mM N-Ethylmaleimide (NEM) for 1.5 h at room temperature with shaking or left untreated. Masking reactions were stopped by three washes in cold (4 °C) 50 mM HEPES buffer (pH 6.8). Cells were resuspended in 5 ml of ice cold lysis buffer, sonicated and membranes collected as above. Membranes were resuspended in 40 μl accessibility buffer (50 mM HEPES [pH 6.8], 1.3% (w/v) SDS). Cysteines that had not been masked with NEM in the first step were labelled with 5 mM methoxypolyethylene glycol maleimide (PEG-mal; Sigma-Aldrich) for 1.5 h at room temperature with agitation. The reaction was stopped with 1× LDS sample buffer (Expedeon) and 250 mM DTT and rested for 5 min. PEG-mal accessible and non-accessible BceS were analyzed by Western Blot.

### Western blot analysis

Protein samples (~150 μg) from *in vivo* cross-linking and Scan-SCAM were subjected to SDS-PAGE on RunBlue™ TEO-Tricine SDS mini gel in either Tris-Tricine-SDS (Sigma-Aldrich) or Teo-Tricine-SDS (Expedeon) buffer, run for 50 min at 120V. Samples were transferred to polyvinylidene fluoride (PVDF) membranes (Merck Millipore, activated in 100 % (v/v) methanol) in 1 × RunBlue SDS Running Buffer (Expedeon), 20 % (v/v) methanol at 75 mA for 70 minutes using a PerfectBlue™ Sedec™ Semi-Dry Electroblotter (Peqlab). Membranes were blocked with 7.5 % (w/v) skimmed milk powder in PBST (137 mM NaCl, 2.7 mM KCl, 10 mM Na_2_HPO_4_, 1.8 mM KH_2_PO_4_ [pH7.4], 0.1 % (v/v) Tween 20), washed three times in PBST, and incubated with primary antibody (Penta·His Antibody, QIAGEN®, 1:2000 dilution) in 1% bovine serum albumin in PBST for 2h at room temperature. Following three washes in PBST, incubation with secondary antibody (Peroxidase-AffiniPure Goat Anti-Rabbit IgG, Jackson ImmunoResearch Europe Ltd, 1:4000 dilution) was carried out in milk-PBST for 1h at room temperature, followed by three final washes in PBST. Detection of proteins was by chemiluminescence using Pierce™ ECL Western Blotting Substrate (Thermo Fisher Scientific Inc.) and a Qiagen CCD camera with Fusion software.

### Homology Modelling

Homology models for the BceS HAMP and DHp domains (positions 56 to 174), and for the C-terminal domain of BceAB (from position 507) were generated using I-TASSER with default parameters (Zhang, 2008; Yang *et al.*, 2015). The respective homology model with the highest C-score was selected, and the top-ranking structural analog from PDB identified by I-TASSER was noted for reference. For quality control, the resulting homology model for BceS was compared to its closest structural analog, PDB 3RZW, using the TM-align structural alignment programme implemented in I-TASSER (Zhang, 2008; Yang *et al.*, 2015). The obtained TM-score was 0.771 (where >0.5 assumes the same fold), with a root mean square deviation (RMSD) of 1.68 Å between the structurally aligned residues. The homology model was analysed for Ramachandran outliers using MolProbity (Williams *et al.*, 2017). Eight outliers were identified, all within unstructured loop regions of the model: six fall in the loop connecting the two helices in the HAMP-like domain, and two in the loop connecting the DHp domain helices. The homology model was therefore regarded as a plausible representation of the BceS structure, but with some uncertainty in the linking loops between helical segments.

### Gaussian-accelerated molecular dynamics (GaMD) Simulations

A homology model of the cytoplasmic region of BceS consisting of the DHp and ATPase domains was constructed using I-TASSER as before (Zhang, 2008; Yang *et al.*, 2015). The HAMP-like domain was excluded due to low sequence homology with conventional HAMP domains. The approximate orientation of each BceS monomer with respect to one another was predicted based on cysteine cross-linking data (Fig. 2) and further refined using Z-Dock to perform rigid body protein-protein docking (Pierce *et al.*, 2014), with subsequent side chain rotamers optimised using SCWRL4 (Krivov *et al.*, 2009). After MD equilibration of the dimer model (see below), no Ramachandran outliers were present in the loop connecting the DHp domain helices or elsewhere (only Gly262, located in a flexible loop in the ATPase domain, was just outside the ‘allowed region’). The N-terminus of the DHp domain of each monomer was acetylated to cap the α-helix and prevent having a positive charge at the termini. ATP was then docked into each ATP binding site using AutoDock Vina (exhaustiveness set to 32) (Trott and Olson, 2010). The generated model of BceS was then manually modified to construct the E115K/K116E variant. All residues were modelled in their standard protonation states, with His108 and His124 singly protonated on the Nδ1 nitrogen, and all other His singly protonated on the Nε2 nitrogen (as indicated by hydrogen bonding networks). Both systems were solvated in an octahedral water box such that no protein/substrate atom was within 10 Å of the box boundaries, with Na^+^ or Cl^−^ counter ions added as necessary to ensure an overall neutral charge. MD simulations were performed using GPU accelerated Amber16, with the ff14SB force field and TIP3P water model used to describe protein and water molecules respectively. Parameters for ATP were taken from a study by Meagher and colleagues (Meagher *et al.*, 2003). Following a minimisation, heating, NVT and NPT equilibration procedure (described below), Gaussian accelerated MD (GaMD) simulations were done in the NVT ensemble at 300 K (Miao *et al.*, 2015), with a boost potential applied to both the total system potential energy as well as the dihedral potential energies (see also GaMD settings in Table S3). To prepare for production GaMD simulations, a 12 ns long conventional MD run was used to obtain maximum, minimum, average and standard deviation of the system potential energies. Following this, a 66 ns long run was used to update the above parameters (updated every 600 ps) to ensure converged GaMD production parameters. Production GaMD simulations were then run for 4 × 500 ns, using different random seeds for each replica. MD simulations were run using a 2 fs time step with the SHAKE algorithm applied to any bond containing a hydrogen atom. An 8 Å direct space non-bonded cut-off was applied with long-range electrostatics evaluated using the particle mesh Ewald algorithm (Darden *et al.*, 1993). Temperature was regulated using Langevin temperature control (collision frequency of 1 ps^−1^). All MD and GaMD simulations were performed with a 10 kcal mol^−1^ restraint on the Phi and Psi dihedrals between the acetyl and first N-terminal residue, to prevent the α-helix fraying. Principal component analysis (PCA) and other trajectory analyses were performed within CPPTRAJ (Roe and Cheatham, 2013), on the Cα atoms of the DHp domain residues. PCA was performed on the trajectories after constructing an average structure of all four simulations combined. Reweighting of PCA and distance measurements (to obtain PMFs) was performed using the ‘PyReweighting’ toolkit (Miao *et al.*, 2014).

To equilibrate structures for GaMD simulations, the following procedure was performed. Minimisation of all hydrogen atoms and solvent molecules was done using 500 steps of steepest descent followed by 500 steps of conjugate gradient. To keep all other atoms (i.e. the protein heavy atoms) in place during the minimisation, 10 kcal mol^−1^ Å^−1^ positional restraints were applied. Retaining the positional restraints on all protein heavy atoms, the system was then heated rapidly from 50 K to 300 K in the NVT ensemble over the course of 200 ps. This system was again minimised for a further 500 steps of steepest descent followed by 500 steps of conjugate gradient, this time only applying positional restraints (of size 5 kcal mol^−1^ Å^−1^) to the Cα carbon atoms. These Cα restraints were retained as the system was again heated from 25 K to 300 K over the course of 50 ps in the NVT ensemble. Simulations were then performed in the NPT ensemble (1 atm, 300 K), first gradually reducing the 5 kcal mol^−1^ Å^−1^ Cα carbon restraints over the course of 50 ps of simulation time. This was done in five steps (5, 4, 3, 2, 1 kcal mol^−1^ Å^−1^) of 10 ps each. A final 20 ns long MD simulation was then performed, in which no restraints were used, to equilibrate the box size. All dynamics steps used the SHAKE algorithm. Simulations performed in the NVT ensemble used Langevin temperature control (with a collision frequency of 1 ps^−1^) and a simulation time-step of 1 fs. Simulations performed in the NPT ensemble again used Langevin temperature control (collision frequency of 1 ps^−1^) and a Berendsen barostat (1 ps pressure relaxation time), with a simulation time-step of 2 fs.

## ACKNOWLEDGEMENTS

The authors thank Roger Draheim for helpful advice in establishing the cysteine crosslinking assays. We also thank our undergraduate research students Jessica Lancaster and Adam McCartan for constructing plasmids pJL705 and pAM703, respectively. We thank Jean van den Elsen and Laurence Hurst for critical reading of and feedback on the manuscript.

Work in SG’s lab was supported by the Biotechnology and Biological Sciences Research Council (BBSRC; BB/M029255/1). MWK is a BBSRC David Phillips Fellow (BB/M026280/1).

## AUTHOR CONTRIBUTIONS

AK and MJG carried out all laboratory work and data analysis; MWK and CRP performed and analyzed the MD simulations; SG supervised all laboratory work and coordinated the research project; SG, CRP and AK wrote the manuscript.

## ABBREVIATED SUMMARY

One of the main counteracting forces working against the development and spread of antibiotic resistance is the associated cost in fitness, which can be minimized through careful gene regulation. We here investigated fine-tuned signalling via a sensory complex consisting of a transporter and a histidine kinase. We show how the kinase is activated upon antibiotic attack, that the transporter exerts complete control over the kinase, and also has itself evolved to conserve energy.

## REFERENCES

Andersson, D.I., and Hughes, D. (2010) Antibiotic resistance and its cost: is it possible to reverse resistance? Nat Rev Microbiol 8: 260–271.

Appleman, J.A., and Stewart, V. (2003) Mutational Analysis of a Conserved Signal-Transducing Element: the HAMP Linker of the *Escherichia coli* Nitrate Sensor NarX. J Bacteriol 185: 89–97.

Arnaud, M., Chastanet, A., and Débarbouillé, M. (2004) New vector for efficient allelic replacement in naturally nontransformable, low-GC-content, Gram-positive bacteria. Appl Environ Microbiol 70: 6887–6891.

Bhate, M.P., Lemmin, T., Kuenze, G., Mensa, B., Ganguly, S., Peters, J.M., et al. (2018) Structure and Function of the Transmembrane Domain of NsaS, an Antibiotic Sensing Histidine Kinase in *Staphylococcus aureus*. J Am Chem Soc 140: 7471–7485.

Bhate, M.P., Molnar, K.S., Goulian, M., and DeGrado, W.F. (2015a) Signal Transduction in Histidine Kinases: Insights from New Structures. Structure/Folding and Design 23: 981–994.

Bhate, M.P., Molnar, K.S., Goulian, M., and DeGrado, W.F. (2015b) Signal Transduction in Histidine Kinases: Insights from New Structures. Structure/Folding and Design 23: 981–994.

Crow, A., Greene, N.P., Kaplan, E., and Koronakis, V. (2017) Structure and mechanotransmission mechanism of the MacB ABC transporter superfamily. Proc Natl Acad Sci USA 23:201712153–6.

Darden, T., York, D., and Pedersen, L. (1993) Particle mesh Ewald: An N·log(N) method for Ewald sums in large systems. The Journal of Chemical Physics 98: 10089–10092.

Dintner, S., Heermann, R., Fang, C., Jung, K., and Gebhard, S. (2014) A sensory complex consisting of an ATP-binding cassette transporter and a two-component regulatory system controls bacitracin resistance in *Bacillus subtilis*. J Biol Chem 289: 27899–27910.

Dintner, S., Staroń, A., Berchtold, E., Petri, T., Mascher, T., and Gebhard, S. (2011) Coevolution of ABC transporters and two-component regulatory systems as resistance modules against antimicrobial peptides in Firmicutes bacteria. J Bacteriol 193: 3851–3862.

Durão, P., Balbontín, R., and Gordo, I. (2018) Evolutionary Mechanisms Shaping the Maintenance of Antibiotic Resistance. Trends Microbiol 26: 677–691.

Ferris, H.U., Dunin-Horkawicz, S., Hornig, N., Hulko, M., Martin, J., Schultz, J.E., et al. (2012) Mechanism of Regulation of Receptor Histidine Kinases. Structure/Folding and Design 20: 56–66.

Ferris, H.U., Dunin-Horkawicz, S., Mondéjar, L.G., Hulko, M., Hantke, K., Martin, J., et al. (2011) The Mechanisms of HAMP-Mediated Signaling in Transmembrane Receptors. Structure/Folding and Design 19: 378–385.

Fritz, G., Dintner, S., Treichel, N.S., Radeck, J., Gerland, U., Mascher, T., and Gebhard, S. (2015) A new way of sensing: need-based activation of antibiotic resistance by a flux-sensing mechanism. mBio 6: e00975–15.

Gao, R., and Stock, A.M. (2009) Biological insights from structures of two-component proteins. Annu Rev Microbiol 63: 133–154.

Gebhard, S. (2012) ABC transporters of antimicrobial peptides in Firmicutes bacteria – phylogeny, function and regulation. Mol Microbiol 86: 1295–1317.

Gebhard, S., and Mascher, T. (2011) Antimicrobial peptide sensing and detoxification modules: unravelling the regulatory circuitry of *Staphylococcus aureus*. Mol Microbiol 81: 581–587.

Gebhard, S., Fang, C., Shaaly, A., Leslie, D.J., Weimar, M.R., Kalamorz, F., et al. (2014) Identification and Characterization of a Bacitracin Resistance Network in *Enterococcus faecalis*. Antimicrob Agents Chemother 58: 1425–1433.

Greene, N.P., Kaplan, E., Crow, A., and Koronakis, V. (2018) Antibiotic Resistance Mediated by the MacB ABC Transporter Family: A Structural and Functional Perspective. Front Microbiol 9: 1923–17.

Gushchin, I., and Gordeliy, V. (2017) Transmembrane Signal Transduction in Two-Component Systems: Piston, Scissoring, or Helical Rotation? BioEssays 40: 1700197–10.

Gushchin, I., Melnikov, I., Polovinkin, V., Ishchenko, A., Yuzhakova, A., Buslaev, P., et al. (2017) Mechanism of transmembrane signaling by sensor histidine kinases. Science 356: eaah6345–9.

Harwood, C.R., and Cutting, S.M. (1990) Molecular biological methods for Bacillus. John Wiley & Sons, Chichester, UK.

Hiron, A., Falord, M., Valle, J., Débarbouillé, M., and Msadek, T. (2011) Bacitracin and nisin resistance in *Staphylococcus aureus:* a novel pathway involving the BraS/BraR two-component system (SA2417/SA2418) and both the BraD/BraE and VraD/VraE ABC transporters. Mol Microbiol 81: 602–622.

Ho, S.N., Hunt, H.D., Horton, R.M., Pullen, J.K., and Pease, L.R. (1989) Site-directed mutagenesis by overlap extension using the polymerase chain reaction. Gene 77: 51–59.

Hsieh, Y.-J., and Wanner, B.L. (2010) Global regulation by the seven-component P_i_ signaling system. Curr Opin Microbiol 13: 198–203.

Hulko, M., Berndt, F., Gruber, M., Linder, J.U., Truffault, V., Schultz, A., et al. (2006) The HAMP Domain Structure Implies Helix Rotation in Transmembrane Signaling. Cell 126: 929–940.

Jacob-Dubuisson, F., Mechaly, A., Betton, J.-M., and Antoine, R. (2018) Structural insights into the signalling mechanisms of two-component systems. Nat Rev Microbiol 16: 1–9.

Kallenberg, F., Dintner, S., Schmitz, R., and Gebhard, S. (2013) Identification of regions important for resistance and signalling within the antimicrobial peptide transporter BceAB of *Bacillus subtilis*. J Bacteriol 195: 3287–3297.

Kobras, C.M., Piepenbreier, H., Emenegger, J., Sim, A., Fritz, G., and Gebhard, S. (2020) BceAB-Type Antibiotic Resistance Transporters Appear To Act by Target Protection of Cell Wall Synthesis. Antimicrob Agents Chemother 64: 321–15.

Krivov, G.G., Shapovalov, M.V., and Dunbrack, R.L. (2009) Improved prediction of protein side-chain conformations with SCWRL4. Proteins 77: 778–795.

Lai, R.-Z., and Parkinson, J.S. (2014) Functional Suppression of HAMP Domain Signaling Defects in the *E. coli* Serine Chemoreceptor. J Mol Biol 426: 3642–3655.

Letunic, I., Doerks, T., and Bork, P. (2012) SMART 7: recent updates to the protein domain annotation resource. Nucleic Acids Res 40: D302–5.

Mascher, T. (2006) Intramembrane-sensing histidine kinases: a new family of cell envelope stress sensors in Firmicutes bacteria. FEMS Microbiol Lett 264: 133–144.

Matamouros, S., Hager, K.R., and Miller, S.I. (2015) HAMP Domain Rotation and Tilting Movements Associated with Signal Transduction in the PhoQ Sensor Kinase. mBio 6: e00616–15.

Meagher, K.L., Redman, L.T., and Carlson, H.A. (2003) Development of polyphosphate parameters for use with the AMBER force field. J Comput Chem 24: 1016–1025.

Miao, Y., Feher, V.A., and McCammon, J.A. (2015) Gaussian Accelerated Molecular Dynamics: Unconstrained Enhanced Sampling and Free Energy Calculation. J Chem Theory Comput 11: 3584–3595.

Miao, Y., Sinko, W., Pierce, L., Bucher, D., Walker, R.C., and McCammon, J.A. (2014) Improved Reweighting of Accelerated Molecular Dynamics Simulations for Free Energy Calculation. J Chem Theory Comput 10: 2677–2689.

Molnar, K.S., Bonomi, M., Pellarin, R., Clinthorne, G.D., Gonzalez, G., Goldberg, S.D., et al. (2014) Cys-scanning disulfide crosslinking and bayesian modeling probe the transmembrane signaling mechanism of the histidine kinase, PhoQ. Structure 22: 1239–1251.

Monzel, C., and Unden, G. (2015) Transmembrane signaling in the sensor kinase DcuS of *Escherichia coli:* A long-range piston-type displacement of transmembrane helix 2. Proc Natl Acad Sci USA 112: 11042–11047.

Ohki, R., Giyanto, Tateno, K., Masuyama, W., Moriya, S., Kobayashi, K., and Ogasawara, N. (2003) The BceRS two-component regulatory system induces expression of the bacitracin transporter, BceAB, in *Bacillus subtilis*. Mol Microbiol 49: 1135–1144.

Ouyang, J., Tian, X.L., Versey, J., Wishart, A., and Li, Y.H. (2010) The BceABRS Four-Component System Regulates the Bacitracin-Induced Cell Envelope Stress Response in *Streptococcus mutans*. Antimicrob Agents Chemother 54: 3895–3906.

Pakula, A.A., and Simon, M.I. (1992) Determination of transmembrane protein structure by disulfide cross-linking: the *Escherichia coli* Tar receptor. Proc Natl Acad Sci USA 89: 4144–4148.

Parkinson, J.S., Hazelbauer, G.L., and Falke, J.J. (2015) Signaling and sensory adaptation in *Escherichia coli* chemoreceptors: 2015 update. Trends Microbiol 23: 257–266.

Piepenbreier, H., Fritz, G., and Gebhard, S. (2017) Transporters as information processors in bacterial signalling pathways. Mol Microbiol 104: 1–15.

Pierce, B.G., Wiehe, K., Hwang, H., Kim, B.-H., Vreven, T., and Weng, Z. (2014) ZDOCK server: interactive docking prediction of protein-protein complexes and symmetric multimers. Bioinformatics 30: 1771–1773.

Radeck, J., Kraft, K., Bartels, J., Cikovic, T., Dürr, F., Emenegger, J., et al. (2013) The *Bacillus* BioBrick Box: generation and evaluation of essential genetic building blocks for standardized work with *Bacillus subtilis*. Journal of Biological Engineering 7: 29.

Revilla-Guarinos, A., Gebhard, S., Alcántara, C., Staroń, A., Mascher, T., and Zúñiga, M. (2013) Characterization of a regulatory network of peptide antibiotic detoxification modules in *Lactobacillus casei* BL23. Appl Environ Microbiol 79: 3160–3170.

Revilla-Guarinos, A., Gebhard, S., Mascher, T., and Zúñiga, M. (2014) Defence against antimicrobial peptides: different strategies in Firmicutes. Environ Microbiol 16: 1225–1237.

Rietkötter, E., Hoyer, D., and Mascher, T. (2008) Bacitracin sensing in *Bacillus subtilis*. Mol Microbiol 68: 768–785.

Roe, D.R., and Cheatham, T.E., III (2013) PTRAJ and CPPTRAJ: Software for Processing and Analysis of Molecular Dynamics Trajectory Data. J Chem Theory Comput 9: 3084–3095.

Salvi, M., Schomburg, B., Giller, K., Graf, S., Unden, G., Becker, S., et al. (2017) Sensory domain contraction in histidine kinase CitA triggers transmembrane signaling in the membrane-bound sensor. Proc Natl Acad Sci USA 114: 3115–3120.

Schindelin, J., Arganda-Carreras, I., Frise, E., Kaynig, V., Longair, M., Pietzsch, T., et al. (2012) Fiji: an open-source platform for biological-image analysis. Nature Methods 9: 676–682.

Staroń, A., Finkeisen, D.E., and Mascher, T. (2011) Peptide antibiotic sensing and detoxification modules of *Bacillus subtilis*. Antimicrob Agents Chemother 55: 515–525.

Tetsch, L., and Jung, K. (2009) The regulatory interplay between membrane-integrated sensors and transport proteins in bacteria. Mol Microbiol 73: 982–991.

Trott, O., and Olson, A.J. (2010) AutoDock Vina: improving the speed and accuracy of docking with a new scoring function, efficient optimization, and multithreading. J Comput Chem 31: 455–461.

Unden, G., Wörner, S., and Monzel, C. (2016) Cooperation of secondary transporters and sensor kinases in transmembrane signalling: the DctA/DcuS and DcuB/DcuS sensor complexes of *Escherichia coli*. Adv Microb Physiol 68: 139–167.

Waglechner, N., and Wright, G.D. (2017) Antibiotic resistance: it’s bad, but why isn’t it worse? BMC Biol 15: 1–8.

Wang, B., Zhao, A., Novick, R.P., and Muir, T.W. (2014) Activation and inhibition of the receptor histidine kinase AgrC occurs through opposite helical transduction motions. Molecular Cell 53: 929–940.

Williams, C.J., Headd, J.J., Moriarty, N.W., Prisant, M.G., Videau, L.L., Deis, L.N., et al. (2017) MolProbity: More and better reference data for improved all-atom structure validation. Protein Science 27: 293–315.

Witan, J., Bauer, J., Wittig, I., Steinmetz, P.A., Erker, W., and Unden, G. (2012) Interaction of the *Escherichia coli* transporter DctA with the sensor kinase DcuS: presence of functional DctA/DcuS sensor units. Mol Microbiol 85: 846–861.

Yang, J., Yan, R., Roy, A., Xu, D., Poisson, J., and Zhang, Y. (2015) The I-TASSER Suite: protein structure and function prediction. Nature Methods 12: 7–8.

Zhang, Y. (2008) I-TASSER server for protein 3D structure prediction. BMC Bioinformatics 9: 76–8.

Zhou, Q., Ames, P., and Parkinson, J.S. (2011) Biphasic control logic of HAMP domain signalling in the *Escherichia coli* serine chemoreceptor. Mol Microbiol 80: 596–611.

