## Supplementary material for "Conformation control of the histidine kinase BceS of *Bacillus subtilis* by its cognate ABC-transporter facilitates need-based activation of antibiotic resistance"

### - Supplementary Information -

**Figure S1.** Cysteine scanning and site-directed mutagenesis of the BceS DHp domain.

**Figure S2.** Cysteine cross-linking following activation of signaling.

**Figure S3.** Equilibration and principle component analysis of Gaussian accelerated molecular dynamics simulations of BceS DHp domain conformation.

**Figure S4.** Cysteine scanning mutagenesis of the BceS HAMP-like domain.

**Figure S5.** Cysteine scanning mutagenesis of the second transmembrane helix of BceS.

**Figure S6.** Arginine substitutions in transmembrane helix 2 of BceS.

**Figure S7.** Signaling phenotypes of BceAB variants.

**Table S1.** Strains, primers and plasmids used in this study.

**Table S2.** Minimal inhibitory concentration of bacitracin in strains carrying BceAB variants.

**Table S3.** Parameters used for GaMD simulations.

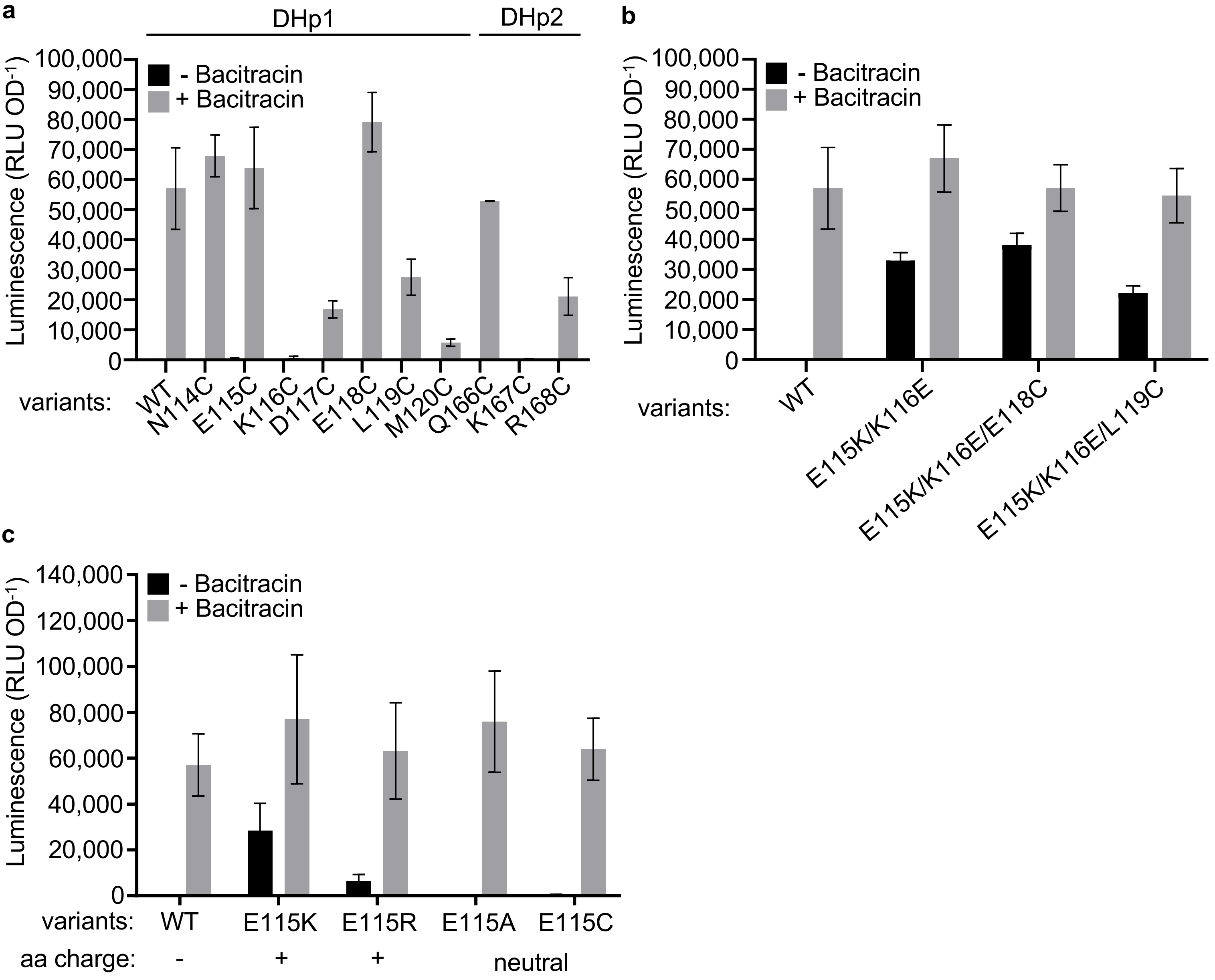


**Figure S1. Cysteine scanning and site-directed mutagenesis of the BceS DHp domain.** BceS signalling activity measured in strains harbouring $P_{bceA}$-*luxABCDE* and $P_{xyl}$-*bceS* wild-type (WT) or variants as indicated. Amino acid substitutions were introduced in the parent construct *bceS*$^{WT}$-His$_8$ (SGB369). **a,** Single Cys-substitution variants. Residues belonging to helices DHp1 or DHp2 are indicated above. **b,** Combination of single Cys substitutions with the constitutive ON substitutions E115K/K116E. **c,** Series of substitutions at position 115, with the resulting side-chain charge indicated below. Signalling was measured in cells challenged with 1 µg ml$^{-1}$ of bacitracin (grey) or left unchallenged (black). Data are shown as mean ± SD from 4-22 biological repeats.

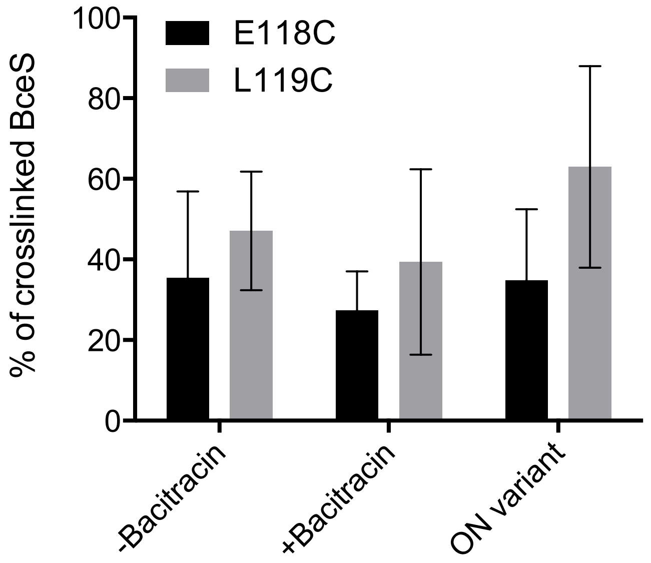


**Figure S2. Cysteine cross-linking following activation of signalling.** The percentage of crosslinking for BceS Cys variants E118C (black) and L119C (grey) was calculated from relative band intensities between monomer and dimer within each lane, following Western blot detection of BceS-His$_8$ as described in Methods. '+/-Bacitracin' shows results for cells producing otherwise wild-type BceS following activation with the native stimulus. 'ON variant' shows results in cells carrying the additional substitutions E115K/K116E causing constitutive activity of BceS.

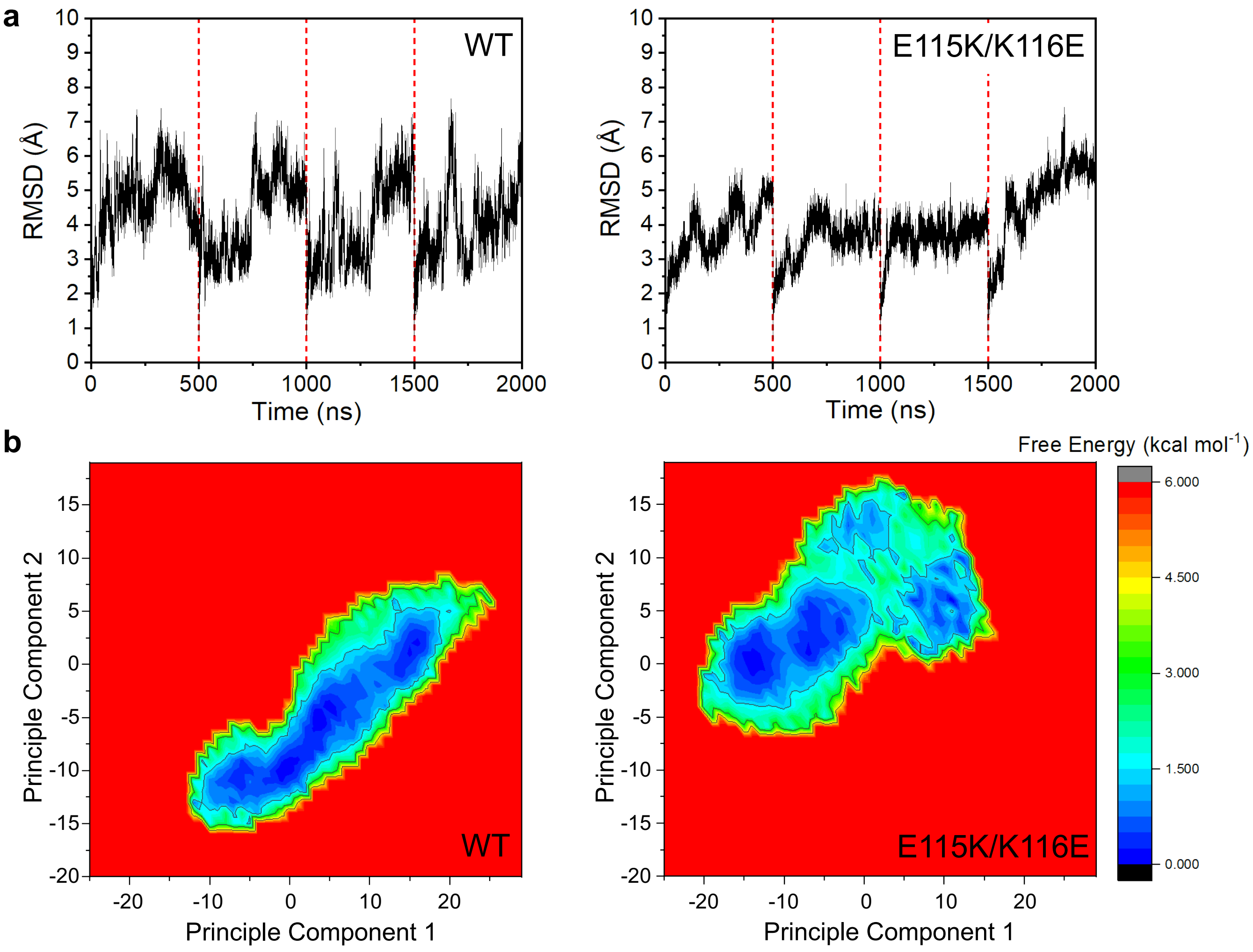


**Figure S3. Equilibration and principle component analysis of Gaussian accelerated molecular dynamics (GAMD) simulations of BceS DHp domain conformation. a,** Cα RMSD over the course of GAMD simulations for simulations of wild type (WT, left) and the E115K/K116E variant (right) protein. The red dotted lines divides up the four different 500 ns long replicas used. RMSD was measured against the homology model starting structure. **b,** Projection of the first two principal components (PCs) from PC analysis (PCA) of the GAMD simulations for the WT (left) and E115K/K116E variant (right) protein. PCA was performed on the Cα carbon of the DHp domain residues.

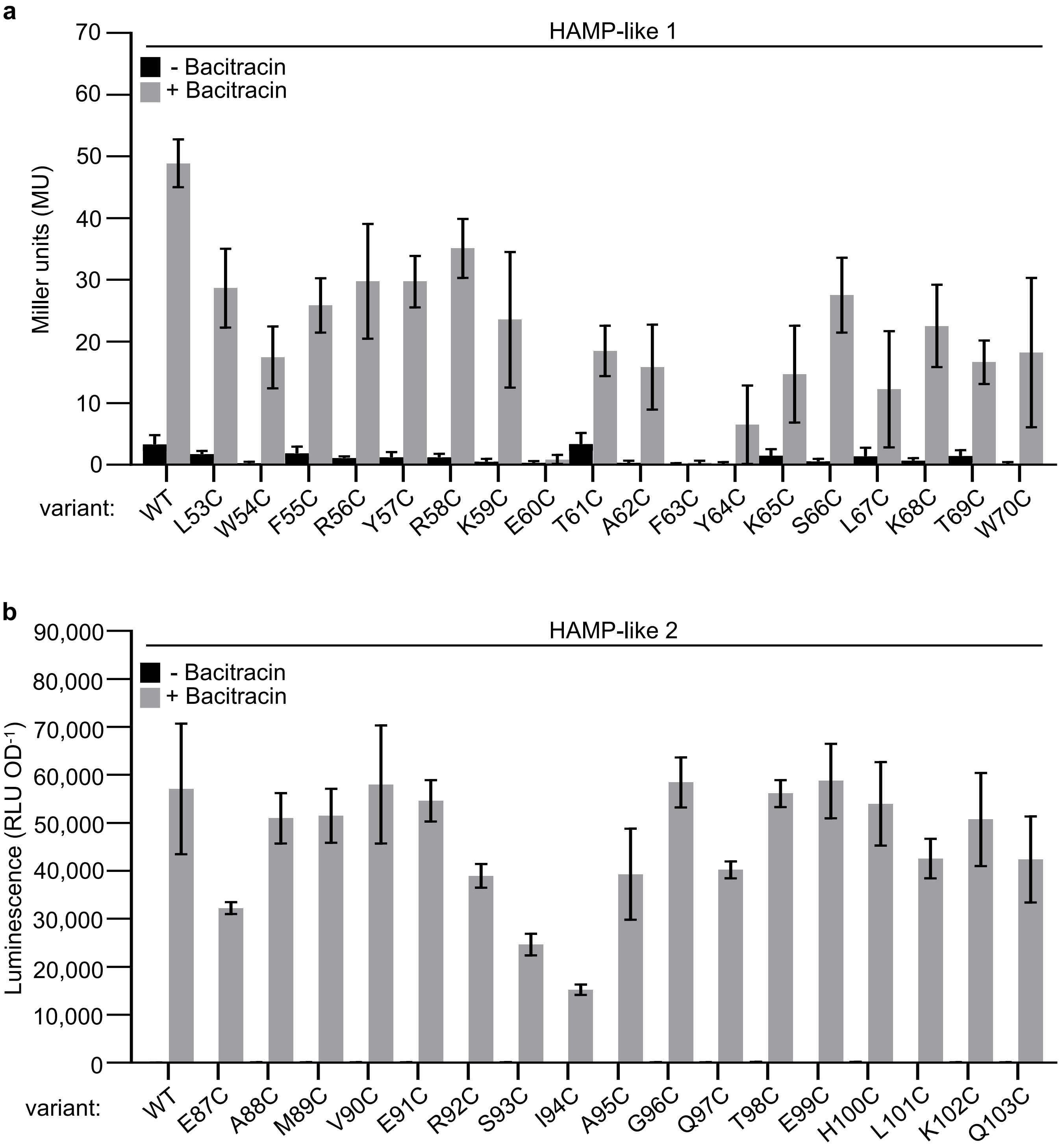


**Figure S4: Cysteine scanning mutagenesis of the BceS HAMP-like domain.** Amino acid substitutions were introduced into the $P_{xyl}$-*bceS*-His$_8$ parent construct, pSD2E01. Cells were challenged with 1 µg ml$^{-1}$ of bacitracin (grey) or left unchallenged (black). **a,** Cys scanning of the helix HAMP-like 1. Signalling was assessed by β-galactosidase activity (Miller Units) in cells carrying the $P_{bceA}$-*lacZ* reporter (SGB401 and derivatives). **b,** Cys scanning of helix HAMP-like 2. Signalling was assessed by luminescence (RLU OD$^{-1}$) in cells carrying the $P_{bceA}$-*luxABCDE* reporter (SGB369 and derivatives). Data are shown as mean ± SD from 3-22 independent repeats.

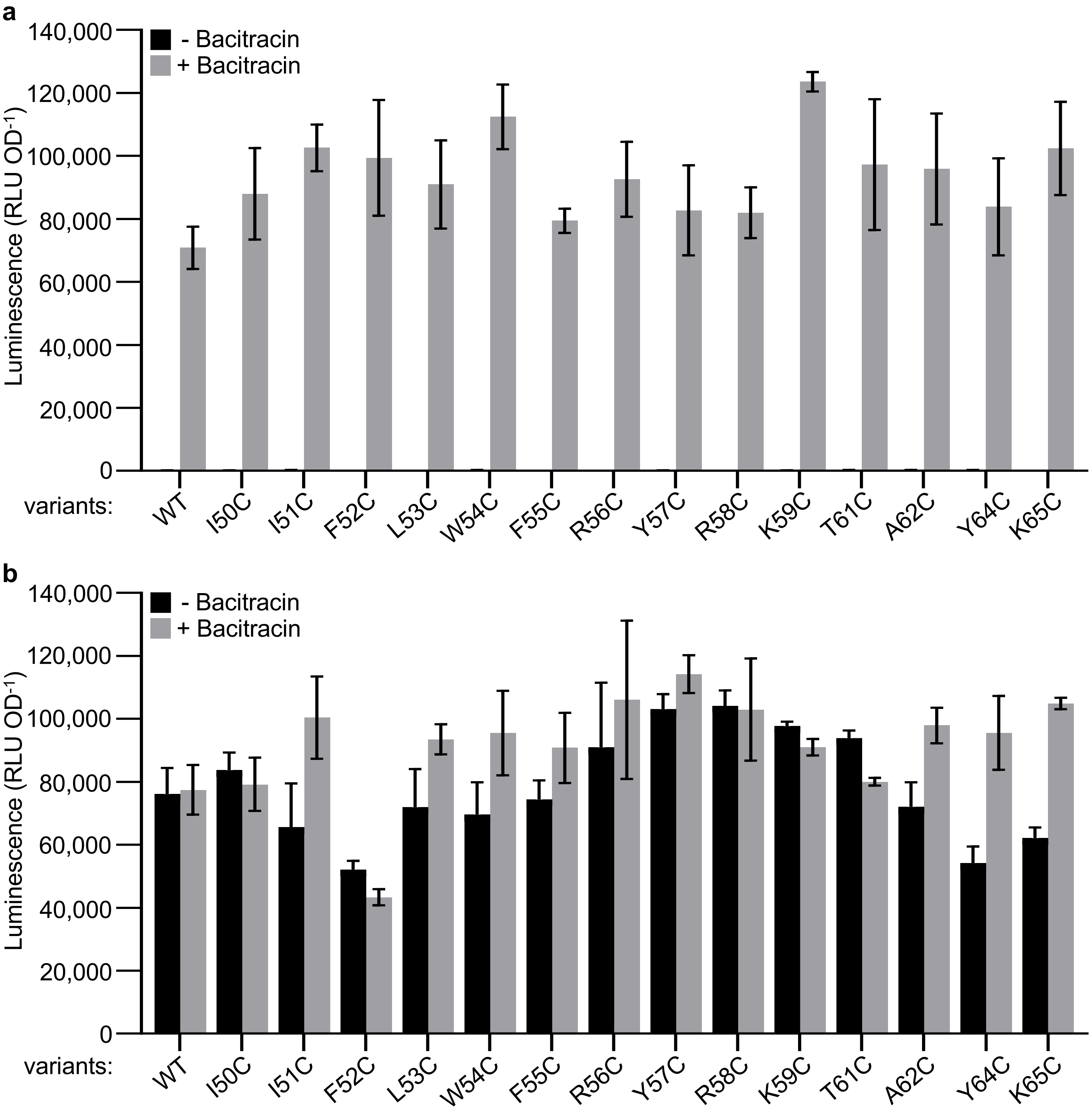


**Figure S5. Cysteine scanning mutagenesis of the second transmembrane helix of BceS.** BceS signaling activity was assessed in cells harbouring the $P_{bceA}$-*luxABCDE* reporter and $P_{xyl}$-*bceS*-$His_8$. Cells were challenged with 1 µg $ml^{-1}$ of bacitracin (grey) or left unchallenged (black), and signalling measured by luminescence (RLU $OD^{-1}$). **a,** Signalling activities of the otherwise wild-type, cysteine-free variant $BceS^{C45S/C198S/C259S}$ (WT, SGB936) and its single-cysteine derivatives as indicated. **b,** Signalling activities of the constitutively active, cysteine-free variant $BceS^{E115K/K116E/C45S/C198S/C259S}$ (WT; SGB951) and its single-cysteine derivatives as indicated. Data are shown as mean ± SD from 3 independent repeats.

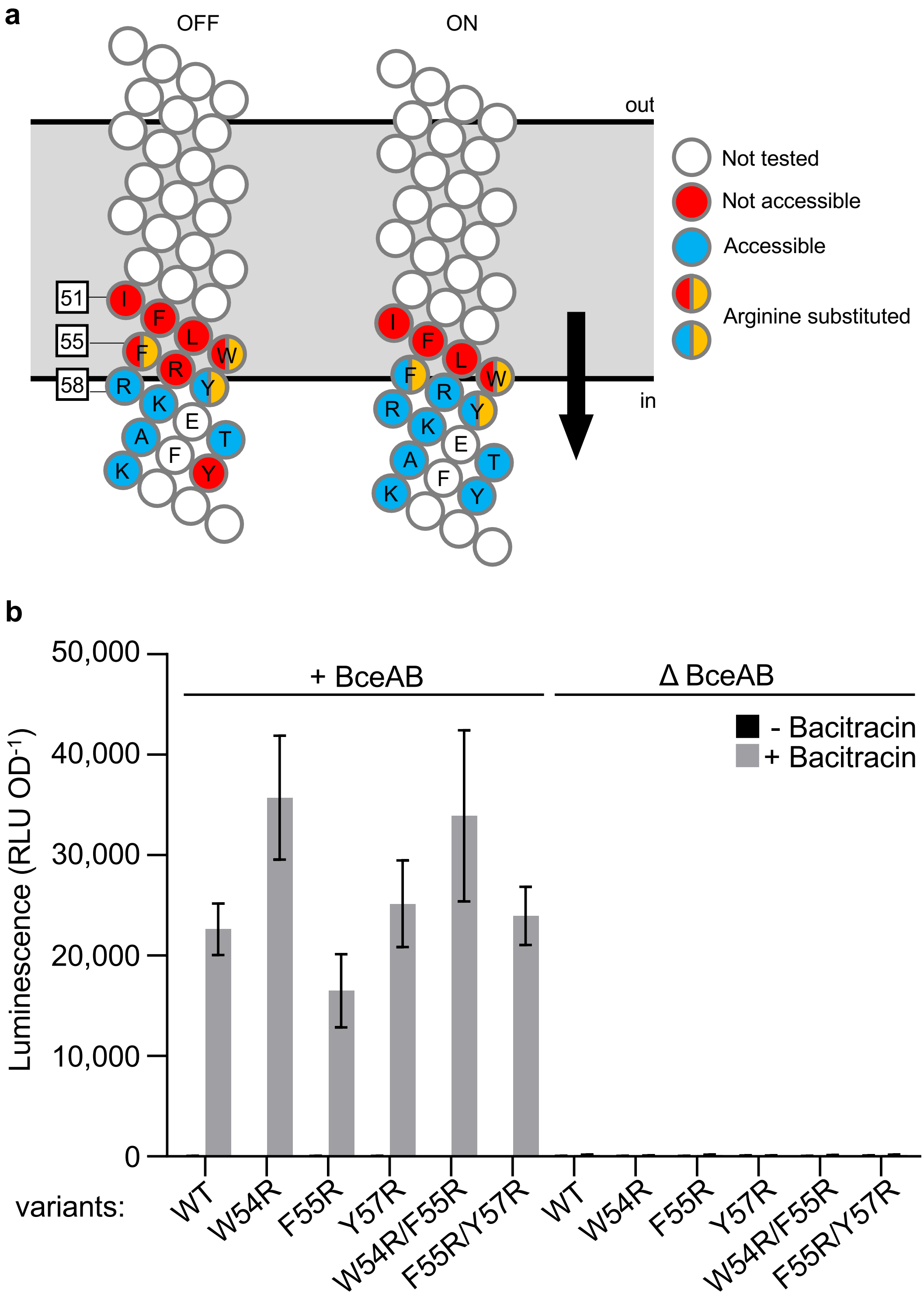


**Figure S6. Arginine substitutions in transmembrane helix 2 of BceS. a,** Schematic representation of Arg substitutions within transmembrane helix 2, shown in the OFF (left) and ON (right) states. Solvent-accessible Cys residues are indicated in blue, inaccessible Cys residues in red. The introduced Arg substitution are shown as yellow half-circles. **b,** Signalling activity of BceS Arg variants. Amino acid substitutions were introduced into the parent construct $P_{xyl}$-*bceS*-$His_8$, in a strain background carrying a Δ*bceS* deletion (+BceAB; SGB792), or a Δ*bceSAB* deletion (ΔBceAB; SGB818). Cells were challenged with 10 µg $ml^{-1}$ of bacitracin (grey) or left unchallenged (black), and signalling was assessed using the $P_{bceA}$-*luxABCDE* reporter, expressed as luminescence (RLU $OD^{-1}$). Data are shown as mean ± SD from 7-12 independent repeats.

a

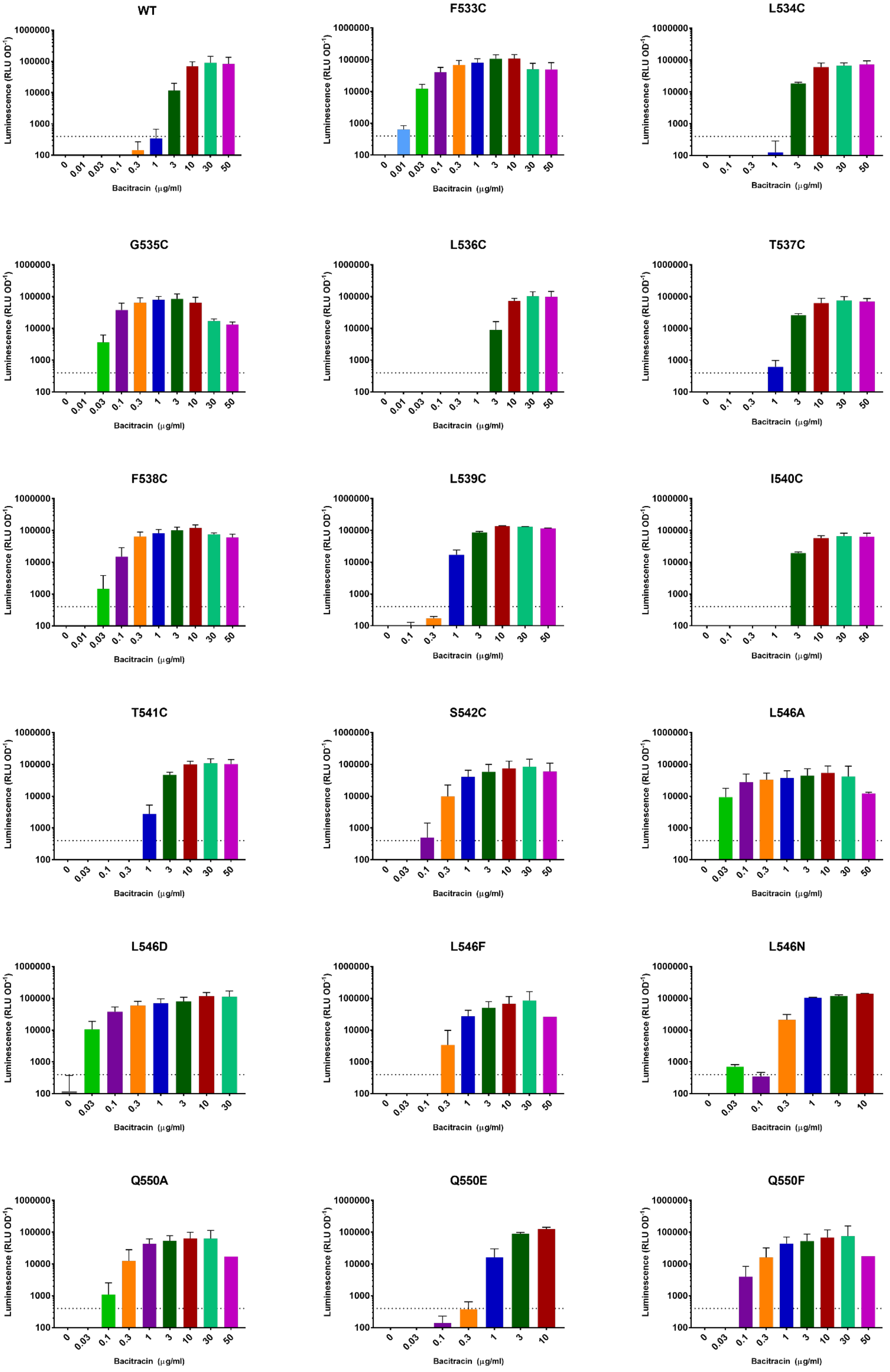


**Figure S7. Signalling phenotypes of BceAB variants.** *complete legend after panel b.*

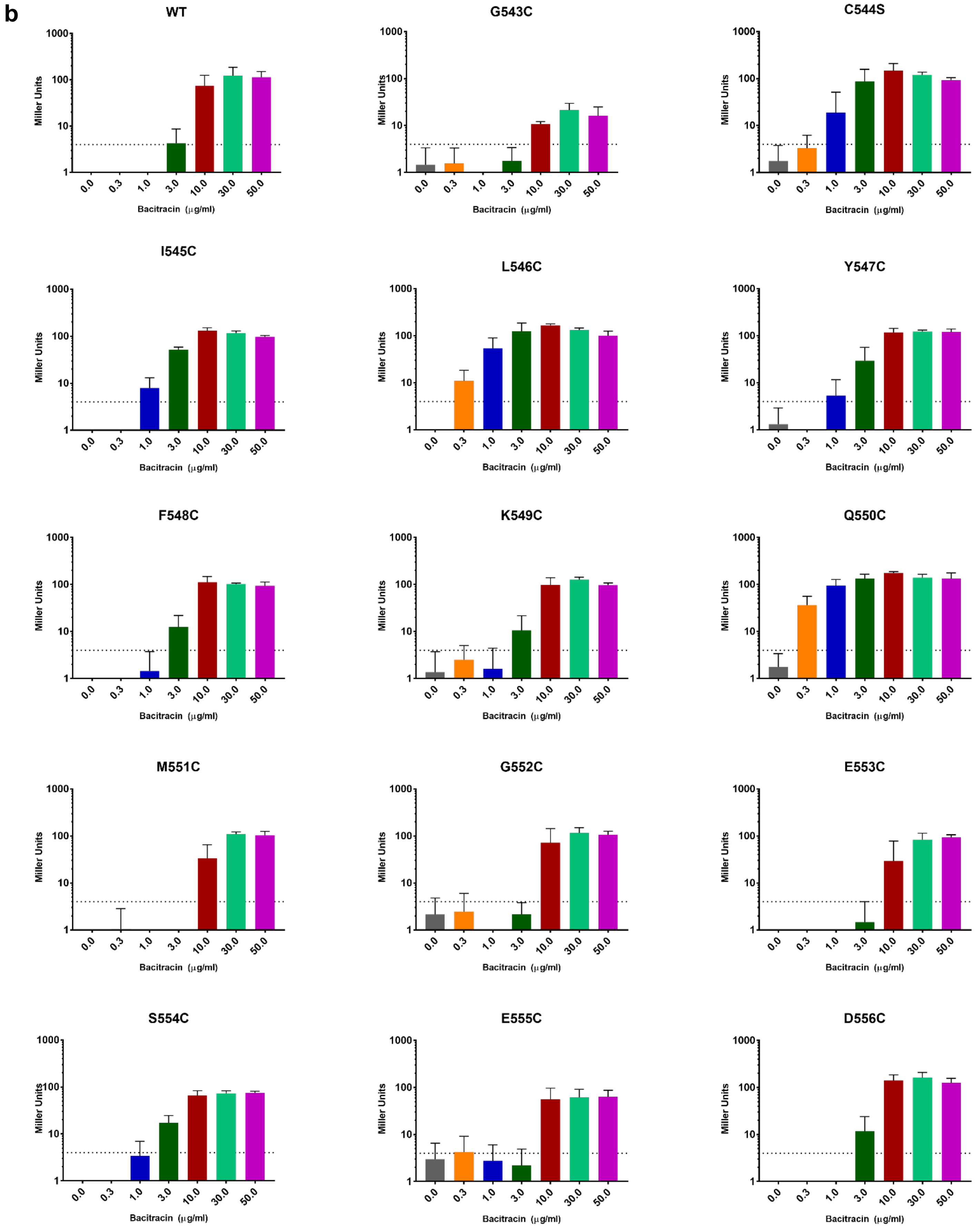


**Figure S7. Signalling phenotypes of BceAB variants.** Amino acid substitutions were introduced into the $P_{xyl}$-*bceAB*-$FLAG_3$ parent construct, pFK727. Cells were challenged with a series of bacitracin concentrations as indicated. **a,** Signalling was assessed by luminescence (RLU $OD^{-1}$) in cells carrying the $P_{bceA}$-*luxABCDE* reporter (SGB731 and derivatives). **b,** Signalling was assessed by b-galactosidase activity (Miller Units) in cells carrying the $P_{bceA}$-*lacZ* reporter (SGB370 and derivatives). Data are shown as mean ± SD of a minimum of three independent repeats. The threshold activity applied to determine signalling above background is indicated by dotted lines and was used to generate the data displayed in Fig. 6b of the main manuscript.

**Table S1. Strains, primers and plasmids used in this study**

**Section A - Strains**

| Strain | Genotype | Parent strain[a] | Plasmid/ Strain used[b] | Source |
|---|---|---|---|---|
| SGB16 | *W168 bceAB::kan amyE::bceA-lacZ* | TMB035 | pER603 | this work |
| SGB79 | *W168 bceAB::kan sacA::PbceA-luxABCDE* | | | (Kallenberg *et al.*, 2013) |
| SGB176 | *W168 bceAB::kan thrC::Pxyl-bceAB-FLAG* | | | (Kallenberg *et al.*, 2013) |
| SGB276 | *W168 ΔbceS sacA::PbceA-luxABCDE lacA::Pxyl-bceS-His8* | | | (Dintner *et al.*, 2014) |
| SGB334 | *W168 bceAB::kan amyE::PbceA-lacZ thrC:: pXT-BceAB*$^{E553C}$*-FLAG* | SGB16 | pMG707 | this work |
| SGB337 | *W168 bceAB::kan amyE::PbceA-lacZ thrC:: pXT-BceAB*$^{L546C}$*-FLAG* | SGB16 | pAM703 | this work |
| SGB338 | *W168 bceAB::kan amyE::PbceA-lacZ thrC:: pXT-BceAB*$^{D556C}$*-FLAG* | SGB16 | pJL705 | this work |
| SGB340 | *W168 sacA::PbceA-luxABCDE* | W168 | SGB276 | this work |
| SGB341 | *W168 sacA::PbceA-luxABCDE thrC::Pxyl-bceAB-FLAG* | SGB340 | SGB176 | this work |
| SGB345 | *W168 bceAB::kan amyE::PbceA-lacZ thrC:: pXT-BceAB*$^{Y547C}$*-FLAG* | SGB16 | pMG704 | this work |
| SGB352 | *W168 bceAB::kan amyE::PbceA-lacZ thrC:: pXT-BceAB*$^{K549C}$*-FLAG* | SGB16 | pMG713 | this work |
| SGB357 | *W168 ΔbceSAB sacA::PbceA-luxABCDE thrC::Pxyl-bceAB-FLAG* | SGB341 | pAK102 | this work |
| SGB359 | *W168 ΔbceSAB sacA::PbceA-luxABCDE thrC::Pxyl-bceAB-FLAG lacA::Pxyl-bceS*$^{E115K}$*-His8* | SGB357 | pAK2E02 | this work |
| SGB369 | *W168 ΔbceSAB sacA::PbceA-luxABCDE thrC::Pxyl-bceAB-FLAG lacA::Pxyl-bceS*$^{WT}$*-His8* | SGB357 | pSD2E01 | this work |
| SGB370 | *W168 bceAB::kan amyE::PbceA-lacZ thrC:: pXT-BceAB-FLAG* | SGB16 | pFK727 | this work |
| SGB371 | *W168 bceAB::kan amyE::PbceA-lacZ thrC:: pXT-BceAB*$^{G543C}$*-FLAG* | SGB16 | pMG718 | this work |
| SGB372 | *W168 bceAB::kan amyE::PbceA-lacZ thrC:: pXT-BceAB*$^{I545C}$*-FLAG* | SGB16 | pMG719 | this work |
| SGB373 | *W168 bceAB::kan amyE::PbceA-lacZ thrC:: pXT-BceAB*$^{F548C}$*-FLAG* | SGB16 | pMG720 | this work |
| SGB374 | *W168 bceAB::kan amyE::PbceA-lacZ thrC:: pXT-BceAB*$^{Q550C}$*-FLAG* | SGB16 | pMG721 | this work |
| SGB375 | *W168 bceAB::kan amyE::PbceA-lacZ thrC:: pXT-BceAB*$^{M551C}$*-FLAG* | SGB16 | pMG722 | this work |
| SGB376 | *W168 bceAB::kan amyE::PbceA-lacZ thrC:: pXT-BceAB*$^{G552C}$*-FLAG* | SGB16 | pMG723 | this work |
| SGB377 | *W168 ΔbceSAB* | W168 | pAK102 | this work |
| SGB378 | *W168 ΔbceSAB sacA::PbceA-luxABCDE* | SGB340 | pAK102 | this work |
| SGB379 | *W168 bceAB::kan amyE::PbceA-lacZ thrC:: pXT-BceAB*$^{C544S}$*-FLAG* | SGB16 | pMG724 | this work |
| SGB380 | *W168 ΔbceSAB sacA::PbceA-luxABCDE thrC::Pxyl-bceAB-FLAG lacA::Pxyl-bceS*$^{E115A}$*-His8* | SGB357 | pAK2E12 | this work |
| SGB381 | *W168 ΔbceSAB sacA::PbceA-luxABCDE thrC::Pxyl-bceAB-FLAG lacA::Pxyl-bceS*$^{E115R}$*-His8* | SGB357 | pAK2E13 | this work |
| SGB382 | *W168 ΔbceSAB sacA::PbceA-luxABCDE thrC::Pxyl-bceAB-FLAG lacA::Pxyl-bceS*$^{E115C}$*-His8* | SGB357 | pAK2E14 | this work |

| | | | | |
|---|---|---|---|---|
| SGB383 | *W168 ΔbceSAB sacA::PbceA-luxABCDE thrC::Pxyl-bceAB-FLAG lacA::Pxyl-bceS*$^{Q166C}$*-His8* | SGB357 | pAK2E15 | this work |
| SGB384 | *W168 ΔbceSAB sacA::PbceA-luxABCDE thrC::Pxyl-bceAB-FLAG lacA::Pxyl-bceS*$^{R168C}$*-His8* | SGB357 | pAK2E16 | this work |
| SGB385 | *W168 ΔbceSAB sacA::PbceA-luxABCDE thrC::Pxyl-bceAB-FLAG lacA::Pxyl-bceS*$^{K116C}$*-His8* | SGB357 | pAK2E17 | this work |
| SGB392 | *W168 ΔbceSAB sacA::PbceA-luxABCDE lacA::Pxyl-bceS*$^{WT}$*-His8* | SGB378 | pSD2E01 | this work |
| SGB395 | *W168 ΔbceSAB sacA::PbceA-luxABCDE thrC::Pxyl-bceAB-FLAG lacA::Pxyl-bceS*$^{N114C}$*-His8* | SGB357 | pAK2E21 | this work |
| SGB396 | *W168 ΔbceSAB sacA::PbceA-luxABCDE thrC::Pxyl-bceAB-FLAG lacA::Pxyl-bceS*$^{K167C}$*-His8* | SGB357 | pAK2E22 | this work |
| SGB400 | *W168 ΔbceSAB amyE::PbceA-lacZ thrC::Pxyl-bceAB*$^{WT}$*-FLAG* | SGB377 | TMB279 and SGB176 | this work |
| SGB401 | *W168 ΔbceSAB amyE::PbceA-lacZ thrC::Pxyl-bceAB*$^{WT}$*-FLAG lacA::Pxyl-bceS*$^{WT}$*-His8* | SGB400 | pSD2E01 | this work |
| SGB402 | *W168 ΔbceSAB amyE::PbceA-lacZ thrC::Pxyl-bceAB*$^{WT}$*-FLAG lacA::Pxyl-bceS*$^{L53C}$*-His8* | SGB400 | pMG2E01 | this work |
| SGB403 | *W168 ΔbceSAB amyE::PbceA-lacZ thrC::Pxyl-bceAB*$^{WT}$*-FLAG lacA::Pxyl-bceS*$^{W54C}$*-His8* | SGB400 | pMG2E02 | this work |
| SGB404 | *W168 ΔbceSAB amyE::PbceA-lacZ thrC::Pxyl-bceAB*$^{WT}$*-FLAG lacA::Pxyl-bceS*$^{F55C}$*-His8* | SGB400 | pMG2E03 | this work |
| SGB405 | *W168 ΔbceSAB amyE::PbceA-lacZ thrC::Pxyl-bceAB*$^{WT}$*-FLAG lacA::Pxyl-bceS*$^{R56C}$*-His8* | SGB400 | pMG2E04 | this work |
| SGB406 | *W168 ΔbceSAB amyE::PbceA-lacZ thrC::Pxyl-bceAB*$^{WT}$*-FLAG lacA::Pxyl-bceS*$^{Y57C}$*-His8* | SGB400 | pMG2E05 | this work |
| SGB407 | *W168 ΔbceSAB amyE::PbceA-lacZ thrC::Pxyl-bceAB*$^{WT}$*-FLAG lacA::Pxyl-bceS*$^{R58C}$*-His8* | SGB400 | pMG2E06 | this work |
| SGB408 | *W168 ΔbceSAB amyE::PbceA-lacZ thrC::Pxyl-bceAB*$^{WT}$*-FLAG lacA::Pxyl-bceS*$^{K59C}$*-His8* | SGB400 | pMG2E07 | this work |
| SGB409 | *W168 ΔbceSAB amyE::PbceA-lacZ thrC::Pxyl-bceAB*$^{WT}$*-FLAG lacA::Pxyl-bceS*$^{E60C}$*-His8* | SGB400 | pMG2E08 | this work |
| SGB410 | *W168 ΔbceSAB amyE::PbceA-lacZ thrC::Pxyl-bceAB*$^{WT}$*-FLAG lacA::Pxyl-bceS*$^{T61C}$*-His8* | SGB400 | pMG2E09 | this work |
| SGB411 | *W168 ΔbceSAB amyE::PbceA-lacZ thrC::Pxyl-bceAB*$^{WT}$*-FLAG lacA::Pxyl-bceS*$^{A62C}$*-His8* | SGB400 | pMG2E10 | this work |
| SGB413 | *W168 ΔbceSAB amyE::PbceA-lacZ thrC::Pxyl-bceAB*$^{WT}$*-FLAG lacA::Pxyl-bceS*$^{E60C/E115K}$*-His8* | SGB400 | pMG2E12 | this work |
| SGB414 | *W168 ΔbceSAB amyE::PbceA-lacZ thrC::Pxyl-bceAB*$^{WT}$*-FLAG lacA::Pxyl-bceS*$^{F63C}$*-His8* | SGB400 | pAK2E54 | this work |
| SGB415 | *W168 ΔbceSAB amyE::PbceA-lacZ thrC::Pxyl-bceAB*$^{WT}$*-FLAG lacA::Pxyl-bceS*$^{Y64C}$*-His8* | SGB400 | pAK2E55 | this work |
| SGB416 | *W168 ΔbceSAB amyE::PbceA-lacZ thrC::Pxyl-bceAB*$^{WT}$*-FLAG lacA::Pxyl-bceS*$^{K65C}$*-His8* | SGB400 | pAK2E56 | this work |
| SGB417 | *W168 ΔbceSAB amyE::PbceA-lacZ thrC::Pxyl-bceAB*$^{WT}$*-FLAG lacA::Pxyl-bceS*$^{S66C}$*-His8* | SGB400 | pAK2E57 | this work |
| SGB418 | *W168 ΔbceSAB amyE::PbceA-lacZ thrC::Pxyl-bceAB*$^{WT}$*-FLAG lacA::Pxyl-bceS*$^{L67C}$*-His8* | SGB400 | pAK2E58 | this work |
| SGB419 | *W168 ΔbceSAB amyE::PbceA-lacZ thrC::Pxyl-bceAB*$^{WT}$*-FLAG lacA::Pxyl-bceS*$^{K68C}$*-His8* | SGB400 | pAK2E59 | this work |
| SGB420 | *W168 ΔbceSAB amyE::PbceA-lacZ thrC::Pxyl-bceAB*$^{WT}$*-FLAG lacA::Pxyl-bceS*$^{T69C}$*-His8* | SGB400 | pAK2E60 | this work |
| SGB421 | *W168 ΔbceSAB amyE::PbceA-lacZ thrC::Pxyl-bceAB*$^{WT}$*-FLAG lacA::Pxyl-bceS*$^{W70C}$*-His8* | SGB400 | pAK2E61 | this work |
| SGB424 | *W168 ΔbceSAB amyE::PbceA-lacZ thrC::Pxyl-bceAB*$^{WT}$*-FLAG lacA::Pxyl-bceS*$^{F63C/E115K}$*-His8* | SGB400 | pMG2E15 | this work |
| SGB430 | *W168 bceAB::kan amyE::PbceA-lacZ thrC:: pXT-BceAB*$^{S554C}$*-FLAG* | SGB16 | pMG738 | this work |

| | | | | |
|---|---|---|---|---|
| SGB431 | *W168 bceAB::kan amyE::PbceA-lacZ thrC:: pXT-BceAB*$^{E555C}$*-FLAG* | SGB16 | pMG739 | this work |
| SGB433 | *W168 ΔbceSAB sacA::PbceA-luxABCDE thrC::Pxyl-bceAB-FLAG lacA::Pxyl-bceS*$^{K116E}$*-His8* | SGB357 | pAK2E26 | this work |
| SGB436 | *W168 ΔbceSAB sacA::PbceA-luxABCDE thrC::Pxyl-bceAB-FLAG lacA::Pxyl-bceS*$^{K167D}$*-His8* | SGB357 | pAK2E29 | this work |
| SGB440 | *W168 ΔbceSAB sacA::PbceA-luxABCDE thrC::Pxyl-bceAB-FLAG lacA::Pxyl-bceS*$^{E118C}$*-His8* | SGB357 | pAK2E33 | this work |
| SGB441 | *W168 ΔbceSAB sacA::PbceA-luxABCDE thrC::Pxyl-bceAB-FLAG lacA::Pxyl-bceS*$^{E115K/K116E}$*-His8* | SGB357 | pAK2E34 | this work |
| SGB444 | *W168 ΔbceSAB sacA::PbceA-luxABCDE thrC::Pxyl-bceAB-FLAG lacA::Pxyl-bceS*$^{I94C}$*-His8* | SGB357 | pAK2E37 | this work |
| SGB445 | *W168 ΔbceSAB sacA::PbceA-luxABCDE thrC::Pxyl-bceAB-FLAG lacA::Pxyl-bceS*$^{A95C}$*-His8* | SGB357 | pAK2E38 | this work |
| SGB446 | *W168 ΔbceSAB sacA::PbceA-luxABCDE thrC::Pxyl-bceAB-FLAG lacA::Pxyl-bceS*$^{G96C}$*-His8* | SGB357 | pAK2E39 | this work |
| SGB447 | *W168 ΔbceSAB sacA::PbceA-luxABCDE thrC::Pxyl-bceAB-FLAG lacA::Pxyl-bceS*$^{Q97C}$*-His8* | SGB357 | pAK2E40 | this work |
| SGB448 | *W168 ΔbceSAB sacA::PbceA-luxABCDE thrC::Pxyl-bceAB-FLAG lacA::Pxyl-bceS*$^{T98C}$*-His8* | SGB357 | pAK2E41 | this work |
| SGB449 | *W168 ΔbceSAB sacA::PbceA-luxABCDE thrC::Pxyl-bceAB-FLAG lacA::Pxyl-bceS*$^{E99C}$*-His8* | SGB357 | pAK2E42 | this work |
| SGB450 | *W168 ΔbceSAB sacA::PbceA-luxABCDE thrC::Pxyl-bceAB-FLAG lacA::Pxyl-bceS*$^{H100C}$*-His8* | SGB357 | pAK2E43 | this work |
| SGB451 | *W168 ΔbceSAB sacA::PbceA-luxABCDE thrC::Pxyl-bceAB-FLAG lacA::Pxyl-bceS*$^{L101C}$*-His8* | SGB357 | pAK2E44 | this work |
| SGB452 | *W168 ΔbceSAB sacA::PbceA-luxABCDE thrC::Pxyl-bceAB-FLAG lacA::Pxyl-bceS*$^{K102C}$*-His8* | SGB357 | pAK2E45 | this work |
| SGB453 | *W168 ΔbceSAB sacA::PbceA-luxABCDE thrC::Pxyl-bceAB-FLAG lacA::Pxyl-bceS*$^{Q103C}$*-His8* | SGB357 | pAK2E46 | this work |
| SGB454 | *W168 ΔbceSAB sacA::PbceA-luxABCDE thrC::Pxyl-bceAB-FLAG lacA::Pxyl-bceS*$^{E87C}$*-His8* | SGB357 | pAK2E47 | this work |
| SGB455 | *W168 ΔbceSAB sacA::PbceA-luxABCDE thrC::Pxyl-bceAB-FLAG lacA::Pxyl-bceS*$^{A88C}$*-His8* | SGB357 | pAK2E48 | this work |
| SGB456 | *W168 ΔbceSAB sacA::PbceA-luxABCDE thrC::Pxyl-bceAB-FLAG lacA::Pxyl-bceS*$^{M89C}$*-His8* | SGB357 | pAK2E49 | this work |
| SGB457 | *W168 ΔbceSAB sacA::PbceA-luxABCDE thrC::Pxyl-bceAB-FLAG lacA::Pxyl-bceS*$^{V90C}$*-His8* | SGB357 | pAK2E50 | this work |
| SGB458 | *W168 ΔbceSAB sacA::PbceA-luxABCDE thrC::Pxyl-bceAB-FLAG lacA::Pxyl-bceS*$^{E91C}$*-His8* | SGB357 | pAK2E51 | this work |
| SGB459 | *W168 ΔbceSAB sacA::PbceA-luxABCDE thrC::Pxyl-bceAB-FLAG lacA::Pxyl-bceS*$^{R92C}$*-His8* | SGB357 | pAK2E52 | this work |
| SGB460 | *W168 ΔbceSAB sacA::PbceA-luxABCDE thrC::Pxyl-bceAB-FLAG lacA::Pxyl-bceS*$^{S93C}$*-His8* | SGB357 | pAK2E53 | this work |
| SGB465 | *W168 ΔbceSAB sacA::PbceA-luxABCDE thrC::Pxyl-bceAB-FLAG lacA::Pxyl-bceS*$^{L67C}$*-His8* | SGB357 | pAK2E58 | this work |
| SGB470 | *W168 ΔbceSAB sacA::PbceA-luxABCDE thrC::Pxyl-bceAB-FLAG lacA::Pxyl-bceS*$^{D117C}$*-His8* | SGB357 | pAK2E63 | this work |
| SGB473 | *W168 ΔbceSAB sacA::PbceA-luxABCDE thrC::Pxyl-bceAB-FLAG lacA::Pxyl-bceS*$^{K167D/K116E}$*-His8* | SGB357 | pAK2E66 | this work |
| SGB513 | *W168 ΔbceSAB amyE::PbceA-lacZ thrC::Pxyl-bceAB*$^{WT}$*-FLAG lacA::Pxyl-bceS*$^{E115K}$*-His8* | SGB400 | pAK2E02 | this work |
| SGB532 | *W168 ΔbceSAB sacA::PbceA-luxABCDE thrC::Pxyl-bceAB-FLAG lacA::Pxyl-bceS*$^{E115K/K167D}$*-His8* | SGB357 | pAK2E118 | this work |
| SGB562 | *W168 ΔbceSAB sacA::PbceA-luxABCDE thrC::Pxyl-bceAB-FLAG lacA::Pxyl-bceS*$^{L119C}$*-His8* | SGB357 | pAK2E134 | this work |
| SGB565 | *W168 ΔbceSAB sacA::PbceA-luxABCDE thrC::Pxyl-bceAB-FLAG lacA::Pxyl-bceS*$^{M120C}$*-His8* | SGB357 | pAK2E137 | this work |

| | | | | |
|---|---|---|---|---|
| SGB693 | *W168 ΔbceSAB sacA::PbceA-luxABCDE thrC::Pxyl-bceAB-FLAG lacA::Pxyl-bceS*$^{L67F}$*-His8* | SGB357 | pAK2E154 | this work |
| SGB696 | *W168 ΔbceSAB sacA::PbceA-luxABCDE thrC::Pxyl-bceAB-FLAG lacA::Pxyl-bceS*$^{L67G}$*-His8* | SGB357 | pAK2E157 | this work |
| SGB707 | *W168 ΔbceSAB sacA::PbceA-luxABCDE thrC::Pxyl-bceAB-FLAG lacA::Pxyl-bceS*$^{I94G}$*-His8* | SGB357 | pAK2E168 | this work |
| SGB709 | *W168 ΔbceSAB sacA::PbceA-luxABCDE thrC::Pxyl-bceAB-FLAG lacA::Pxyl-bceS*$^{I94F}$*-His8* | SGB357 | pAK2E170 | this work |
| SGB718 | *W168 bceAB::kan sacA::PbceA-luxABCDE thrC::pXT-BceAB*$^{T541C}$*-FLAG* | SGB79 | pMG759 | this work |
| SGB719 | *W168 bceAB::kan sacA::PbceA-luxABCDE thrC::pXT-BceAB*$^{S542C}$*-FLAG* | SGB79 | pMG760 | this work |
| SGB721 | *W168 bceAB::kan sacA::PbceA-luxABCDE thrC::pXT-BceAB*$^{L546A}$*-FLAG* | SGB79 | pMG762 | this work |
| SGB722 | *W168 bceAB::kan sacA::PbceA-luxABCDE thrC::pXT-BceAB*$^{L546F}$*-FLAG* | SGB79 | pMG763 | this work |
| SGB723 | *W168 bceAB::kan sacA::PbceA-luxABCDE thrC::pXT-BceAB*$^{Q550A}$*-FLAG* | SGB79 | pMG764 | this work |
| SGB724 | *W168 bceAB::kan sacA::PbceA-luxABCDE thrC::pXT-BceAB*$^{Q550F}$*-FLAG* | SGB79 | pMG765 | this work |
| SGB731 | *W168 bceAB::kan sacA::PbceA-luxABCDE thrC::pXT-BceAB-FLAG* | SGB79 | pFK727 | this work |
| SGB732 | *W168 bceAB::kan sacA::PbceA-luxABCDE thrC::pXT-BceAB*$^{F538C}$*-FLAG* | SGB79 | pMG712 | this work |
| SGB765 | *W168 bceAB::kan sacA::PbceA-luxABCDE thrC::pXT-BceAB*$^{L546N}$*-FLAG* | SGB79 | pMG770 | this work |
| SGB766 | *W168 bceAB::kan sacA::PbceA-luxABCDE thrC::pXT-BceAB*$^{Q550E}$*-FLAG* | SGB79 | pMG771 | this work |
| SGB771 | *W168 ΔbceS sacA::PbceA-luxABCDE* | TMB1036 | SGB378 | this work |
| SGB790 | *W168 ΔbceSAB sacA::PbceA-luxABCDE thrC::Pxyl-bceAB-FLAG lacA::Pxyl-bceS*$^{E115K/K116E/E118C}$*-His8* | SGB357 | pAK2E176 | this work |
| SGB791 | *W168 ΔbceSAB sacA::PbceA-luxABCDE thrC::Pxyl-bceAB-FLAG lacA::Pxyl-bceS*$^{E115K/K116E/L119C}$*-His8* | SGB357 | pAK2E177 | this work |
| SGB792 | *W168 ΔbceS sacA::PbceA-luxABCDE thrC::Pxyl-bceS*$^{WT}$ | SGB771 | pAKXT03 | this work |
| SGB795 | *W168 ΔbceS sacA::PbceA-luxABCDE thrC::Pxyl-bceS*$^{E115K}$ | SGB771 | pAKXT06 | this work |
| SGB807 | *W168 bceAB::kan sacA::PbceA-lux thrC::pXT-BceAB*$^{L546D}$*-FLAG* | SGB79 | pMG772 | this work |
| SGB818 | *W168 ΔbceSAB sacA::PbceA-luxABCDE thrC::Pxyl-bceS*$^{WT}$ | SGB378 | pAKXT03 | this work |
| SGB820 | *W168 ΔbceSAB sacA::PbceA-luxABCDE thrC::Pxyl-bceS*$^{E115K}$ | SGB378 | pAKXT06 | this work |
| SGB868 | *W168 ΔbceS sacA::PbceA-luxABCDE thrC::Pxyl-bceS*$^{E115K/K116E}$ | SGB771 | pAKXT13 | this work |
| SGB869 | *W168 ΔbceSAB sacA::PbceA-luxABCDE thrC::Pxyl-bceS*$^{E115K/K116E}$ | SGB378 | pAKXT13 | this work |
| SGB886 | *W168 ΔbceSAB sacA::PbceA-luxABCDE lacA::Pxyl-bceS*$^{L67F}$*-His8* | SGB378 | pAK2E154 | this work |
| SGB895 | *W168 ΔbceS sacA::PbceA-luxABCDE thrC::Pxyl-bceS*$^{W54R}$ | SGB771 | pAKXT15 | this work |
| SGB896 | *W168 ΔbceSAB sacA::PbceA-luxABCDE thrC::Pxyl-bceS*$^{W54R}$ | SGB378 | pAKXT15 | this work |
| SGB897 | *W168 ΔbceS sacA::PbceA-luxABCDE thrC::Pxyl-bceS*$^{F55R}$ | SGB771 | pAKXT22 | this work |
| SGB898 | *W168 ΔbceSAB sacA::PbceA-luxABCDE thrC::Pxyl-bceS*$^{F55R}$ | SGB378 | pAKXT22 | this work |
| SGB899 | *W168 ΔbceS sacA::PbceA-luxABCDE thrC::Pxyl-BceS*$^{Y57R}$ | SGB771 | pAKXT20 | this work |
| SGB900 | *W168 ΔbceSAB sacA::PbceA-luxABCDE thrC::Pxyl-bceS*$^{Y57R}$ | SGB378 | pAKXT20 | this work |
| SGB901 | *W168 ΔbceS sacA::PbceA-luxABCDE thrC::Pxyl-BceS*$^{W54R/F55R}$ | SGB771 | pAKXT16 | this work |
| SGB902 | *W168 ΔbceSAB sacA::PbceA-luxABCDE thrC::Pxyl-bceS*$^{W55R/F55R}$ | SGB378 | pAKXT16 | this work |
| SGB903 | *W168 ΔbceS sacA::PbceA-luxABCDE thrC::Pxyl-BceS*$^{F55R/Y57R}$ | SGB771 | pAKXT21 | this work |

| | | | | |
|---|---|---|---|---|
| SGB904 | *W168 ΔbceSAB sacA::PbceA-luxABCDE thrC::Pxyl-bceS*$^{F55R/Y57R}$ | SGB378 | pAKXT21 | this work |
| SGB936 | *W168 ΔbceSAB sacA::PbceA-luxABCDE thrC::Pxyl-bceAB-FLAG*<br>*lacA::Pxyl-bceS*$^{C45S/C198S/C259S}$*-His8* | SGB357 | pAK2E185 | this work |
| SGB937 | *W168 ΔbceSAB sacA::PbceA-luxABCDE thrC::Pxyl-bceAB-FLAG*<br>*lacA::Pxyl-bceS*$^{C45S/C198S/C259S/I50C}$*-His8* | SGB367 | pAK2E186 | this work |
| SGB938 | *W168 ΔbceSAB sacA::PbceA-luxABCDE thrC::Pxyl-bceAB-FLAG*<br>*lacA::Pxyl-bceS*$^{C45S/C198S/C259S/I51C}$*-His8* | SGB357 | pAK2E187 | this work |
| SGB939 | *W168 ΔbceSAB sacA::PbceA-luxABCDE thrC::Pxyl-bceAB-FLAG*<br>*lacA::Pxyl-bceS*$^{C45S/C198S/C259S/F52C}$*-His8* | SGB357 | pAK2E188 | this work |
| SGB940 | *W168 ΔbceSAB sacA::PbceA-luxABCDE thrC::Pxyl-bceAB-FLAG*<br>*lacA::Pxyl-bceS*$^{C45S/C198S/C259S/L53C}$*-His8* | SGB357 | pAK2E189 | this work |
| SGB941 | *W168 ΔbceSAB sacA::PbceA-luxABCDE thrC::Pxyl-bceAB-FLAG*<br>*lacA::Pxyl-bceS*$^{C45S/C198S/C259S/W54C}$*-His8* | SGB357 | pAK2E190 | this work |
| SGB942 | *W168 ΔbceSAB sacA::PbceA-luxABCDE thrC::Pxyl-bceAB-FLAG*<br>*lacA::Pxyl-bceS*$^{C45S/C198S/C259S/F55C}$*-His8* | SGB357 | pAK2E191 | this work |
| SGB943 | *W168 ΔbceSAB sacA::PbceA-luxABCDE thrC::Pxyl-bceAB-FLAG*<br>*lacA::Pxyl-bceS*$^{C45S/C198S/C259S/R56C}$*-His8* | SGB357 | pAK2E192 | this work |
| SGB944 | *W168 ΔbceSAB sacA::PbceA-luxABCDE thrC::Pxyl-bceAB-FLAG*<br>*lacA::Pxyl-bceS*$^{C45S/C198S/C259S/Y57C}$*-His8* | SGB357 | pAK2E193 | this work |
| SGB945 | *W168 ΔbceSAB sacA::PbceA-luxABCDE thrC::Pxyl-bceAB-FLAG*<br>*lacA::Pxyl-bceS*$^{C45S/C198S/C259S/R58C}$*-His8* | SGB357 | pAK2E194 | this work |
| SGB946 | *W168 ΔbceSAB sacA::PbceA-luxABCDE thrC::Pxyl-bceAB-FLAG*<br>*lacA::Pxyl-bceS*$^{C45S/C198S/C259S/K59C}$*-His8* | SGB357 | pAK2E195 | this work |
| SGB947 | *W168 ΔbceSAB sacA::PbceA-luxABCDE thrC::Pxyl-bceAB-FLAG*<br>*lacA::Pxyl-bceS*$^{C45S/C198S/C259S/T61C}$*-His8* | SGB357 | pAK2E196 | this work |
| SGB948 | *W168 ΔbceSAB sacA::PbceA-luxABCDE thrC::Pxyl-bceAB-FLAG*<br>*lacA::Pxyl-bceS*$^{C45S/C198S/C259S/A62C}$*-His8* | SGB357 | pAK2E197 | this work |
| SGB949 | *W168 ΔbceSAB sacA::PbceA-luxABCDE thrC::Pxyl-bceAB-FLAG*<br>*lacA::Pxyl-bceS*$^{C45S/C198S/C259S/Y64C}$*-His8* | SGB357 | pAK2E198 | this work |
| SGB950 | *W168 ΔbceSAB sacA::PbceA-luxABCDE thrC::Pxyl-bceAB-FLAG*<br>*lacA::Pxyl-bceS*$^{C45S/C198S/C259S/K65C}$*-His8* | SGB357 | pAK2E199 | this work |
| SGB951 | *W168 ΔbceSAB sacA::PbceA-luxABCDE thrC::Pxyl-bceAB-FLAG*<br>*lacA::Pxyl-bceS*$^{E115K/K116E/C45S/C198S/C259S}$*-His8* | SGB357 | pAK2E202 | this work |
| SGB952 | *W168 ΔbceSAB sacA::PbceA-luxABCDE thrC::Pxyl-bceAB-FLAG*<br>*lacA::Pxyl-bceS*$^{E115K/K116E/C45S/C198S/C259S/I50C}$*-His8* | SGB357 | pAK2E203 | this work |
| SGB953 | *W168 ΔbceSAB sacA::PbceA-luxABCDE thrC::Pxyl-bceAB-FLAG*<br>*lacA::Pxyl-bceS*$^{E115K/K116E/C45S/C198S/C259S/I51C}$*-His8* | SGB357 | pAK2E204 | this work |
| SGB954 | *W168 ΔbceSAB sacA::PbceA-luxABCDE thrC::Pxyl-bceAB-FLAG*<br>*lacA::Pxyl-bceS*$^{E115K/K116E/C45S/C198S/C259S/F52C}$*-His8* | SGB357 | pAK2E205 | this work |
| SGB955 | *W168 ΔbceSAB sacA::PbceA-luxABCDE thrC::Pxyl-bceAB-FLAG*<br>*lacA::Pxyl-bceS*$^{E115K/K116E/C45S/C198S/C259S/L53C}$*-His8* | SGB357 | pAK2E206 | this work |
| SGB956 | *W168 ΔbceSAB sacA::PbceA-luxABCDE thrC::Pxyl-bceAB-FLAG*<br>*lacA::Pxyl-bceS*$^{E115K/K116E/C45S/C198S/C259S/W54C}$*-His8* | SGB357 | pAK2E207 | this work |
| SGB957 | *W168 ΔbceSAB sacA::PbceA-luxABCDE thrC::Pxyl-bceAB-FLAG*<br>*lacA::Pxyl-bceS*$^{E115K/K116E/C45S/C198S/C259S/F55C}$*-His8* | SGB357 | pAK2E208 | this work |
| SGB958 | *W168 ΔbceSAB sacA::PbceA-luxABCDE thrC::Pxyl-bceAB-FLAG*<br>*lacA::Pxyl-bceS*$^{E115K/K116E/C45S/C198S/C259S/R56C}$*-His8* | SGB357 | pAK2E209 | this work |
| SGB959 | *W168 ΔbceSAB sacA::PbceA-luxABCDE thrC::Pxyl-bceAB-FLAG*<br>*lacA::Pxyl-bceS*$^{E115K/K116E/C45S/C198S/C259S/Y57C}$*-His8* | SGB357 | pAK2E210 | this work |
| SGB960 | *W168 ΔbceSAB sacA::PbceA-luxABCDE thrC::Pxyl-bceAB-FLAG*<br>*lacA::Pxyl-bceS*$^{E115K/K116E/C45S/C198S/C259S/R58C}$*-His8* | SGB357 | pAK2E211 | this work |
| SGB961 | *W168 ΔbceSAB sacA::PbceA-luxABCDE thrC::Pxyl-bceAB-FLAG*<br>*lacA::Pxyl-bceS*$^{E115K/K116E/C45S/C198S/C259S/K59C}$*-His8* | SGB357 | pAK2E212 | this work |
| SGB962 | *W168 ΔbceSAB sacA::PbceA-luxABCDE thrC::Pxyl-bceAB-FLAG*<br>*lacA::Pxyl-bceS*$^{E115K/K116E/C45S/C198S/C259S/T61C}$*-His8* | SGB357 | pAK2E213 | this work |
| SGB963 | *W168 ΔbceSAB sacA::PbceA-luxABCDE thrC::Pxyl-bceAB-FLAG*<br>*lacA::Pxyl-bceS*$^{E115K/K116E/C45S/C198S/C259S/A62C}$*-His8* | SGB357 | pAK2E214 | this work |
| SGB964 | *W168 ΔbceSAB sacA::PbceA-luxABCDE thrC::Pxyl-bceAB-FLAG*<br>*lacA::Pxyl-bceS*$^{E115K/K116E/C45S/C198S/C259S/Y64C}$*-His8* | SGB357 | pAK2E215 | this work |

| | | | | |
|---|---|---|---|---|
| SGB965 | *W168 ΔbceSAB sacA::PbceA-luxABCDE thrC::Pxyl-bceAB-FLAG lacA::Pxyl-bceS*$^{E115K/K116E/C45S/C198S/C259S/K65C}$*-His8* | SGB357 | pAK2E216 | this work |
| SGB976 | *W168 bceAB::kan sacA::PbceA-luxABCDE thrC::pXT-BceAB*$^{F533C}$*-FLAG* | SGB79 | pMG775 | this work |
| SGB977 | *W168 bceAB::kan sacA::PbceA-luxABCDE thrC::pXT-BceAB*$^{L534C}$*-FLAG* | SGB79 | pMG776 | this work |
| SGB978 | *W168 bceAB::kan sacA::PbceA-luxABCDE thrC::pXT-BceAB*$^{G535C}$*-FLAG* | SGB79 | pMG777 | this work |
| SGB979 | *W168 bceAB::kan sacA::PbceA-luxABCDE thrC::pXT-BceAB*$^{L536C}$*-FLAG* | SGB79 | pMG778 | this work |
| SGB980 | *W168 bceAB::kan sacA::PbceA-luxABCDE thrC::pXT-BceAB*$^{T537C}$*-FLAG* | SGB79 | pMG779 | this work |
| SGB981 | *W168 bceAB::kan sacA::PbceA-luxABCDE thrC::pXT-BceAB*$^{L539C}$*-FLAG* | SGB79 | pMG780 | this work |
| SGB982 | *W168 bceAB::kan sacA::PbceA-luxABCDE thrC::pXT-BceAB*$^{I540C}$*-FLAG* | SGB79 | pMG781 | this work |
| TMB035 | *W168 bceAB::kan* | | | (Rietkötter *et al.*, 2008) |
| TMB1036 | *trpC2 ΔbceS* | | | gift from Mascher lab |
| W168 | Laboratory (wild-type) strain of *Bacillus subtilis*; *trpC2* | | | laboratory stock |

**[a]** Parent strain indicates the previous step in the genetic construction of the strain.

**[b]** Plasmids or genomic DNA of the indicated strain used in the last transformation step of genetic construction of the strain.

### Section B - Primers[a]

| Primer | Sequence (5'→3') |
|---|---|
| SG0068 | CCTTTATTTTTGTCAAATGGGTGAAAGTGAAGATGAAAAACCGAG |
| SG0069 | CACCCATTTGACAAAAATAAAGGATACAACCTGATGTAATCAGG |
| SG0109 | TGAAAGTGAATGTGAAAAACCGAGCTATACAATTTTAAGAAAACTCGG |
| SG0110 | TCGGTTTTTCACATTCACTTTCACCCATTTGTTTAAAATAAAGGATAC |
| SG0117 | AGGTTGTATCTGTTATTTTAAACAAATGGGTGAAAGTGAAGATGAAAAACC |
| SG0118 | GTTTAAAATAACAGATACAACCTGATGTAATCAGGAACGTTAACCC |
| SG0119 | ACAAATGGGTTGTAGTGAAGATGAAAAACCGAGCTATACAATTTTAAG |
| SG0120 | CATCTTCACTACAACCCATTTGTTTAAAATAAAGGATACAACCTGATG |
| SG0127 | CATCGCTTAGCATTGGAAAATAAAAAAGATG |
| SG0128 | CCATGCCATCAGCTCATCTTTTTTATTTTC |
| SG0175 | GTATCCTTTGTTTTAAACAAATGGGTGAAAGTGAAGATGAAAAACCGAGC |
| SG0176 | CATTTGTTTAAAACAAAGGATACAACCTGATGTAATCAGGAACGTTAACCC |
| SG0179 | AATTTAATTTCCATGGCTTGTCAACCTGGCCGTTCCGATG |
| SG0180 | AATTTAATTTGAATTCCTGCTGAAACAAAAACGCGGCAAT |
| SG0181 | AATTTAATTTGAATTCCTGTCAGTTCTTTATTATAAGAAG |
| SG0182 | AATTTAATTTGGATCCGAAATGGTATTAGCATTCTTAGGT |
| SG0197 | GGGTTAACGTGCCTGATTACATCAGGTTGTATCCTTTATTTTAAAC |
| SG0198 | GTAATCAGGCACGTTAACCCTAGAAGCCGACGATG |
| SG0215 | CGGCATCGCTTAGCATTGGAAAATGCTAAAGATGAG |
| SG0216 | GATCCATGCCATCAGCTCATCTTTAGCATTTTCC |
| SG0217 | CGGCATCGCTTAGCATTGGAAAATCGTAAAGATGAG |
| SG0218 | GATCCATGCCATCAGCTCATCTTTACGATTTTCC |
| SG0219 | GGCATCGCTTAGCATTGGAATGCGAAAAAG |
| SG0220 | GCCATCAGCTCATCTTTTTCGCATTCCAATG |
| SG0221 | CATCGCTTAGCATTGGAAAATTGCAAAGATG |
| SG0222 | CCATGCCATCAGCTCATCTTTGCAATTTTC |
| SG0223 | GCTTAGCATTGGAAAATGAATGCGATGAG |
| SG0224 | GATCCATGCCATCAGCTCATCGCATTCAT |
| SG0225 | CTTCTTGATCAGCAGCTTCATTGCAAACG |
| SG0226 | GTTTTCAATAAATGAAATGCGTTTGCAATGAAG |
| SG0227 | CTTGATCAGCAGCTTCATCAATGCCGCATTTC |
| SG0228 | GTCGTTTTCAATAAATGAAATGCGGCATTGATG |
| SG0229 | GATCAGCAGCTTCATCAAAAATGCATTTC |
| SG0230 | GGTCGTTTTCAATAAATGAAATGCATTTTTG |
| SG0231 | CATCATGTTGTATCCTTTATTTTAAACAAATGGGTG |
| SG0232 | GATACAACATGATGTAATCAGGAACGTTAACC |
| SG0233 | CAGGTTGTTGCCTTTATTTTAAACAAATGGGTG |
| SG0234 | AAATAAAGGCAACAACCTGATGTAATCAGG |
| SG0235 | CTTTATTGTAAACAAATGGGTGAAAGTGAAGATG |
| SG0236 | GTTTACAATAAAGGATACAACCTGATGTAATCAG |
| SG0237 | TTGTATCCTTTATTTTAAATGCATGGGTG |
| SG0238 | CATCTTCACTTTCACCCATGCATTTAA |

| | |
|---|---|
| SG0239 | GGTTGTATCCTTTATTTTAAACAATGCGGTGAAAG |
| SG0240 | CGGTTTTTCATCTTCACTTTCACCGCATTGTT |
| SG0241 | CAAATGTGTGAAAGTGAAGATGAAAAACCGAGC |
| SG0242 | CTTTCACACATTTGTTTAAAATAAAGGATACAACC |
| SG0243 | CAGGTAGTATCCTTTATTTTAAACAAATGGG |
| SG0244 | GGATACTACCTGATGTAATCAGGAACG |
| SG0245 | GGCAACCGAGCGTTCTG |
| SG0246 | CTGACAGCGTTTCGATCC |
| SG0247 | CATATTCTGTTGGTTCCGCTATCGGAAAGAAAC |
| SG0248 | CGGAACCAACAGAATATGATAAAAAACAAGATGCAC |
| SG0249 | GTGCATCTTGTTTTTTATCATATTCCTTTGCTTCCGC |
| SG0250 | GCTGTTTCTTTCCGATAGCGGAAGCAAAGG |
| SG0251 | TTTGGTGCCGCTATCGGAAAGAAACAGCGTTTTATAAAAGC |
| SG0252 | CCGATAGCGGCACCAAAGGAATATGATAAAAAACAAGATGC |
| SG0253 | TGGTTCTGCTATCGGAAAGAAACAGC |
| SG0254 | CCGATAGCAGAACCAAAGGAATATGATAAA |
| SG0255 | CGCTGTCGGAAAGAAACAGCGTTTTATAAAAGC |
| SG0256 | CTTTCCGACAGCGGAACCAAAGGAATATGATAAA |
| SG0257 | CCGCTATTGTAAAGAAACAGCGTTTTATAAAAGC |
| SG0258 | GTTTCTTTACAATAGCGGAACCAAAGGAATATG |
| SG0259 | TATCGGTGCGAAACAGCGTTTTATAAAAGCTTGAAAACATGG |
| SG0260 | CGCTGTTTCACAGCGATAGCAGAACCAAAGG |
| SG0261 | GGAAATGCACAGCGTTTTATAAAAGCTTGAAAACATGGG |
| SG0262 | GCTGTGCATTTCCGATAGCGGAACCAAAGG |
| SG0263 | CGGAAAGAATGCGCGTTTTATAAAAGCTTGAAAACATGG |
| SG0264 | AACGCGCATTCTTTGCGATAGCAGAACCAAAGG |
| SG0265 | GAAACATGCTTTTATAAAAGCTTGAAAACATGGGAGAAC |
| SG0266 | TAAAAGCATGTTTCTTTCCGATAGCGGAACCAAAG |
| SG0267 | GCTTAGCATTGGAAAATGAAGAAGATGAG |
| SG0268 | GATCCATGCCATCAGCTCATCTTCTTCAT |
| SG0275 | CTTGATCAGCAGCTTCATCAAGACCGCATTTC |
| SG0276 | GTCGTTTTCAATAAATGAAATGCGGTCTTGATG |
| SG0283 | CTTAGCATTGGAAAATGAAAAATGCGAGCTG |
| SG0284 | GGATCCATGCCATCAGCTCGCATTTTTC |
| SG0285 | GCATTGGAAAATGAAAAAGATTGCCTGATG |
| SG0286 | CTCATGGATCCATGCCATCAGGCAATCTTT |
| SG0289 | CGCTTAGCATTGGAAAATAAAGAAGATGAG |
| SG0290 | GATCCATGCCATCAGCTCATCTTCTTTATTTTC |
| SG0295 | GAAGCAATGGTTGAAAGAAGCTGCGCCGGG |
| SG0296 | CAAGTGTTCTGTTTGCCCGGCGCAGCTTC |
| SG0297 | GCAATGGTTGAAAGAAGCATTTGCGGGCAA |
| SG0298 | CTTCAAGTGTTCTGTTTGCCCGCAAATGC |
| SG0299 | CAATGGTTGAAAGAAGCATTGCCTGCCAAAC |
| SG0300 | CTGCTTCAAGTGTTCTGTTTGGCAGGCAATG |

| | |
|---|---|
| SG0301 | GTTGAAAGAAGCATTGCCGGGTGCACAGAAC |
| SG0302 | GTCTGCTTCAAGTGTTCTGTGCACCCGGC |
| SG0303 | GAAAGAAGCATTGCCGGGCAATGCGAACAC |
| SG0304 | GCGGTCTGCTTCAAGTGTTCGCATTGCCC |
| SG0305 | GAAGCATTGCCGGGCAAACATGCCACTTG |
| SG0306 | GCTGCGGTCTGCTTCAAGTGGCATGTTTG |
| SG0307 | GCATTGCCGGGCAAACAGAATGCTTGAAG |
| SG0308 | CGTGCTGCGGTCTGCTTCAAGCATTCTG |
| SG0309 | CATTGCCGGGCAAACAGAACACTGCAAGCAG |
| SG0310 | GCCGTGCTGCGGTCTGCTTGCAGTGTTC |
| SG0311 | CCGGGCAAACAGAACACTTGTGCCAGACC |
| SG0312 | GATGCCGTGCTGCGGTCTGGCACAAGTG |
| SG0313 | GGGCAAACAGAACACTTGAAGTGCACCGC |
| SG0314 | TAAGCGATGCCGTGCTGCGGTGCACTTCAA |
| SG0315 | GAACCGGAAACGCCGTTTTGCGCAATG |
| SG0316 | CAATGCTTCTTTCAACCATTGCGCAAAACGG |
| SG0317 | GAACCGGAAACGCCGTTTGAATGCATGGTTG |
| SG0318 | CGGCAATGCTTCTTTCAACCATGCATTCAAAC |
| SG0319 | CCGGAAACGCCGTTTGAAGCATGCGTTGAA |
| SG0320 | CCCGGCAATGCTTCTTTCAACGCATGCTTC |
| SG0321 | GAAACGCCGTTTGAAGCAATGTGCGAAAG |
| SG0322 | GCCCGGCAATGCTTCTTTCGCACATTGC |
| SG0323 | ACGCCGTTTGAAGCAATGGTTTGCAGAAGC |
| SG0324 | CTGTTTGCCCGGCAATGCTTCTGCAAACCAT |
| SG0325 | CCGTTTGAAGCAATGGTTGAATGCAGCATTG |
| SG0326 | GTTCTGTTTGCCCGGCAATGCTGCATTCAAC |
| SG0327 | GTTTGAAGCAATGGTTGAAAGATGCATTGCCG |
| SG0328 | GTGTTCTGTTTGCCCGGCAATGCATCTTTC |
| SG0329 | CGCTATCGGAAAGAAACAGCGTGCTATAAAAG |
| SG0330 | CTCCCATGTTTTCAAGCTTTTATAGCACGCTGT |
| SG0331 | GCTATCGGAAAGAAACAGCGTTTTGCAAAAGC |
| SG0332 | CTCCCATGTTTTCAAGCTTTTGCAAAACGC |
| SG0333 | CGGAAAGAAACAGCGTTTTATTGCAGCTTG |
| SG0334 | GTTCTCCCATGTTTTCAAGCTGCAATAAAAC |
| SG0335 | TCGGAAAGAAACAGCGTTTTATAAATGCTTGAAAAC |
| SG0336 | GAGATTGTTCTCCCATGTTTTCAAGCATTTATA |
| SG0337 | GAAACAGCGTTTTATAAAAGCTGCAAAACATG |
| SG0338 | GAGATTGTTCTCCCATGTTTTGCAGCTTTTA |
| SG0339 | GAAACAGCGTTTTATAAAAGCTTGTGCACATGG |
| SG0340 | CATCGAGATTGTTCTCCCATGTGCACAAGCT |
| SG0341 | GCGTTTTATAAAAGCTTGAAATGCTGGGAGA |
| SG0342 | CACATCGAGATTGTTCTCCCAGCATTTCAAG |
| SG0343 | GCGTTTTATAAAAGCTTGAAAACATGCGAGAAC |
| SG0344 | GTCACATCGAGATTGTTCTCGCATGTTTTC |

| | |
|---|---|
| SG0356 | GGTGAATGTGAAGATGAAAAACCGAGCTATAC |
| SG0357 | CATCTTCACATTCACCCATTTGTTTAAAATAAAGGATAC |
| SG0358 | GAAAGTTGCGATGAAAAACCGAGCTATACAATTTTAAG |
| SG0359 | TTCATCGCAACTTTCACCCATTTGTTTAAAATAAAGGATAC |
| SG0400 | CAGCGTGTTATAAAAGCTTGAAAACATGGGAGAACAATCTCG |
| SG0401 | CTTTTATAACACGCTGTTTCTTTCCGATAGCGGAACCAAAG |
| SG0402 | GCGTTTTGTAAAAGCTTGAAAACATGGGAGAACAATCTCGATGTGAC |
| SG0403 | GCTTTTACAAAACGCTGTTTCTTTCCGATAGCGGAACCAAAGG |
| SG0404 | GTTTTATTGTAGCTTGAAAACATGGGAGAACAATCTCGATGTGACAGC |
| SG0405 | CAAGCTACAATAAAACGCTGTTTCTTTCCGATAGCGGAACC |
| SG0480 | TGAGTGCATGGCATGGATCCATGAGGTC |
| SG0481 | GCCATGCACTCATCTTTTTCATTTTCC |
| SG0486 | GCTGTGCGCATGGATCCATGAGGTCAAA |
| SG0487 | ATGCGCACAGCTCATCTTTTTCATTTTC |
| SG0534 | AAGCTTCAAAACATGGGAGAACAATCTCGATG |
| SG0535 | GTTTTGAAGCTTTTATAAAACGCTGTTTCTTTC |
| SG0542 | AAGCGGTAAAACATGGGAGAACAATCTCGATG |
| SG0543 | GTTTTACCGCTTTTATAAAACGCTGTTTCTTTC |
| SG0579 | GAAGCGGTGCCGGGCAAACAGAACACTTGAAG |
| SG0580 | CCGGCACCGCTTCTTTCAACCATTGCTTCAAAC |
| SG0583 | GAAGCTTCGCCGGGCAAACAGAACACTTGAAG |
| SG0584 | CCGGCGAAGCTTCTTTCAACCATTGCTTCAAAC |
| SG0614 | GTATCGCTTATTTTAAACAAATGGGTG |
| SG0615 | AATAAGCGATACAACCTGATGTAATCAG |
| SG0616 | GTATCTTTTATTTTAAACAAATGGGTG |
| SG0617 | AATAAAAGATACAACCTGATGTAATCAG |
| SG0618 | TTGTATCCTTTATTTTAAAGCAATGGGTG |
| SG0619 | CATCTTCACTTTCACCCATTGCTTTAA |
| SG0620 | TTGTATCCTTTATTTTAAATTTATGGGTG |
| SG0621 | CATCTTCACTTTCACCCATAAATTTAA |
| SG0622 | GATTACATGCGGTTGTATCCTTTATTTTAA |
| SG0623 | CAACCGCATGTAATCAGGAACGTTA |
| SG0624 | CTGATTTGTTCAGGTTGTATCCTTTA |
| SG0625 | ACCTGAACAAATCAGGAACGTTAAC |
| SG0659 | GAAAATAAAGAAGATTGCCTGATGGCATGGATCCATGAG |
| SG0660 | CATCAGGCAATCTTCTTTATTTTCCAATGCTAAGCGATG |
| SG0661 | GAAAATAAAGAAGATGAGTGCATGGCATGGATCCATGAGGTC |
| SG0662 | TGCCATGCACTCATCTTCTTTATTTTCCAATGCTAAGCGATG |
| SG0671 | AGTCGGTCTCGGATCCTACATCGCGAAGGAAGAGG |
| SG0672 | AGTCGGTCTCGAATTCGGGCTTTTCCTTCGATACGG |
| SG0675 | GTATCAATTATTTTAAACAAATGGGTG |
| SG0676 | AATAATTGATACAACCTGATGTAATCAG |
| SG0677 | TTGTATCCTTTATTTTAAAGAAATGGGTG |
| SG0678 | CATCTTCACTTTCACCCATTTCTTTAA |

| | |
|---|---|
| SG0715 | GTATCGATTATTTTAAACAAATGGGTG |
| SG0716 | AATAATCGATACAACCTGATGTAATCAG |
| SG0727 | TTTTTGCATATTCCTTTGGTTCCGCTAT |
| SG0729 | TATCTGCTTCCTTTGGTTCCGCTATCGG |
| SG0731 | CATATGCCTTTGGTTCCGCTATCGGAAA |
| SG0733 | ATTCTGCTGGTTCCGCTATCGGAAAGAA |
| SG0734 | ACCAGCAGAATATGATAAAAAACAAGAT |
| SG0735 | CCTTTGCTTCCGCTATCGGAAAGAAACA |
| SG0736 | GGAAGCAAAGGAATATGATAAAAAACAA |
| SG0737 | TTGGTGCCGCTATCGGAAAGAAACAGCG |
| SG0738 | AGCGGCACCAAAGGAATATGATAAAAAA |
| SG0739 | GTTCTGCTATCGGAAAGAAACAGCGTTT |
| SG0740 | GATAGCAGAACCAAAGGAATATGATAAA |
| SG0741 | CCGCTGCCGGAAAGAAACAGCGTTTTAT |
| SG0742 | TCCGGCAGCGGAACCAAAGGAATATGAT |
| SG0743 | CTATTGCAAAGAAACAGCGTTTTATAAA |
| SG0744 | CTTTGCAATAGCGGAACCAAAGGAATAT |
| SG0745 | TCGGTGCGAAACAGCGTTTTATAAAAGC |
| SG0746 | TTTCGCACCGATAGCGGAACCAAAGGAA |
| SG0779 | GTGGTCTATCCAAAAAGGGATCGGTTTT |
| SG0780 | GGATAGACCACGATTGTAAATCTTTGAT |
| SG0781 | AGATTCTGGCAGGGGCATTGATCCAAAA |
| SG0782 | TGCCAGAATCTTTCACTTGGAGCTGCGT |
| SG0783 | ATATGCAAAAAAACAAGATAGACAAATA |
| SG0784 | GGAAGCAGATAAAAAACAAGATAGACAA |
| SG0785 | AAAGGCATATGATAAAAAACAAGATAGA |
| SG0806 | AAATAAAGAAGATGAGCTGATGGCATGGATCCATGA |
| SG0807 | CATCTTCTTTATTTTCCAATGCTAAGCGATGCCGTG |
| SG0812 | CCTTCGTTTCCGCTATCGGAAAGAAACAGCGTT |
| SG0813 | GGAAACGAAGGAATATGATAAAAAACAAGATGC |
| SG0814 | TTGGCGTCGCTATCGGAAAGAAACAGCGTTTTA |
| SG0815 | AGCGACGCCAAAGGAATATGATAAAAAACAAGA |
| SG0816 | CCTTCGTCGTCGCTATCGGAAAGAAACAGCGTTTTA |
| SG0817 | AGCGACGACGAAGGAATATGATAAAAAACAAGATGC |
| SG0828 | TTTGTCTATCTTGTTTTTTATCATATTCCTTTG |
| SG0829 | AGATAGACAAATACACCATATAAAGTACATTCC |
| SG0897 | TCCGCCGTCGGAAAGAAACAGCGTTTTAT |
| SG0898 | TCCGACGGCGGAACCAAAGGAATATGAT |
| SG899 | TTGGCGTCGCCGTCGGAAAGAAACAGCGTTTTAT |
| SG0900 | TCCGACGGCGACGCCAAAGGAATATGATAAAAAAC |
| SG0932 | CGTCGGCTGCTTAGGGTTAACGTTCCTG |
| SG0933 | CCCTAAGCAGCCGACGATGAACATCACC |
| SG0934 | CGGCTTCTGTGGGTTAACGTTCCTGATTAC |
| SG0935 | GTTAACCCACAGAAGCCGACGATGAACATCACC |

| | |
|---|---|
| SG0936 | GCTTCTTATGTTTAACGTTCCTGATTACATC |
| SG0937 | GAACGTTAAACATAAGAAGCCGACGATGAAC |
| SG0938 | CTTAGGGTGCACGTTCCTGATTACATCAGG |
| SG0939 | GGAACGTGCACCCTAAGAAGCCGACG |
| SG0940 | GGGTTATGCTTCCTGATTACATCAGGTTG |
| SG0941 | CAGGAAGCATAACCCTAAGAAGCCGACG |
| SG0942 | TAACGTTCTGCATTACATCAGGTTGTATCCTT |
| SG0943 | GATGTAATGCAGAACGTTAACCCTAAGAAGC |
| SG0944 | CGTTCCTGTGTACATCAGGTTGTATCCTTTAT |
| SG0945 | CCTGATGTACACAGGAACGTTAACCCTAAG |

[a] See section C for use of each primer

### Section C - Plasmids

| Plasmid | Description[a] | Parent plasmid | Plasmid construction[b,c] | Reference |
|---|---|---|---|---|
| pAK101 | *bla 'ytrF' bgaB erm* | pMAD | SG0179/SG0180 (W168 gDNA) | this work |
| pAK102 | *bla 'ytrF bgaB erm yttA'* | pMAD | SG0181/SG0182 (W168 gDNA) | this work |
| pAK2E02 | *bla lacA*::($P_{xyl}$-*bceS*$^{E115K}$-*His8*) *mls* | pSD2E01 | SG0127/SG0128 (QC) | this work |
| pAK2E12 | *bla lacA*::($P_{xyl}$-*bceS*$^{E115A}$-*His8*) *mls* | pSD2E01 | SG0215/SG0216 (QC) | this work |
| pAK2E13 | *bla lacA*::($P_{xyl}$-*bceS*$^{E115R}$-*His8*) *mls* | pSD2E01 | SG0217/SG0218 (QC) | this work |
| pAK2E14 | *bla lacA*::($P_{xyl}$-*bceS*$^{E115C}$-*His8*) *mls* | pSD2E01 | SG0221/SG0222 (QC) | this work |
| pAK2E15 | *bla lacA*::($P_{xyl}$-*bceS*$^{Q166C}$-*His8*) *mls* | pSD2E01 | SG0225/SG0226 (QC) | this work |
| pAK2E16 | *bla lacA*::($P_{xyl}$-*bceS*$^{R168C}$-*His8*) *mls* | pSD2E01 | SG0229/SG0230 (QC) | this work |
| pAK2E17 | *bla lacA*::($P_{xyl}$-*bceS*$^{K116C}$-*His8*) *mls* | pSD2E01 | SG0223/SG0224 (QC) | this work |
| pAK2E21 | *bla lacA*::($P_{xyl}$-*bceS*$^{N114C}$-*His8*) *mls* | pSD2E01 | SG0245/SG0220 & SG0246/SG0219 | this work |
| pAK2E22 | *bla lacA*::($P_{xyl}$-*bceS*$^{K167C}$-*His8*) *mls* | pSD2E01 | SG0245/SG0228 & SG0246/SG0227 | this work |
| pAK2E26 | *bla lacA*::($P_{xyl}$-*bceS*$^{K116E}$-*His8*) *mls* | pSD2E01 | SG0245/SG0268 & SG0246/SG0267 | this work |
| pAK2E29 | *bla lacA*::($P_{xyl}$-*bceS*$^{K167D}$-*His8*) *mls* | pSD2E01 | SG0245/SG0276 & SG0246/SG0275 | this work |
| pAK2E33 | *bla lacA*::($P_{xyl}$-*bceS*$^{E118C}$-*His8*) *mls* | pSD2E01 | SG0245/SG0286 & SG0246/SG0285 | this work |
| pAK2E34 | *bla lacA*::($P_{xyl}$-*bceS*$^{E115K/K116E}$-*His8*) *mls* | pSD2E02 | SG0245/SG0290 & SG0246/SG0289 | this work |
| pAK2E37 | *bla lacA*::($P_{xyl}$-*bceS*$^{I94C}$-*His8*) *mls* | pSD2E01 | SG0245/SG0296 & SG0246/SG0295 | this work |
| pAK2E38 | *bla lacA*::($P_{xyl}$-*bceS*$^{A95C}$-*His8*) *mls* | pSD2E01 | SG0245/SG0298 & SG0246/SG0297 | this work |
| pAK2E39 | *bla lacA*::($P_{xyl}$-*bceS*$^{G96C}$-*His8*) *mls* | pSD2E01 | SG0245/SG0300 & SG0246/SG0299 | this work |
| pAK2E40 | *bla lacA*::($P_{xyl}$-*bceS*$^{Q97C}$-*His8*) *mls* | pSD2E01 | SG0245/SG0302 & SG0246/SG0301 | this work |
| pAK2E41 | *bla lacA*::($P_{xyl}$-*bceS*$^{T98C}$-*His8*) *mls* | pSD2E01 | SG0245/SG0304 & SG0246/SG0303 | this work |
| pAK2E42 | *bla lacA*::($P_{xyl}$-*bceS*$^{E99C}$-*His8*) *mls* | pSD2E01 | SG0245/SG0306 & SG0246/SG0305 | this work |
| pAK2E43 | *bla lacA*::($P_{xyl}$-*bceS*$^{H100C}$-*His8*) *mls* | pSD2E01 | SG0245/SG0308 & SG0246/SG0307 | this work |
| pAK2E44 | *bla lacA*::($P_{xyl}$-*bceS*$^{L101C}$-*His8*) *mls* | pSD2E01 | SG0245/SG0310 & SG0246/SG0309 | this work |
| pAK2E45 | *bla lacA*::($P_{xyl}$-*bceS*$^{K102C}$-*His8*) *mls* | pSD2E01 | SG0245/SG0312 & SG0246/SG0311 | this work |
| pAK2E46 | *bla lacA*::($P_{xyl}$-*bceS*$^{Q103C}$-*His8*) *mls* | pSD2E01 | SG0245/SG0314 & SG0246/SG0313 | this work |
| pAK2E47 | *bla lacA*::($P_{xyl}$-*bceS*$^{E87C}$-*His8*) *mls* | pSD2E01 | SG0245/SG0316 & SG0246/SG0315 | this work |
| pAK2E48 | *bla lacA*::($P_{xyl}$-*bceS*$^{A88C}$-*His8*) *mls* | pSD2E01 | SG0245/SG0318 & SG0246/SG0317 | this work |
| pAK2E49 | *bla lacA*::($P_{xyl}$-*bceS*$^{M89C}$-*His8*) *mls* | pSD2E01 | SG0245/SG0320 & SG0246/SG0319 | this work |
| pAK2E50 | *bla lacA*::($P_{xyl}$-*bceS*$^{V90C}$-*His8*) *mls* | pSD2E01 | SG0245/SG0322 & SG0246/SG0321 | this work |
| pAK2E51 | *bla lacA*::($P_{xyl}$-*bceS*$^{E91C}$-*His8*) *mls* | pSD2E01 | SG0245/SG0324 & SG0246/SG0323 | this work |
| pAK2E52 | *bla lacA*::($P_{xyl}$-*bceS*$^{R92C}$-*His8*) *mls* | pSD2E01 | SG0245/SG0326 & SG0246/SG0325 | this work |
| pAK2E53 | *bla lacA*::($P_{xyl}$-*bceS*$^{S93C}$-*His8*) *mls* | pSD2E01 | SG0245/SG0328 & SG0246/SG0327 | this work |
| pAK2E54 | *bla lacA*::($P_{xyl}$-*bceS*$^{F63C}$-*His8*) *mls* | pBS2E01 | SG0245/SG0330 & SG0246/SG0329 | this work |
| pAK2E55 | *bla lacA*::($P_{xyl}$-*bceS*$^{Y64C}$-*His8*) *mls* | pBS2E01 | SG0245/SG0332 & SG0246/SG0331 | this work |
| pAK2E56 | *bla lacA*::($P_{xyl}$-*bceS*$^{K65C}$-*His8*) *mls* | pBS2E01 | SG0245/SG0334 & SG0246/SG0333 | this work |
| pAK2E57 | *bla lacA*::($P_{xyl}$-*bceS*$^{S66C}$-*His8*) *mls* | pBS2E01 | SG0245/SG0336 & SG0246/SG0335 | this work |
| pAK2E58 | *bla lacA*::($P_{xyl}$-*bceS*$^{L67C}$-*His8*) *mls* | pSD2E01 | SG0245/SG0338 & SG0246/SG0337 | this work |
| pAK2E59 | *bla lacA*::($P_{xyl}$-*bceS*$^{K68C}$-*His8*) *mls* | pBS2E01 | SG0245/SG0340 & SG0246/SG0339 | this work |
| pAK2E60 | *bla lacA*::($P_{xyl}$-*bceS*$^{T69C}$-*His8*) *mls* | pBS2E01 | SG0245/SG0342 & SG0246/SG0341 | this work |
| pAK2E61 | *bla lacA*::($P_{xyl}$-*bceS*$^{W70C}$-*His8*) *mls* | pBS2E01 | SG0245/SG0344 & SG0246/SG0343 | this work |
| pAK2E63 | *bla lacA*::($P_{xyl}$-*bceS*$^{D117C}$-*His8*) *mls* | pSD2E01 | SG0245/SG0284 & SG0246/SG0283 | this work |
| pAK2E66 | *bla lacA*::($P_{xyl}$-*bceS*$^{K167D/K116E}$-*His8*) *mls* | pAK2E26 | SG0245/SG0276 & SG0246/SG0275 | this work |

| | | | | |
|---|---|---|---|---|
| pAK2E118 | *bla lacA*::(P$_{xyl}$-*bceS*$^{E115K/K167D}$-*His8*) *mls* | pAK2E29 | SG0245/SG0128 & SG0246/SG0127 | this work |
| pAK2E134 | *bla lacA*::(P$_{xyl}$-*bceS*$^{D117C}$-*His8*) *mls* | pSD2E01 | SG0480/SG0481(QC) | this work |
| pAK2E137 | *bla lacA*::(P$_{xyl}$-*bceS*$^{M120C}$-*His8*) *mls* | pSD2E01 | SG0486/SG0487 (QC) | this work |
| pAK2E154 | *bla lacA*::(P$_{xyl}$-*bceS*$^{L67F}$-*His8*) *mls* | pSD2E01 | SG0534/SG0535 (QC) | this work |
| pAK2E157 | *bla lacA*::(P$_{xyl}$-*bceS*$^{L67G}$-*His8*) *mls* | pSD2E01 | SG0542/SG0543 (QC) | this work |
| pAK2E168 | *bla lacA*::(P$_{xyl}$-*bceS*$^{I94G}$-*His8*) *mls* | pSD2E01 | SG0579/SG0580 (QC) | this work |
| pAK2E170 | *bla lacA*::(P$_{xyl}$-*bceS*$^{I94F}$-*His8*) *mls* | pSD2E01 | SG0583/SG0584 (QC) | this work |
| pAK2E176 | *bla lacA*::(P$_{xyl}$-*bceS*$^{E115K/K116E/E118C}$-*His8*) *mls* | pSD2E01 | SG0659/SG0660 (QC) | this work |
| pAK2E177 | *bla lacA*::(P$_{xyl}$-*bceS*$^{E115K/K116E/L119C}$-*His8*) *mls* | pSD2E01 | SG0661/SG0662 (QC) | this work |
| pAK2E183 | *bla lacA*::(P$_{xyl}$-*bceS*$^{C45S}$-*His8*) *mls* | pSD2E01 | SG0245/SG0829 & SG0246/SG0828 | this work |
| pAK2E184 | *bla lacA*::(P$_{xyl}$-*bceS*$^{C45S/C198S}$-*His8*) *mls* | pAK2E183 | SG0245/SG0780 & SG0246/SG0779 | this work |
| pAK2E185 | *bla lacA*::(P$_{xyl}$-*bceS*$^{C45S/C198S/C259S}$-*His8*) *mls* | pAK2E184 | SG0245/SG0782 & SG0246/SG0781 | this work |
| pAK2E186 | *bla lacA*::(P$_{xyl}$-*bceS*$^{C45S/C198S/C259S/I50C}$-*His8*) *mls* | pAK2E185 | SG0245/SG0783 & SG0246/SG0727 | this work |
| pAK2E187 | *bla lacA*::(P$_{xyl}$-*bceS*$^{C45S/C198S/C259S/I51C}$-*His8*) *mls* | pAK2E185 | SG0245/SG0784 & SG0246/SG0729 | this work |
| pAK2E188 | *bla lacA*::(P$_{xyl}$-*bceS*$^{C45S/C198S/C259S/F52C}$-*His8*) *mls* | pAK2E185 | SG0245/SG0785 & SG0246/SG0731 | this work |
| pAK2E189 | *bla lacA*::(P$_{xyl}$-*bceS*$^{C45S/C198S/C259S/L53C}$-*His8*) *mls* | pAK2E185 | SG0245/SG0734 & SG0246/SG0733 | this work |
| pAK2E190 | *bla lacA*::(P$_{xyl}$-*bceS*$^{C45S/C198S/C259S/W54C}$-*His8*) *mls* | pAK2E185 | SG0245/SG0736 & SG0246/SG0735 | this work |
| pAK2E191 | *bla lacA*::(P$_{xyl}$-*bceS*$^{C45S/C198S/C259S/F55C}$-*His8*) *mls* | pAK2E185 | SG0245/SG0738 & SG0246/SG0737 | this work |
| pAK2E192 | *bla lacA*::(P$_{xyl}$-*bceS*$^{C45S/C198S/C259S/R56C}$-*His8*) *mls* | pAK2E185 | SG0245/SG0740 & SG0246/SG0739 | this work |
| pAK2E193 | *bla lacA*::(P$_{xyl}$-*bceS*$^{C45S/C198S/C259S/Y57C}$-*His8*) *mls* | pAK2E185 | SG0245/SG0742 & SG0246/SG0741 | this work |
| pAK2E194 | *bla lacA*::(P$_{xyl}$-*bceS*$^{C45S/C198S/C259S/R58C}$-*His8*) *mls* | pAK2E185 | SG0245/SG0744 & SG0246/SG0743 | this work |
| pAK2E195 | *bla lacA*::(P$_{xyl}$-*bceS*$^{C45S/C198S/C259S/K59C}$-*His8*) *mls* | pAK2E185 | SG0245/SG0746 & SG0246/SG0745 | this work |
| pAK2E196 | *bla lacA*::(P$_{xyl}$-*bceS*$^{C45S/C198S/C259S/T61C}$-*His8*) *mls* | pAK2E185 | SG0245/SG0264 & SG0246/SG0263 | this work |
| pAK2E197 | *bla lacA*::(P$_{xyl}$-*bceS*$^{C45S/C198S/C259S/A62C}$-*His8*) *mls* | pAK2E185 | SG0245/SG0266 & SG0246/SG0265 | this work |
| pAK2E198 | *bla lacA*::(P$_{xyl}$-*bceS*$^{C45S/C198S/C259S/Y64C}$-*His8*) *mls* | pAK2E185 | SG0245/SG0403 & SG0246/SG0402 | this work |
| pAK2E199 | *bla lacA*::(P$_{xyl}$-*bceS*$^{C45S/C198S/C259S/K65C}$-*His8*) *mls* | pAK2E185 | SG0245/SG0405 & SG0246/SG0404 | this work |
| pAK2E200 | *bla lacA*::(P$_{xyl}$-*bceS*$^{E115K/K116E/C45S}$-*His8*) *mls* | pAK2E34 | SG0245/SG0829 & SG0246/SG0828 | this work |
| pAK2E201 | *bla lacA*::(P$_{xyl}$-*bceS*$^{E115K/K116E/C45S/C198S}$-*His8*) *mls* | pAK2E200 | SG0245/SG0780 & SG0246/SG0779 | this work |
| pAK2E202 | *bla lacA*::(P$_{xyl}$-*bceS*$^{E115K/K116E/C45S/C198S/C259S}$-*His8*) *mls* | pAK2E201 | SG0245/SG0782 & SG0246/SG0781 | this work |
| pAK2E203 | *bla lacA*::(P$_{xyl}$-*bceS*$^{E115K/K116E/C45S/C198S/C259S/I50C}$-*His8*) *mls* | pAK2E202 | SG0245/SG0783 & SG0246/SG727 | this work |
| pAK2E204 | *bla lacA*::(P$_{xyl}$-*bceS*$^{E115K/K116E/C45S/C198S/C259S/I51C}$-*His8*) *mls* | pAK2E202 | SG0245/SG0784 & SG0246/SG0729 | this work |
| pAK2E205 | *bla lacA*::(P$_{xyl}$-*bceS*$^{E115K/K116E/C45S/C198S/C259S/F52C}$-*His8*) *mls* | pAK2E202 | SG0245/SG0785 & SG0246/SG0731 | this work |

| | | | | |
|---|---|---|---|---|
| pAK2E206 | *bla lacA*::($P_{xyl}$-*bceS*$^{E115K/K116E/C45S/C198S/C259S/L53C}$-*His8*) *mls* | pAK2E202 | SG0245/SG0734 & SG0246/SG0733 | this work |
| pAK2E207 | *bla lacA*::($P_{xyl}$-*bceS*$^{E115K/K116E/C45S/C198S/C259S/W54C}$-*His8*) *mls* | pAK2E202 | SG0245/SG0736 & SG0246/SG0735 | this work |
| pAK2E208 | *bla lacA*::($P_{xyl}$-*bceS*$^{E115K/K116E/C45S/C198S/C259S/F55C}$-*His8*) *mls* | pAK2E202 | SG0245/SG0738 & SG0246/SG0737 | this work |
| pAK2E209 | *bla lacA*::($P_{xyl}$-*bceS*$^{E115K/K116E/C45S/C198S/C259S/R56C}$-*His8*) *mls* | pAK2E202 | SG0245/SG0740 & SG0246/SG0739 | this work |
| pAK2E210 | *bla lacA*::($P_{xyl}$-*bceS*$^{E115K/K116E/C45S/C198S/C259S/Y57C}$-*His8*) *mls* | pAK2E202 | SG0245/SG0742 & SG0246/SG0741 | this work |
| pAK2E211 | *bla lacA*::($P_{xyl}$-*bceS*$^{E115K/K116E/C45S/C198S/C259S/R58C}$-*His8*) *mls* | pAK2E202 | SG0245/SG0744 & SG0246/SG0743 | this work |
| pAK2E212 | *bla lacA*::($P_{xyl}$-*bceS*$^{E115K/K116E/C45S/C198S/C259S/K59C}$-*His8*) *mls* | pAK2E202 | SG0245/SG0746 & SG0246/SG0745 | this work |
| pAK2E213 | *bla lacA*::($P_{xyl}$-*bceS*$^{E115K/K116E/C45S/C198S/C259S/T61C}$-*His8*) *mls* | pAK2E202 | SG0245/SG0264 & SG0246/SG0263 | this work |
| pAK2E214 | *bla lacA*::($P_{xyl}$-*bceS*$^{E115K/K116E/C45S/C198S/C259S/A62C}$-*His8*) *mls* | pAK2E202 | SG0245/SG0266 & SG0246/SG0265 | this work |
| pAK2E215 | *bla lacA*::($P_{xyl}$-*bceS*$^{E115K/K116E/C45S/C198S/C259S/Y64C}$-*His8*) *mls* | pAK2E202 | SG0245/SG0403 & SG0246/SG0402 | this work |
| pAK2E216 | *bla lacA*::($P_{xyl}$-*bceS*$^{E115K/K116E/C45S/C198S/C259S/K65C}$-*His8*) *mls* | pAK2E202 | SG0245/SG0405 & SG0246/SG0404 | this work |
| pAKXT03 | *bla thrC*::($P_{xyl}$-*bceS*$^{WT}$) *spec* | pXT | SG0671/SG0672 (pAS719) | this work |
| pAKXT06 | *bla thrC*::($P_{xyl}$-*bceS*$^{E115K}$) *spec* | pXT | SG0671/SG0672 (PCF705) | this work |
| pAKXT13 | *bla thrC*::($P_{xyl}$-*bceS*$^{E115K/K116E}$) *spec* | pXT | SG0671/SG0807 & SG0672/SG0806 (pAS719) | this work |
| pAKXT15 | *bla thrC*::($P_{xyl}$-*bceS*$^{W54R}$) *spec* | pAKXT03 | SG0671/SG0813 & SG0672/SG0812 (pAKXT03) | this work |
| pAKXT16 | *bla thrC*::($P_{xyl}$-*bceS*$^{W54R/F55R}$) *spec* | pAKXT03 | SG0671/SG0817 & SG0672/SG0816 (pAKXT03) | this work |
| pAKXT20 | *bla thrC*::($P_{xyl}$-*bceS*$^{Y57R}$) *spec* | pAKXT03 | SG0671/SG0898 & SG0672/SG0897 (pAKXT03) | this work |
| pAKXT21 | *bla thrC*::($P_{xyl}$-*bceS*$^{F55R/Y57R}$) *spec* | pAKXT03 | SG0671/SG0900 & SG0672/SG0899 (pAKXT03) | this work |
| pAKXT22 | *bla thrC*::($P_{xyl}$-*bceS*$^{F55R}$) *spec* | pAKXT03 | SG0671/SG0815 & SG0672/SG0814 (pAKXT03) | this work |
| pAM703 | *bla thrC*::($P_{xyl}$-bceAB$^{L546C}$-Flag$_3$) *spec* | pFK727 | SG0117/SG0118 (QC) | this work |
| pAS719 | *bla thrC*::($P_{xyl}$-*bceS*$^{WT}$) *spec* | pXT | | gift from Mascher lab |
| pCF705 | *bla thrC*::($P_{xyl}$-*bceS*$^{E115K}$) *spec* | pXT | | Laboratory stock |
| pER603 | *bla amyE::$P_{bceA}$-lacZ cm* | | | (Rietkötter *et al.*, 2008) |
| pFK727 | *bla thrC*::($P_{xyl}$-bceAB$^{WT}$-Flag$_3$) *spec* | pXT | | (Kallenberg *et al.*, 2013) |
| pJL705 | *bla thrC*::($P_{xyl}$-bceAB$^{D556C}$-Flag$_3$) *spec* | pFK727 | SG0109/SG0110 (QC) | this work |

| | | | | |
|---|---|---|---|---|
| pMAD | *bla bgaB erm* | | | (Arnaud *et al.*, 2004) |
| pMG2E01 | *bla lacA*::($P_{xyl}$-*bceS*$^{L53C}$-*His8*) *mls* | pBS2E01 | SG0247/SG0248 (QC) | this work |
| pMG2E02 | *bla lacA*::($P_{xyl}$-*bceS*$^{W54C}$-*His8*) *mls* | pBS2E01 | SG0245/SG0250 & SG0246/SG0249 | this work |
| pMG2E03 | *bla lacA*::($P_{xyl}$-*bceS*$^{F55C}$-*His8*) *mls* | pBS2E01 | SG0251/ SG0252 (QC) | this work |
| pMG2E04 | *bla lacA*::($P_{xyl}$-*bceS*$^{R56C}$-*His8*) *mls* | pBS2E01 | SG0253/ SG0254 (QC) | this work |
| pMG2E05 | *bla lacA*::($P_{xyl}$-*bceS*$^{Y57C}$-*His8*) *mls* | pBS2E01 | SG0255/ SG0256 (QC) | this work |
| pMG2E06 | *bla lacA*::($P_{xyl}$-*bceS*$^{R58C}$-*His8*) *mls* | pBS2E01 | SG0257/ SG0258 (QC) | this work |
| pMG2E07 | *bla lacA*::($P_{xyl}$-*bceS*$^{K59C}$-*His8*) *mls* | pBS2E01 | SG0259/ SG0260 (QC) | this work |
| pMG2E08 | *bla lacA*::($P_{xyl}$-*bceS*$^{E60C}$-*His8*) *mls* | pBS2E01 | SG0261/SG0262 (QC) | this work |
| pMG2E09 | *bla lacA*::($P_{xyl}$-*bceS*$^{T61C}$-*His8*) *mls* | pBS2E01 | SG0263/SG0264 (QC) | this work |
| pMG2E10 | *bla lacA*::($P_{xyl}$-*bceS*$^{A62C}$-*His8*) *mls* | pBS2E01 | SG0265/SG0266 (QC) | this work |
| pMG2E12 | *bla lacA*::($P_{xyl}$-*bceS*$^{E60C/E115K}$-*His8*) *mls* | pBS2E01 | SG0261/SG0262 (QC) | this work |
| pMG2E15 | *bla lacA*::($P_{xyl}$-*bceS*$^{F63C/E115K}$-*His8*) *mls* | pBS2E01 | SG0400/SG0401 (QC) | this work |
| pMG704 | *bla thrC*::($P_{xyl}$-bceAB$^{Y547C}$-Flag$_3$) *spec* | pFK727 | SG0175/SG0176 (QC) | this work |
| pMG707 | *bla thrC*::($P_{xyl}$-bceAB$^{E553C}$-Flag$_3$) *spec* | pFK727 | SG0119/SG0120 (QC) | this work |
| pMG712 | *bla thrC*::($P_{xyl}$-bceAB$^{F538C}$-Flag$_3$) *spec* | pFK727 | SG0197/SG0198 (QC) | this work |
| pMG713 | *bla thrC*::($P_{xyl}$-bceAB$^{K549C}$-Flag$_3$) *spec* | pFK727 | SG0068/SG0069 (QC) | this work |
| pMG718 | *bla thrC*::($P_{xyl}$-bceAB$^{G543C}$-Flag$_3$) *spec* | pFK727 | SG0231/SG0232 (QC) | this work |
| pMG724 | *bla thrC*::($P_{xyl}$-bceAB$^{C544S}$-Flag$_3$) *spec* | pFK727 | SG0243/SG0244 (QC) | this work |
| pMG719 | *bla thrC*::($P_{xyl}$-bceAB$^{I545C}$-Flag$_3$) *spec* | pFK727 | SG0233/SG0234 (QC) | this work |
| pMG720 | *bla thrC*::($P_{xyl}$-bceAB$^{F548C}$-Flag$_3$) *spec* | pFK727 | SG0235/SG0236 (QC) | this work |
| pMG721 | *bla thrC*::($P_{xyl}$-bceAB$^{Q550C}$-Flag$_3$) *spec* | pFK727 | SG0237/SG0238 (QC) | this work |
| pMG722 | *bla thrC*::($P_{xyl}$-bceAB$^{M551C}$-Flag$_3$) *spec* | pFK727 | SG0239/SG0240 (QC) | this work |
| pMG723 | *bla thrC*::($P_{xyl}$-bceAB$^{G552C}$-Flag$_3$) *spec* | pFK727 | SG0241/SG0242 (QC) | this work |
| pMG738 | *bla thrC*::($P_{xyl}$-bceAB$^{S554C}$-Flag$_3$) *spec* | pFK727 | SG0356/SG0357 (QC) | this work |
| pMG739 | *bla thrC*::($P_{xyl}$-bceAB$^{E555C}$-Flag$_3$) *spec* | pFK727 | SG0358/SG0359 (QC) | this work |
| pMG759 | *bla thrC*::($P_{xyl}$-bceAB$^{T541C}$-Flag$_3$) *spec* | pFK727 | SG0624/SG0625 (QC) | this work |
| pMG760 | *bla thrC*::($P_{xyl}$-bceAB$^{S542C}$-Flag$_3$) *spec* | pFK727 | SG0622/SG0623 (QC) | this work |
| pMG762 | *bla thrC*::($P_{xyl}$-bceAB$^{L546A}$-Flag$_3$) *spec* | pFK727 | SG0614/SG0615 (QC) | this work |
| pMG763 | *bla thrC*::($P_{xyl}$-bceAB$^{L546F}$-Flag$_3$) *spec* | pFK727 | SG0616/SG0617 (QC) | this work |
| pMG764 | *bla thrC*::($P_{xyl}$-bceAB$^{Q550A}$-Flag$_3$) *spec* | pFK727 | SG0618/SG0619 (QC) | this work |
| pMG765 | *bla thrC*::($P_{xyl}$-bceAB$^{Q550F}$-Flag$_3$) *spec* | pFK727 | SG0620/SG0621 (QC) | this work |
| pMG770 | *bla thrC*::($P_{xyl}$-bceAB$^{L546N}$-Flag$_3$) *spec* | pFK727 | SG0675/SG0676 (QC) | this work |
| pMG771 | *bla thrC*::($P_{xyl}$-bceAB$^{Q550E}$-Flag$_3$) *spec* | pFK727 | SG0677/SG0678 (QC) | this work |
| pMG772 | *bla thrC*::($P_{xyl}$-bceAB$^{L546D}$-Flag$_3$) *spec* | pFK727 | SG0715/SG0716 (QC) | this work |
| pMG775 | *bla thrC*::($P_{xyl}$-bceAB$^{F533C}$-Flag$_3$) *spec* | pFK727 | SG0932/SG0933 (QC) | this work |
| pMG776 | *bla thrC*::($P_{xyl}$-bceAB$^{L534C}$-Flag$_3$) *spec* | pFK727 | SG0934/SG0935 (QC) | this work |
| pMG777 | *bla thrC*::($P_{xyl}$-bceAB$^{G535C}$-Flag$_3$) *spec* | pFK727 | SG0936/SG0937 (QC) | this work |
| pMG778 | *bla thrC*::($P_{xyl}$-bceAB$^{L536C}$-Flag$_3$) *spec* | pFK727 | SG0938/SG0939 (QC) | this work |
| pMG779 | *bla thrC*::($P_{xyl}$-bceAB$^{T537C}$-Flag$_3$) *spec* | pFK727 | SG0940/SG0941 (QC) | this work |
| pMG780 | *bla thrC*::($P_{xyl}$-bceAB$^{L539C}$-Flag$_3$) *spec* | pFK727 | SG0942/SG0943 (QC) | this work |
| pMG781 | *bla thrC*::($P_{xyl}$-bceAB$^{I540C}$-Flag$_3$) *spec* | pFK727 | SG0944/SG0945 (QC) | this work |
| pSD2E01 | *bla lacA*::($P_{xyl}$-*bceS*$^{WT}$-*His8*) *mls* | | | (Dintner *et al.*, 2014) |

| | | | | |
|---|---|---|---|---|
| pSDlux101 | pAH328-P$_{bceA}$-*luxABCDE* | | | (Kallenberg *et al.*, 2013) |
| pXT | *bla thrC*::(P$_{xyl}$) *spec* | | | (Derré *et al.*, 2000) |

[a] *E. coli* and *B. subtilis* antibiotic resistance markers are listed first and second respectively. *bla*, ampicillin resistance; cm, chloramphenicol resistance. *mls*, macrolide-lincosamide-streptogramin resistance; *spec*, spectinomycin resistance.
[b] Plasmid mutagenesis were performed by the PCR overlap extension method (Ho *et al.*, 1989) where two primer pairs are given, or by QuickChange (QC) with a single primer pair where indicated.
[c] DNA template used for mutagenesis was plasmid pSD2E01 unless stated otherwise in parentheses.

**References**


Arnaud, M., Chastanet, A., and Débarbouillé, M. (2004) New vector for efficient allelic replacement in naturally nontransformable, low-GC-content, Gram-positive bacteria. *Appl Environ Microbiol* **70**: 6887–6891.

Derré, I., Rapoport, G., and Msadek, T. (2000) The CtsR regulator of stress response is active as a dimer and specifically degraded *in vivo* at 37 degrees C. *Mol Microbiol* **38**: 335–347.

Dintner, S., Heermann, R., Fang, C., Jung, K., and Gebhard, S. (2014) A sensory complex consisting of an ATP-binding cassette transporter and a two-component regulatory system controls bacitracin resistance in *Bacillus subtilis*. *J Biol Chem* **289**: 27899–27910.

Ho, S.N., Hunt, H.D., Horton, R.M., Pullen, J.K., and Pease, L.R. (1989) Site-directed mutagenesis by overlap extension using the polymerase chain reaction. *Gene* **77**: 51–59.

Kallenberg, F., Dintner, S., Schmitz, R., and Gebhard, S. (2013) Identification of regions important for resistance and signalling within the antimicrobial peptide transporter BceAB of *Bacillus subtilis*. *J Bacteriol* **195**: 3287–3297.

Rietkötter, E., Hoyer, D., and Mascher, T. (2008) Bacitracin sensing in *Bacillus subtilis*. *Mol Microbiol* **68**: 768–785.

**Table S2. Minimal inhibitory concentration (MIC) of bacitracin in strains carrying BceAB variants.**

| Amino acid exchange in BceB | MIC of bacitracin (µg $ml^{-1}$)[a] |
|---|---|
| F533C | 16 |
| L534C | 32 |
| G535C | 4 |
| L536C | 32 |
| T537C | 64 |
| F538C | 32 |
| L539C | 32 |
| I540C | 32 |
| T541C | 32 |
| S542C | 32-64 |
| G543C | 16 |
| C544S | 16-32 |
| I545C | 16-32 |
| L546C | 32-64 |
| L546A | 32-64 |
| L546F | 64 |
| L546D | 16-32 |
| L546N | 32 |
| Y547C | 16 |
| F548C | 16 |
| K549C | 16-32 |
| Q550C | 32-64 |
| Q550A | 32-64 |
| Q550F | 64 |
| Q550E | 32-64 |

[a] MICs given as a range of values showed variable results between biological triplicates; the MIC for strains carrying the wild-type construct was 32-64 µg $ml^{-1}$; the MIC of strains lacking *bceAB* was 4 µg $ml^{-1}$.

**Table S3. Parameters used for GaMD simulations (timestep = 2 fs).**

| Parameter | Assignment |
|---|---|
| igamd | 3 (Boost on both dihedral and total potential energy). |
| ntcmdprep | 3000000 (Number of preparation MD steps). |
| ntcmd | 6000000 (Number of initial conventional MD steps to calculate system potential energies). |
| ntebprep | 3000000 (Number of preparation biasing MD steps). |
| nteb | 33000000 (Total number of biasing MD steps). |
| ntave | 300000 (Time step to calculate standard deviation of potential energies). |
| sigma0P | 6 (Upper limit of standard deviation of total potential boost). |
| sigma0D | 6 (Upper limit of standard deviation of dihedral potential boost). |
